# De novo design of allosterically switchable protein assemblies

**DOI:** 10.1101/2023.11.01.565167

**Authors:** Arvind Pillai, Abbas Idris, Annika Philomin, Connor Weidle, Rebecca Skotheim, Philip J. Y. Leung, Adam Broerman, Cullen Demakis, Andrew J. Borst, Florian Praetorius, David Baker

## Abstract

Allosteric modulation of protein function, wherein the binding of an effector to a protein triggers conformational changes at distant functional sites, plays a central role in the control of metabolism and cell signaling^1–3^. There has been considerable interest in designing allosteric systems, both to gain insight into the mechanisms underlying such “action at a distance” modulation and to create synthetic proteins whose functions can be regulated by effectors^4–7^. However, emulating the subtle conformational changes distributed across many residues, characteristic of natural allosteric proteins, is a significant challenge^8,9^. Here, inspired by the classic Monod-Changeux-Wyman model of cooperativity^10^, we investigate the de novo design of allostery through rigid-body coupling of designed effector-switchable hinge modules^11^ to protein interfaces^12^ that direct the formation of alternative oligomeric states. We find that this approach can be used to generate a wide variety of allosterically switchable systems, including cyclic rings that incorporate or eject subunits in response to effector binding and dihedral cages that undergo effector-induced disassembly. Size-exclusion chromatography, mass photometry^13^, and electron microscopy reveal that these designed allosteric protein assemblies closely resemble the design models in both the presence and absence of effectors and can have ligand-binding cooperativity comparable to classic natural systems such as hemoglobin^14^. Our results indicate that allostery can arise from global coupling of the energetics of protein substructures without optimized sidechain-sidechain allosteric communication pathways and provide a roadmap for generating allosterically triggerable delivery systems, protein nanomachines, and cellular feedback control circuitry.

## MAIN TEXT

Cellular control of signaling and metabolic pathways requires context-dependent modulation of protein function. This modulation is primarily achieved through allosteric regulation, where a regulatory “effector” molecule binds to a specific site on a protein and alters the structure and function at a distant active site or binding interface via long-range conformational coupling^3,10,15,16^. A particularly important case of allostery involves coupling between subunits in oligomeric complexes, resulting in the synchronization of global conformational transitions^10,17^. In proteins such as hemoglobin^10,14,17^ and aspartate transcarbamoylase^18^, binding of a ligand to one subunit enhances the binding affinity of the other subunits in the complex, resulting in a sharp, switch-like response to ligand across a narrow concentration range (i.e. cooperativity). Tight allosteric coupling between binding pockets and oligomeric interfaces also links the rotor motions of ATP-synthases to ATP-formation^19^, enables GROES/GROEL to cycle between loading and releasing its protein cargo^20^, and underpins the regulation of numerous enzyme functions via selective stabilization of distinct oligomeric forms that differ in activity^21^.

Designing synthetic complexes that can allosterically toggle between distinct oligomeric forms is an important goal in protein engineering as it could enable the construction of switchable nanomaterials, cooperatively activated biosensors, and molecular machines that perform work via coordinated movements among their components. Previous work on the de novo design of allostery has demonstrated coupling between two ligand binding events by incorporating both binding modules into a shared monomeric helical bundle architecture^5^, and between metal binding and disruption of a spatially distant disulfide bridge in an oligomeric interface^22^. However, a general approach for designing allosterically modulable oligomeric assemblies which couple an internal binding event to a global change in quaternary structure, as in hemoglobin, has not been described.

We set out to design allosterically coupled oligomeric assemblies taking inspiration from the classic Monod-Wyman-Changeux (MWC) model of allostery^10^, which models each subunit as having two states with different affinities for ligand, and a strong preference for oligomeric assemblies with all subunits in the same state (**Fig 1a**). The transition between the two quaternary states is treated as a large-scale, rigid-body movement in which the subunits change orientation relative to one another^10^. This is a considerable simplification of actual allosteric communication between sites or subunits in native proteins, which typically involves subtle coordinated structural changes at dozens of residues^8,23^, and many intermediary conformational states may be occupied^24–26^. The intricate nature of these conformational shifts can obscure the underlying biophysical mechanisms, making it difficult to modify or reverse engineer these systems to perform new tasks. While the simplifications of the MWC model have the advantage of enabling quantitative modeling, they come with a potential loss of physical realism.

**Figure 1.**
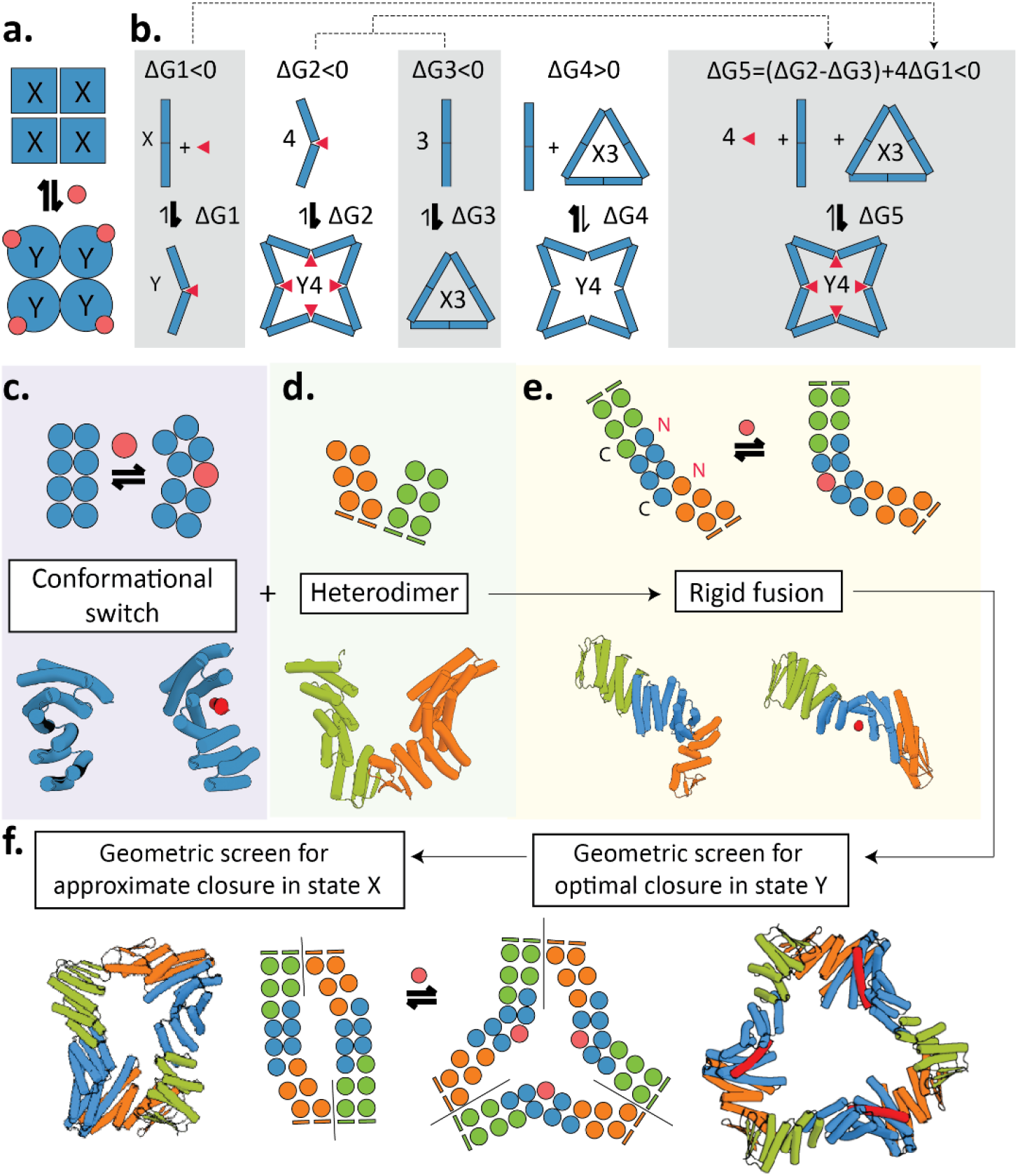
Design strategy for building switchable oligomers. **A.** The two-state nature of the Monod-Changeux-Wyman model in the case of tetrameric Hemoglobin, which involves an equilibrium between two defined oligomeric states in which the monomers must all adopt either the “X” (blue squares) or “Y” (blue circle) conformation. Cooperativity arises from the fact that the two quaternary conformations differ in their affinity for a ligand (red circle). **B.** Equilibria relevant for the construction of an oligomeric switch. ΔG1 refers to the energy difference between the hinge in the apo X conformation and the bound Y conformation. ΔG2 is the energy for the assembly of n Y-state monomers into a Yn ring in the presence of the ligand. ΔG3 is the energy for the assembly of m X-state monomers into a Xm ring in the absence of ligand. ΔG4 is the energy differentiating the Xm ring from the Yn ring in the absence of peptide and must be positive for the switch to operate. ΔG5 is a linear function of these energies, and must be negative for the switch to operate. **C-E.** Schematic 2D representation (top, circles represent individual helices) and exemplary protein structure (bottom, cylinders represent individual helices) illustrating the building blocks and fusion based design approach. **C.** “Hinge” building block (blue) switches between conformations X (left) and Y (right). Binding of an effector peptide (red) stabilizes conformation Y. **D.** Reversible heterodimer consisting of two individually stable and soluble monomers (green and orange). **E.** Example chimeric protein resulting from fusion of components shown in C. and D. shown in state X (left) and state Y (right). **F.** Oligomers formed by the fusion proteins shown in E. with state X (left) adopting a different oligomeric state than state Y (right). Colors and representations as in C-E.

We reasoned that for the de novo design of allosterically coupled assemblies, the reductive nature of the MWC model could be an advantage: rather than having to model the complexities of sidechain-to-sidechain communication between distant sites, and the large number of corresponding micro-states, we could focus on designing systems built from monomers that can adopt just two distinct conformations with different affinities for a ligand, and that can assemble into different oligomeric conformations (**Fig 1b**). To facilitate experimental characterization, we chose to design proteins that can adopt two alternative oligomeric states with different numbers of subunits, rather than toggling between oligomeric conformations that contain the same number of monomers, as is typically envisioned in the MWC model (**Fig 1a**). This choice allows for easy differentiation of quaternary conformations using size exclusion chromatography (SEC), mass photometry^13^ (MP), and negative stain electron microscopy (nsEM) while preserving the underlying principles of the MWC model^10^. Experimental characterization of the designs could then reveal the extent of coupling between ligand binding and change of oligomeric state, and whether explicitly designing side-chain-based pathways of intramolecular communication is truly necessary for the design of allostery.

### Design of switchable ring-forming proteins

We began by designing proteins that can adopt two different oligomeric ring states that differ in their radius and number of subunits (**Fig 1b**). Within the monomeric subunits of these oligomers, we embed a two-state “hinge” module^27^ that can toggle between two structurally defined alternative conformations, a closed “X” state, and an open “Y” state, the latter of which presents a groove that can bind to an effector peptide with nanomolar affinity^11^. In the absence of the effector, the “X” predominates, while the “Y” dominates in the presence of saturating amounts of effector (**Fig 1c, EFig.1a,** ΔG1 in **Fig 1b**). We reasoned that rigidly fusing these hinge modules to protein interaction modules (**Fig 1d-f, EFig.1**) could enable the design of a wide array of cyclic assemblies whose oligomerization can be modulated by peptide binding, provided four constraints are satisfied: In monomers with the hinge in state Y, the interaction surfaces should direct the formation of an n-subunit cyclic oligomer (Yn; constraint 1; ΔG2 in **Fig 1b**), and, with the hinge in state X, the formation of a distinct m-subunit oligomer (Xm; constraint 2; ΔG3 in **Fig 1b**) where m and n are different numbers. Xm should be populated in the absence of effector (constraint 3, ΔG4 in **Fig 1b**), but the interfaces must be sufficiently strained that upon addition of effector the system transitions fully to state Yn, which is not populated in the absence of effector (constraint 4, ΔG5 in **Fig 1b**).

We developed a computational procedure for designing allosterically switchable systems satisfying the four constraints (**Fig c-f, EFig.1c**). As interaction surfaces we used previously designed and characterized heterodimeric interface modules which are sufficiently polar to reversibly assemble and disassemble without having to denature them (“LHDs”)^12^. To satisfy constraint 1, we used the WORMS software^28^ to fuse hinge modules in the Y state with LHD interaction modules such that n copies of the resulting Y state monomer close perfectly into an n subunit cyclic oligomer (**Fig1d-f**, **EFig.1c**). WORMS rapidly scans through hundreds of millions of possible fusions differing in subunit identity and fusion cut points for those satisfying specified geometric criteria^20^. We sequence designed the newly formed helical junctions generated by WORMS between the hinge and interaction modules with proteinMPNN^29^ to favor rigidity, solubility and stability of the fused monomers and selected designs which alphafold2 (AF2)^30^ predicted to fold to the target monomeric state (plod>0.85 and C-α rmsd <2.75Å, **EFig.1c**). To satisfy constraint 2, we used AF2 to predict state X conformations of our hinge fusions (in the absence of peptide) and used iterative alignment-based docking to generate cyclic oligomers with different copy numbers of X monomers. We then selected those designs for which one of these X-state oligomers come close to cyclic closure, allowing a slightly strained ground state assembly (**Fig1f, EFig.2**). We implemented this by sub-selecting designs for which a distinct number (between 2 and 5) of copies of the X state monomer was close to closing (<24Å between the N-terminal LHD domain of the first subunit and the C terminal LHD domain of the mth subunit, **EFig.2a**), but perfect ring closure (as modeled using AF2 multimer^31^) required significant bending by 1.5 Å-7Å relative to the ground state monomer conformation (**EFig.2c-e)**. Furthermore, if one LHD-interface remains unsatisfied in a partially open ring state, the close approach of the LHD interfaces should sterically occlude assembly into oligomers larger than Xm. (**EFig.6e-f**). We hypothesized that imperfect closure *in silico* in this state would yield strained rings that are sufficiently suboptimal to be broken by peptide-binding, but stable enough to assemble at relevant protein concentrations (**Fig1b**).

We used this approach to design proteins predicted to adopt distinct cyclic symmetries in the presence and absence of effector, utilizing as building blocks diverse sets of LHD interaction modules, hinge modules, and peptide effectors. 17 out of 26 tested rings, were soluble and monodisperse by size exclusion chromatography (SEC) (**EFig.3-5**). For 10 of the 17 soluble rings we observed a sizable shift in SEC retention volume of the predominant protein peak when 10 μM of peptide was added to 5 μM of hinge fusion, consistent with either increasing or decreasing size as expected from the design model (**Fig 2a,b,d, EFig.5)**.

**Figure 2.**
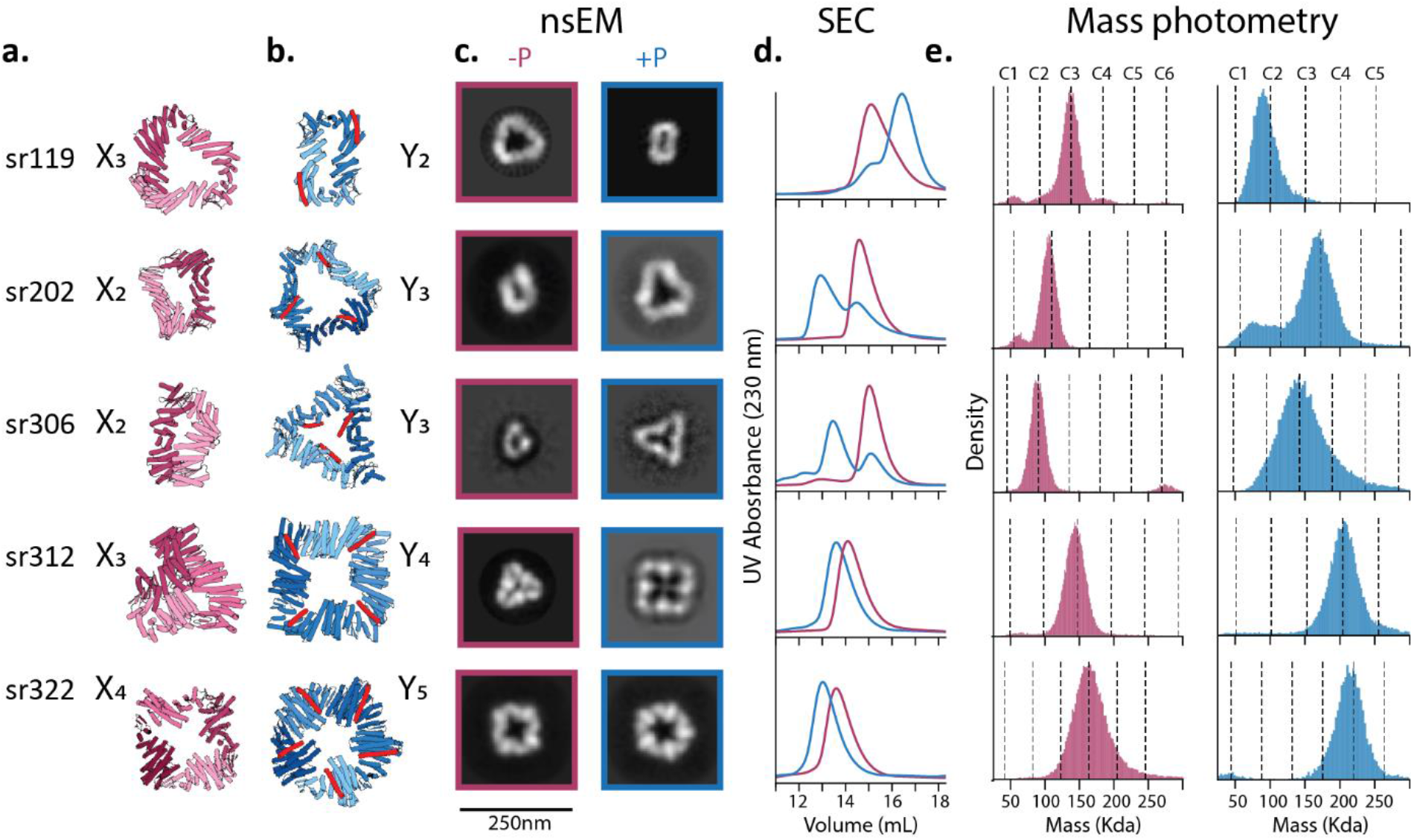
Design of allosterically controlled cyclic assemblies. **A.** Design models in their unbound Xm oligomeric state (pink). **B.** Design models in the peptide-bound Yn state assembly (blue: protein, red: effector peptide). **C.** nsEM 2D class averages for designs in the absence (left) and presence (right) of peptide representing top views of the predominant oligomeric species by particle count. **D.** Size exclusion chromatography on each design at 5 μM in the absence (pink) and presence (blue) of 10 μM peptide. **E.** Mass photometry on these designs at 100 nM final protein concentration in the absence (left) and presence (right) of 10-fold molar excess of peptide.

To precisely determine the oligomerization state of the switchable assemblies in the presence and absence of saturating amounts of effector, we used mass photometry. We measured the assembly state of 5 of the rings using MP^13^ in the presence and absence of 10-fold molar excess of effector. There was a clear shift in oligomeric state in all cases, with the predominant mass peaks consistent with expected masses of Xm and Yn within <±5% mass error (**Fig 2e, EFig.7**). The mass distributions in this nanomolar concentration regime show little to no detectable occupancy of alternative oligomeric forms. The affinity of the 5 designs for their peptide effector was quantified with fluorescence polarization, and their apparent dissociation constants ranged from low to high nanomolar affinity (**EFig.8**).

To characterize the structures of the candidate assemblies in the bound and unbound state, we visualized them with negative stain electron microscopy, both in the presence and absence of 20-fold molar excess of their effectors (**Fig 2c**). Sr119 was designed to adopt a roughly triangular C3 shape in the X3 state and collapse into a dimeric ring with a narrow aperture when the effector peptide (FF74B) binds to its outer rim (**Fig 2a**). 2D classes obtained from nsEM comparisons of the peptide-free and peptide-added samples show precisely this shape transition from a C3 to a C2, corroborating the data from MP (**Fig 2c, Efig 9**). Xm appears to be a closed ring which, according to the AF2 multimer prediction, requires a slight deformation of the X-state monomers (**EFig.6c,d**). sr202 and sr306 were designed to undergo the opposite oligomeric transition, growing in size from a compact dimer to a C3 symmetry, driven by two distinct effectors. In both cases a clear shift from a dimer assembly to a Y3 assembly is apparent, with the Y3 form closing into a triangular ring clearly visible in top-views and bottom-views obtained from nsEM (**Fig 2c, EFig.10-11**). The nsEM data for the X2 state shows that, rather than forming fully closed dimers, one of the interfaces in these dimers is not satisfied, as represented in a rigid dock of X-state monomers, resulting in a partially open assembly. However, the closeness of approach of the exposed LHD termini blocks assembly into higher multimers, including trimers or open-ended fibers (**EFig.6e,d**); both appear to be sterically prohibited as neither are observed by nsEM (**EFig.10-11**) or MP (**Fig 2e**).

Sr312 and sr322 were designed to switch between even higher-order ring states, from C3 to C4 and from C4 to C5, respectively. Top-views of Sr312 from nsEM show closure into a compact, triangular ring with a small pore (<20Å), in agreement with design expectations (**Fig 2c, EFig.12)**. Addition of the effector drives the system into a square-like C4, with a substantial increase in the size of interior pore (∼80Å diameter, **Fig 2c, EFig.12)**. Sr322, is an X4 tetramer in which the subunits close into a C4-symmetric ring. Peptide binding to the top side of the ring induces the subunits to twist and kink to accommodate one additional subunit, giving rise to a pentagonal Y5 ring which is the predominant species observed in nsEM grids of the peptide-treated samples (**Fig 2c, EFig.13**). For sr322, we also observed a minor population of hexagonal Yn+1 rings (<10% by particle count) in the presence of peptide (**EFig.12,13),** a likely byproduct of the same backbone flexibility that permits closure in the X state. These data demonstrate that our design approach can generate switchable oligomeric assemblies spanning a diversity of symmetries, radii and cognate peptides.

To better understand the role of protein flexibility and deformation in closure of our rings we used cryogenic electron microscopy (CryoEM) to obtain higher resolution structure for two different tetramers: sr312 in its peptide-bound Y4 state, and sr322 in its peptide-free (“apo”) X4 state (**Fig. 3**). The structure of sr312 (which was designed to form a perfect C4-symmetric ring in its peptide-bound state) exhibited high agreement with the computationally predicted design model, with the cryoEM structural data fully resolving the presence of the designed effector peptide. While a slight swelling of the cryoEM structure was noted compared to the designed oligomer model (tetramer RMSD = 5.14 Å), the agreement between monomer backbone and design was exceptionally high (monomer RMSD = 1.82 Å) (**Fig. 3a-d, Efig. 14**). For sr322 (which was designed to form a perfect C5-symmetric ring in its bound state), our iterative alignment-based docking protocol predicted a slightly open tetramer in the apo state (X4) (**Fig. 3f**). The cryo-EM structure of the sr322_apo reveals the formation of a perfectly closed C4 assembly in the X4 oligomer state at a final resolution of 4.62 Å (**Fig. 3g, Efig. 15-18,** see *Supplementary Note*). Comparisons between the structure of the monomer and an AF2-predicted X-state model show that this closure is made possible through deformation of the individual ring components (monomeric RMSD Cα = 3.03 Å) (**Fig. 3g, Efig. 17**). This bending away from the predicted conformation occurs in both the N-terminal LHD-domain and the hinge region, the latter being a likely consequence of the flexibility inherent in its alpha-helical repeat architecture^32^.

**Figure 3.**
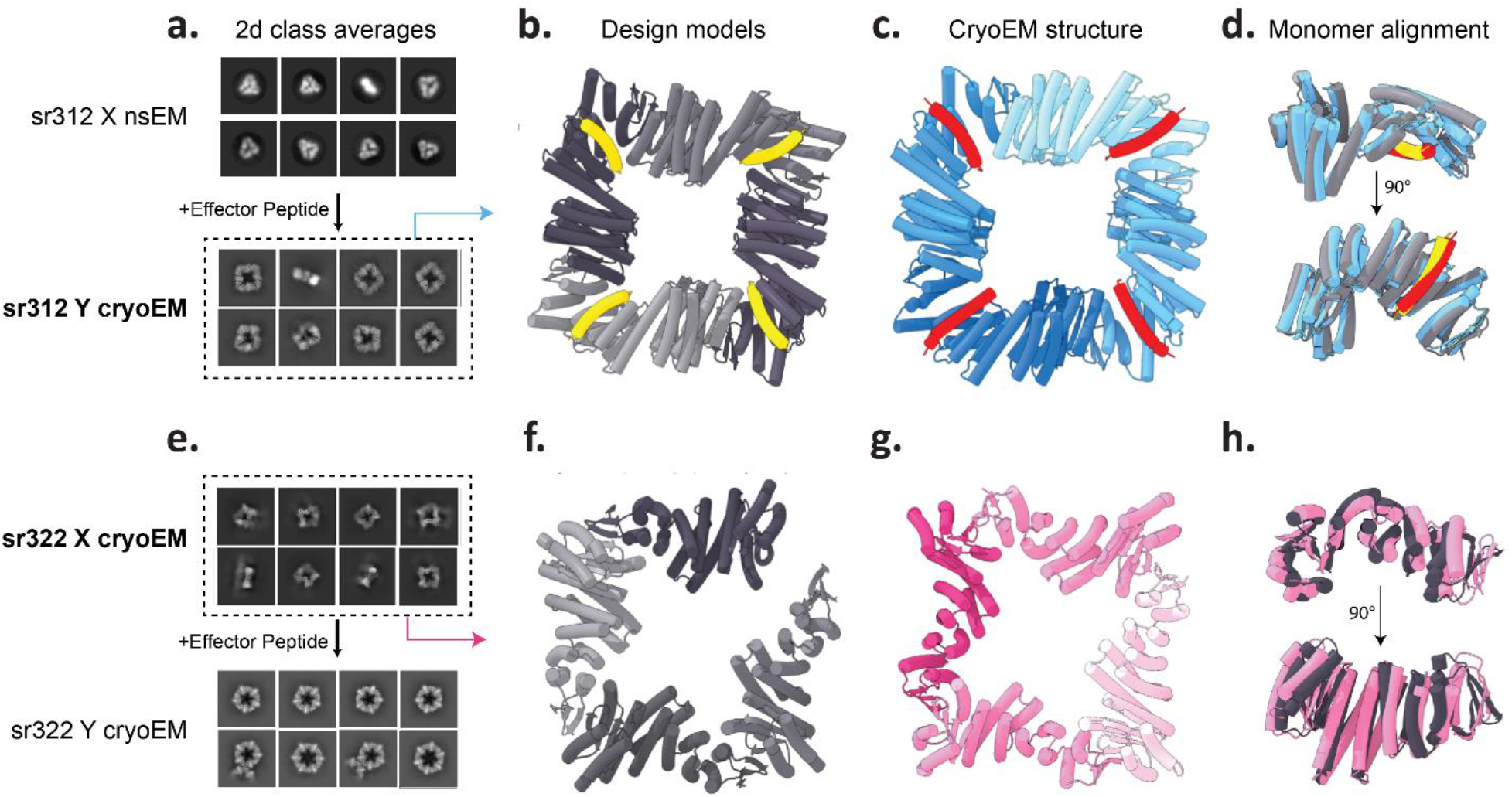
Structural characterization of sr312 and sr322 by electron microscopy. **A.** Comparison of 2D class averages of sr312 in the absence of peptide (top) from nsEM and presence of peptide (bottom) from cryoEM. **B.** Y-state C4 Design model for sr312 (gray) with effector peptide bound (yellow). **C.** CryoEM structure (resolution 4.40Å) of sr312 (blue) with effector peptide bound (red). **D.** Alignment of model and experimental structure for a single monomeric subunit of sr312. Grey/yellow: design model; blue/red: cryoEM structure; RMSD = 1.82 Å. **E.** Comparison of CryoEM 2D class averages of sr322 in the absence of peptide (top) and presence of peptide (bottom, See supplementary note 1 for details). **F.** X4 design model of sr322 (gray) generated by alignment-based docking **G.** CryoEM structure (pink, resolution 4.55Å) of sr322 without peptide. **H.** Alignment of model and experimental structure for a single monomeric subunit of sr322. Grey: design model; pink: cryoEM structure; RMSD = 3.03 Å.

To test the necessity of imperfect closure and strain in the X state for switching of our rings (constraint 4), we also designed “state X rings” in which the X state, in the absence of an effector, is expected to form a perfectly closed ring (see methods for details, **EFig.19**), while the Y-state oligomer would be slightly strained. 9 out of 24 tested state X rings were soluble and monodisperse by SEC, but no shifts in retention volume were observed upon addition of peptide (**EFig.19**). These data suggest that the peptide-dependent switching of our rings requires a strained or suboptimal ground state. To test the necessity of sterically occluding the LHD interfaces in the imperfectly closed state X assembly (**EFig.6e**), we also tested 10 designs that would leave the LHD interfaces accessible in state X, allowing for unbounded assembly formation (an example is shown in **Efig 2b**). As expected, all of these designs were insoluble.

### sr312 displays homotropic ligand binding cooperativity

We hypothesized that the switchable rings would bind to the effector peptide in a cooperative manner because binding events to the ground state Xm ring are energetically penalized relative to the Yn ring (**Fig 4a,f**). The first binding event could result either in an asymmetric assembly with one Y-state hinge and m-1 X-state hinges, or a closed symmetric structure with n-1 unbound Y state proteins, neither of which are expected to be more stable than the ground-state Xm ring because they expose unsatisfied peptide or LHD binding interfaces. In contrast, the final binding event involves the closure and complete ligation of an optimal peptide-binding Yn ring. We reasoned that this non-equivalence and non-independence of sequential binding events should manifest as MWC-like binding cooperativity^10^. The TRAP protein, where partial ligand saturation of a Trp-binding oligomeric ring results in an ensemble that is dominated by unbound ring and fully bound ring, with low occupancy of partially ligated species^33^, provides a close natural analog for our rings (**Fig 4a**). The large mass and radius shift of sr312 (**Fig2c, EFig.12**) upon peptide-binding make this design an ideal candidate for interrogating this question, because mixed populations of X3 and Y4 ring states can be readily distinguished by SEC, MP and nsEM.

**Figure 4.**
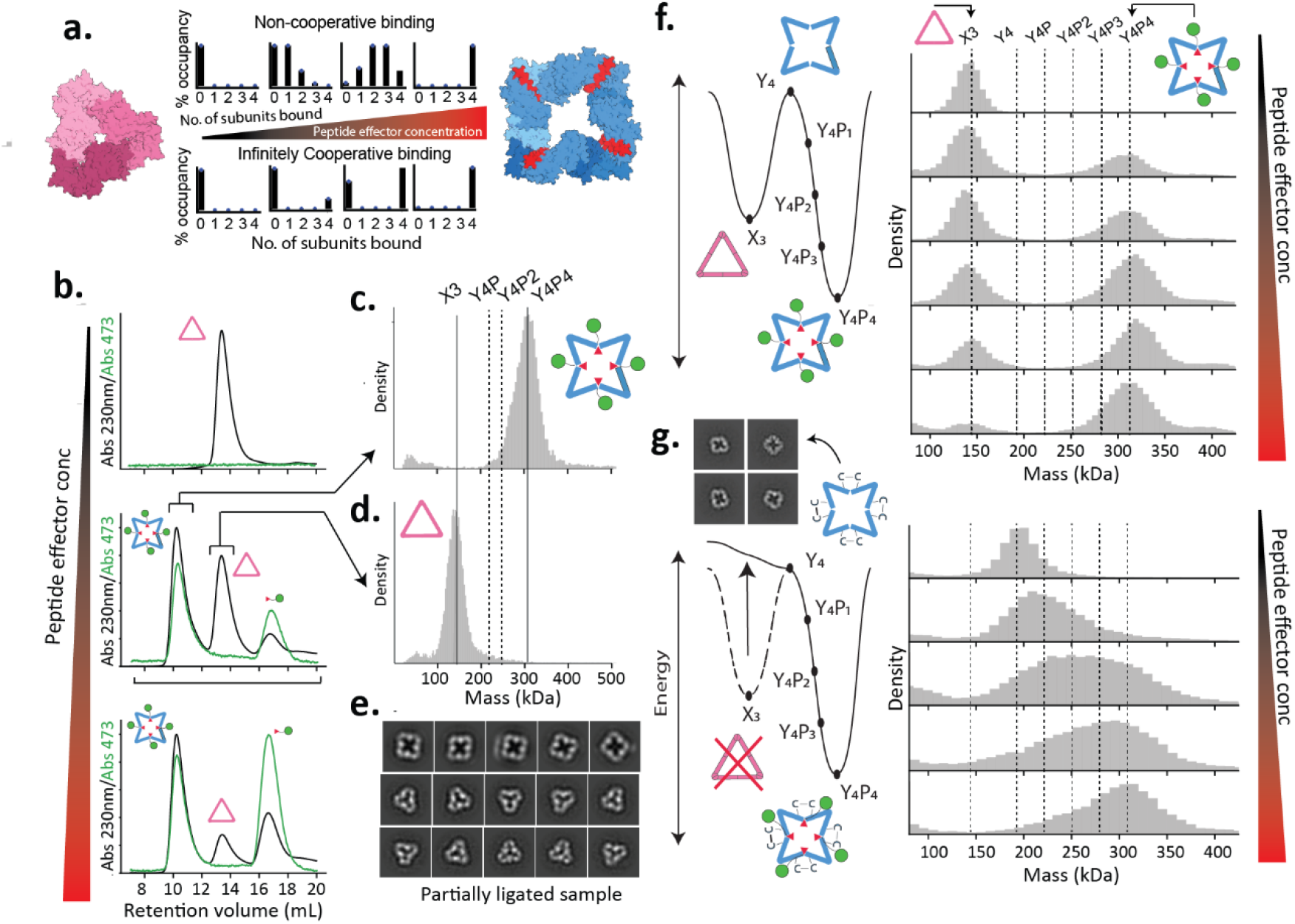
Allosteric state switching is cooperative. **A.** Theoretical distributions of peptide-bound ring states (i.e. rings bound to 0,1,2, 3 or 4 peptides) for a tetrameric ring across a peptide-titration series, from 0% to 100% fractional saturation of binding sites for a non-cooperative binder (top) and an infinitely cooperative binder (bottom). Left and right, modeled structures of sr312 in the Xm and Yn states. **B.** Size exclusion chromatography of 5 μM sr312 in absence of peptide (top), with 5 μM peptide (middle), and with 10 μM peptide (bottom). Protein (230 nm) and GFP absorbance (473 nm) are shown in black and green respectively. **C, D.** Mass photometry on fractions corresponding to earlier and later peaks in B., middle subpanel. Icon shows oligomeric species that matches the estimation of mass. Vertical lines indicate expected masses of fully bound, unbound, and partially ligated species. **E.** 15 most populated 2D nsEM classes from nsEM on the mixture shown in B. **F.** Characterization of sr312’s cooperativity. Left, a representation of the presumptive energy landscape of the apo (pink) and bound (blue) states of the X3 to Y4P4 ring switch. “P” refers to the number of peptides bound to the ring. Right, MP on samples of 3 μM sr312 incubated with 0 μM, 2 μM, 3 μM, 5 μM, 7 μM, 8 μM GFP-tagged peptide. Vertical lines represent expected masses of GFP-bound and apo species. **G.** Presumptive energy landscape for a variant of sr312 that is locked into the Y state via a disulfide bridge. Inset: nsEM class averages of this apo-state tetramer. Right, MP on samples of 3 μM of sr312_locked incubated with 0 μM, 1 μM, 2.5 μM,4 μM and 5 μM GFP-tagged peptide.

To test ligand-binding cooperativity, we measured the population distribution of oligomeric species at different peptide concentrations using SEC and MP. The ∼30kDa GFP-tagged peptide allows for sufficient mass resolution to distinguish between different occupancies in SEC and MP without introducing overlapping background signals. In a perfectly cooperative system, at ∼50% saturation, we would expect a bimodal distribution with two cleanly separated peaks, one of which corresponds to the trimeric, unliganded X3 ring and the other corresponding to a tetrameric Y4 ring with four ligands bound (**Fig 4a**). Meanwhile, in a non-cooperative system, there would be a binomial distribution of bound states that shifts to higher mean occupancies as the concentration of ligand is increased (**Fig 4a**). At 5 µM sr312 (monomer concentration) and 5 µM peptide, we observe SEC traces featuring two well-resolved peaks with baseline separation (**Fig 4b, EFig.20a**). Peak integration indicates that, under these conditions, 48% of the proteins are in the tetrameric state and the remaining 52% are in the trimeric state. To determine the ligation state of these two peaks, we isolated the relevant fractions and measured their mass with MP. The earlier elution peak corresponded to the mass of a Y4 ring bound to 4 peptides, denoted Y4P4, (**Fig 4c**), while the later peak corresponded to the mass of an X3 ring with no peptide, denoted X3P0, with little to no signal for other mass species (**Fig 4d**), consistent with the cooperativity hypothesis. To structurally characterize the ring-states present within this mixture, we imaged the partially ligated sample with nsEM. 2D class averages revealed that Y4 and X3 rings identical in appearance to the fully apo and fully holo forms (**Fig2c**) are the predominant species (**Fig 4e**), with no obvious signature of intermediate mixed XY species (**EFig.21b**).

To map this cooperative transition with higher resolution, we measured the occupancy of the multiple possible bound ring states across a titration series using MP spanning a ligand concentration range from 2 to 8 µM. We found that at ∼50% ring saturation, the mass distribution is dominated by X3 (with no peptide bound) and Y4P4, with sparse to no occupancy of the Y4P2 form that would be expected to dominate under these conditions in a non-cooperative system (**Fig 4a, f**). More generally, across the titration regime we see a clear conversion from X3 to Y4P4, with only limited occupancy of partially liganded intermediates. (**Fig 4f**). Fitting of the fractional occupancies of the bound state hinge estimated from MP across the titration range indicates an apparent Hill coefficient of ∼2.9 (**EFig 20i**). In conclusion, the bimodal nature of the oligomer distribution across the ligand titration series strongly suggests that sr312 binds to its cognate peptide cooperatively.

To determine if the two-state, conformationally switchable nature of the ring is essential for binding cooperativity, we generated a static version of sr312 (sr312_locked) containing two cysteine mutations in the hinge region which, under oxidizing conditions, form a disulfide bridge that locks the hinge into the Y state^11^ and thus locks the ring in the Y4 state (**Fig 4g**). nsEM on this construct shows that, after oxidation, the protein assembles into C4-symmetric tetramers comparable to the peptide-bound Y4 state of sr312 even without peptide (**Fig 4g, EFig.20g**). Binding experiments in MP show that, rather than primarily populating only the Y4 and Y4P4 states in the presence of peptide, the Y-stapled rings span a broad distribution of oligomeric states, with significant relative occupancy of Y4P1 and Y4P2 at partial saturation. Eliminating the X3 state therefore appears to eliminate cooperativity. These data show that the conformationally switchable nature of the designed ring subunits is critical for cooperative binding to peptide (**Fig 4g**).

The ligand-binding process could pass through various possible, marginally populated intermediates, including either asymmetric oligomers (i.e. X2Y, XY2, **EFig.21a**), or partially ligated Y4s. Significant presence of the former would be consistent with the Koshland-Némethy-Filmer (KNF) sequential model of cooperativity^34^, where tertiary changes in individual subunits drive the transition, while the latter is consistent with the MWC model, where global changes in quaternary structure are dominant. These partially ligated ring species should be more abundant at lower fractional saturation of the hinge (<50%), as has been observed in other cooperative binding systems, like the TRAP complex^33^. Both SEC and MP show evidence of these intermediate species (**EFig.20a,h**) at ∼10% saturation, but the low signal from these populations, and the resolution limits of these instruments preclude reliable assignment of the species present. We conducted two experiments at low fractional saturation to explore which of these intermediates are actually populated. First, we conducted a SEC binding titration series with fluorescently labeled peptide, to monitor binding of the peptide to possible trimeric intermediates and to determine the stoichiometry of binding to the tetramer (by comparing the ratio of A230 protein signal to TAMRA emission signal). We found that the peptide co-elutes as a single peak with the tetramer across this titration range (from 100 nM to 10 μM of peptide, against 1 μM of ring), implying that trimer-populations with bound peptide are negligible even at low (1:10 peptide:protein) substoichiometric ratios (**EFig.20b,e**). Additionally, the ratio of tetramer protein A230 to TAMRA fluorescence increases at lower peptide concentrations, suggesting the presence of tetramers that are only partially bound to peptide (**EFig.20f**). Next, we mixed peptide with protein at a concentration that yields small amounts of oligomers intermediate in size between Y4P4 and X3 that are detectable in both SEC and MP (3 μM protein:500nM peptide, ∼10% saturation by MP), isolated the bound fraction in SEC and examined these intermediate species with nsEM. In the nsEM 2D class averages, no significant subpopulations of broken or visibly asymmetric species were observed, and the particle counts are dominated by closed C4 oligomers (**EFig.21c**). MP on this isolated, partially ligated subpopulation clearly indicated a mass corresponding to a Y4 ring bound to two peptides, Y2P2 (**EFig. 20j**), a species we expect to observe under the MWC model. Although our experiments cannot absolutely rule out the existence of transiently populated bound, broken trimer states, they suggest a mechanism of cooperativity that primarily involves the preservation of symmetry during the binding process, directly analogous to the MWC model. The binding of one or two peptides enhances the stability of a transiently populated C4 oligomer, increasing the probability of unbound subunits being in the Y state, and thereby increasing the strength of subsequent binding events.

### Coupling peptide-binding to dimerization

In nature, allosteric modulation of oligomerization extends beyond toggling between alternative oligomer states to include cases where binding of an effector quantitatively influences the affinity of a dimeric protein-protein interface^35,36^. Given the many applications of ligand inducible dimerization systems in synthetic biology^37^, we sought to design an inducible homodimer where assembly across a C2 interface is allosterically enhanced by the addition of effector peptide. We used the WORMS based design procedure described above to generate LHD-Hinge fusions that yield a C2-symmetric homodimer assembly in the peptide-bound Y2 state. We computationally selected for and experimentally characterized designs in which the X-state monomers are sterically blocked from forming optimal interactions via their LHD binding surfaces (**Fig 5a,b**) to prevent formation of the dimer in the absence of effector. The addition of effector should relieve these steric clashes, allowing for the monomers to assemble into Y-state dimers.

**Figure 5.**
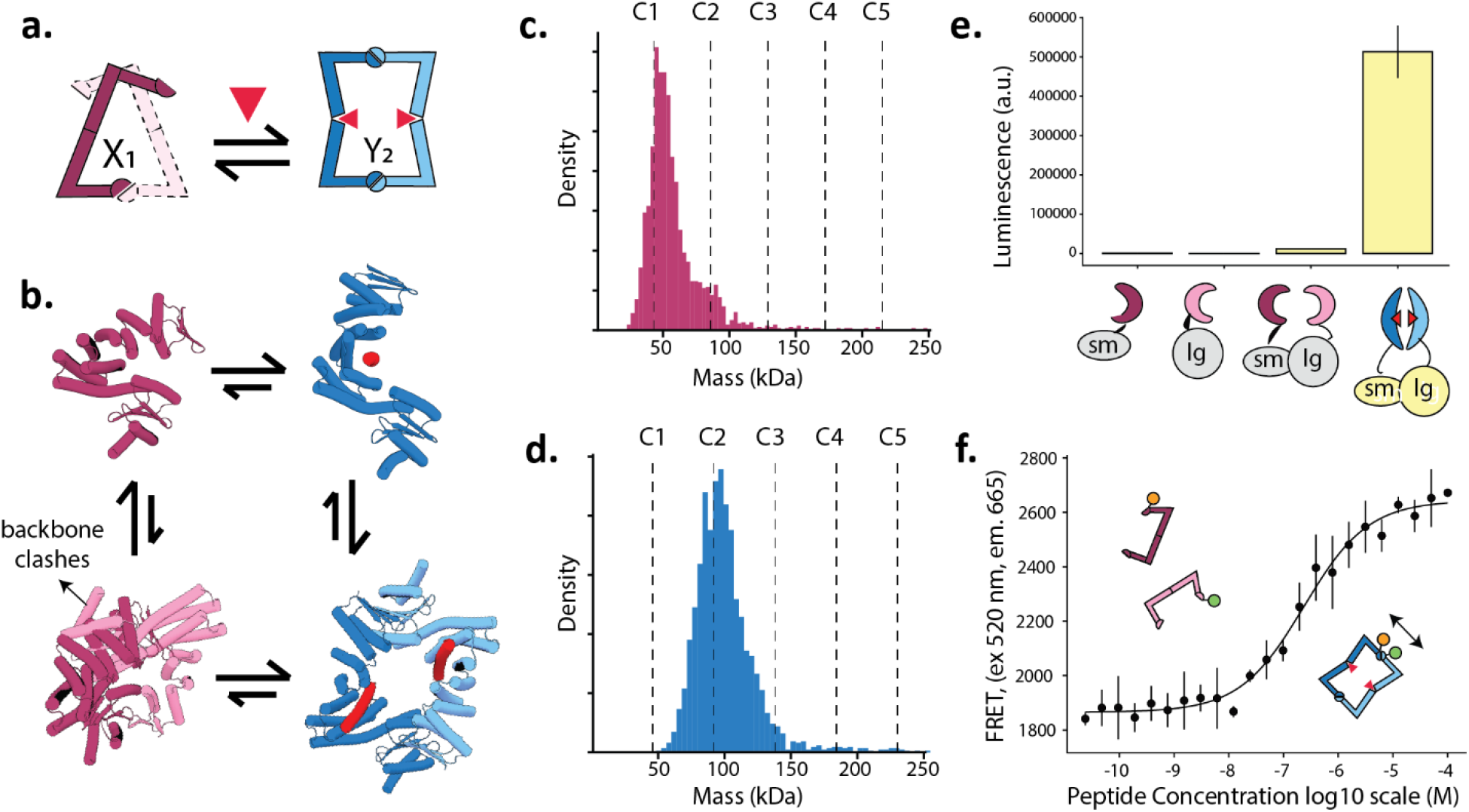
Design of allosterically controlled homodimers. **A.** Schematic showing the equilibrium between a predominantly monomeric apo-state protein (dark pink: first monomer, light pink: second monomer that would sterically clash upon binding), and a predominantly dimeric Y state (blue), where steric clashes that disfavor dimerization are relieved when peptide binds to monomer**. B.** Modelled states of IHA10 in the absence (pink) and presence (blue) of peptide (red). Weight of the equilibrium arrows indicate their relative energies. **C,D.** Mass photometry on 10 nM of IHA10 without (top) and with (bottom) 5 μM peptide. **E.** Luminescence generated in the presence of peptide when lgbit and smbit tagged IHA10 components are mixed at 10 nM each, and 20 μM peptide is added. Luminesence from control samples of smBit-only, lgBit only and lgBit-smBit mixtures without peptide is also plotted for comparison. Average and standard deviation of 10 technical replicates is shown. **F.** FRET signal measured across an effector peptide concentration series where the FRET labelled components are kept constant at 5 nM each and the peptide is titrated from 23.8 fM to 100 μM in two-fold steps. The average and standard deviation of 4 measurements at each concentration is shown. A response curve was fitted to the data (black) by nonlinear regression.

To test the peptide-dependent assembly of these designed dimers, we fused each design separately to one component of a split nanoluciferase^38^ (called lgBit or SmBit respectively). lgBit and SmBit fused designs were mixed in equimolar concentrations (5 nM) with an excess of substrate, and luminescence was measured in the presence and absence of peptide for each protein. In 2/12 cases, addition of 1μM effector induced a greater than 5x increase in luminescent signal (**EFig.22a-b**). For one of the designed proteins, IHA10, addition of 20 μM of peptide cause a 50-fold increase in luminescence signal over controls (**Fig 5b,e**). This enhancement of luminescent signal is consistent with peptide-driven dimerization.

To further investigate effector induced IHA10 dimerization, we used MP, SEC, and Förster Fluorescence Resonance Energy Transfer (FRET). At a protein concentration of 10 nM, mass photometry revealed a nearly complete transition from monomer to dimer upon the addition of 10 μM peptide (**Fig 5b,c,d**). To determine the effector concentration dependence of dimerization, two different IHA10 samples were labeled with donor and acceptor fluorophores, respectively, and then mixed at equimolar ratios so that dimerization should result in enhanced FRET signal. Titration of peptide with each labelled IHA10 construct at 5 nM resulted in an increase in FRET signal with a transition midpoint of ∼750 nM of added peptide, consistent with peptide-driven dimerization. The thermodynamic driving force for dimerization becomes stronger as the monomer concentration increases; we found that IHA10 displayed some degree of dimerization by SEC at 1 μM in the absence of the peptide (**EFig.22c**), but the occupancy of the dimer state is enhanced by the addition of peptide (**EFig.22c**), as evidenced by a clear shift in elution profile. We found that a single alanine substitution at the dimer interface substantially reduced dimer signal even at micromolar protein concentrations, while retaining the ability to allosterically respond to peptide (**EFig.22d-f**). These data suggest that tight control of the monomer-dimer transition by allosteric effector can be achieved through simple rigid-body conformational transitions that relieve steric clashes at an interface, and that this sensitivity can be mutationally tuned.

### Design of allosterically triggerable disassembly

Finally, we explored the possibility of designing assemblies that can be allosterically weakened by effector-binding, such that they disassemble into smaller components. This addresses a major current protein design challenge: the construction of packaging systems that can release a payload upon encountering specific signals that trigger their disassembly^39,40^. Efforts thus far have relied on direct effects at protein-protein interfaces rather than allosteric coupling^41,42^. To explore the applicability of our approach towards driving the disassembly of larger cage-like protein architectures into smaller parts, we sought to design D3 and D5 homomeric assemblies comprising 6 and 10 hinge-containing subunits, respectively, that disassemble into their constituent cyclic components in the presence of the effector peptide. We generated *in silico* a wide range of fusions of hinges to designed C3^43^ and C5^44^ symmetric oligomers (**Fig 6a**), using WORMS^28^ and proteinMPNN^29^, and docked these into D3 and D5 assemblies using RPXDock^45^ such that the exposed N-terminal helices of the opposing cyclic oligomers pack against one another along a dihedral plane of symmetry (**Fig 6b**). We used ProteinMPNN^29^ to design the interface across the dihedral axis between the C3/C5 oligomer and the hinge module. The dihedral interfaces were chosen to be very small (<1000A^2^) so that the free energy of peptide binding would be sufficient to drive disassembly.

**Figure 6.**
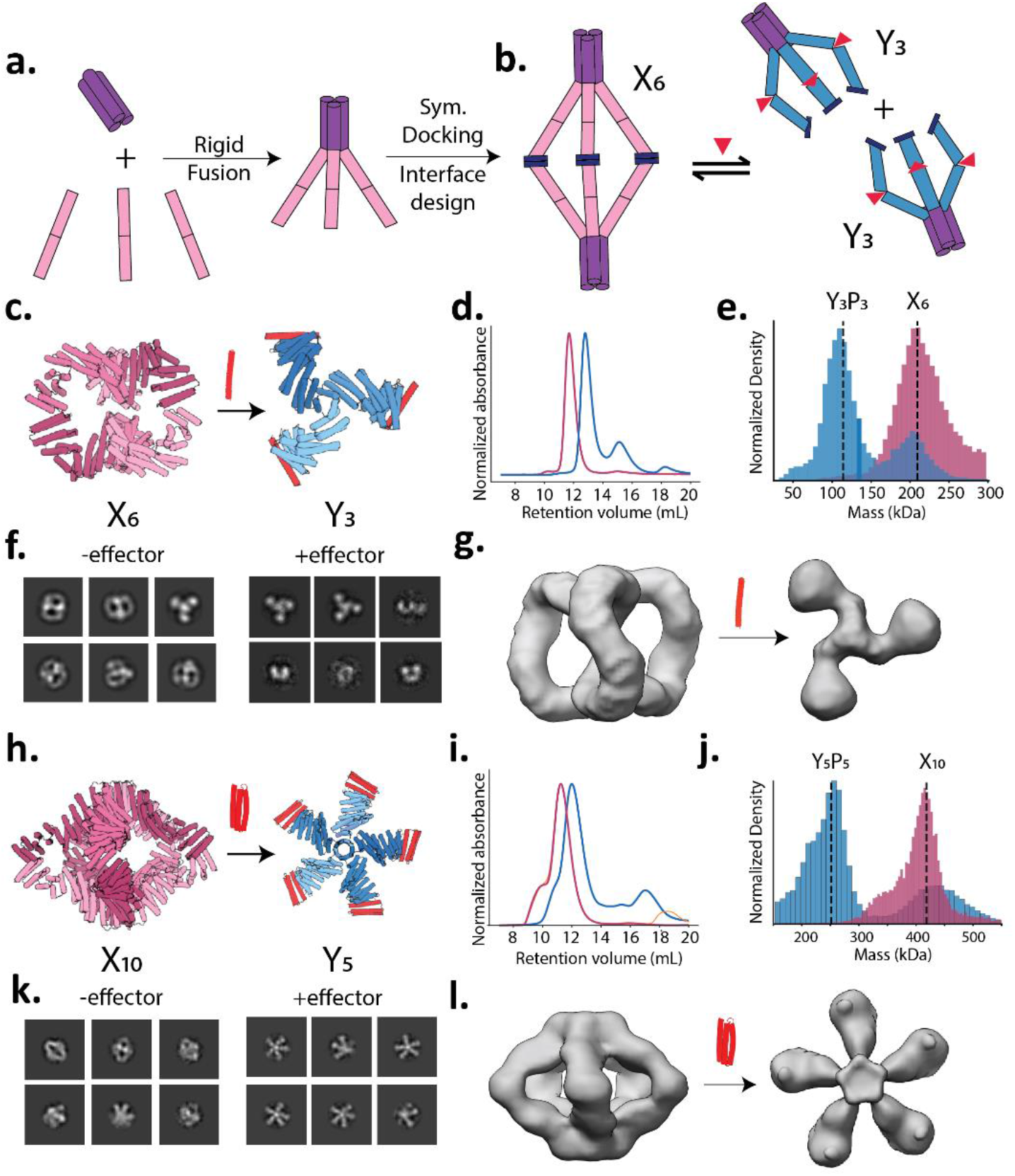
Design of allosterically inducible dihedral cage dissociation. **A.** Fusion strategy for dihedral design; each monomer in a C3 oligomer (purple) is rigidly fused to the hinge protein in the X state (pink). These cyclic components are subsequently docked into dihedral assemblies. **B.** Architecture and behavior of switchable dihedrals. Left, hexameric D3 dihedral generated through symmetric docking and design of new interfaces (dark blue) between C3 components, and right, disassembly of the D3 dihedral into C3 components upon binding to effector peptide. **C.** Design models of D3_29 in the apo (left, pink) and peptide-bound (right, blue) conformations. **D.** SEC chromatogram of 5 μM D3_29 in the absence (pink) and presence (blue) of 10 μM peptide (red). **E.** MP on 100 nM D3_29 in the absence (pink) and presence (blue) of 2x molar excess of peptide, with dashed lines indicating expected theoretical masses of the two states. **F**,**G**, nsEM 2D class averages (f.) and 3D reconstructions (g.) for D3_29 in the absence (left) and presence (right) of saturating amounts of peptide (red). **H.** Design models of D3_29 in its expected oligomeric state in the apo (left) and peptide-bound (right) conformations. **I.** SEC chromatogram of 5 μM D5_05 in the absence (pink) and presence (blue) of 10 μM effector protein. **J.** MP on 100 nM D5_05 in the absence (pink) and presence (blue) of 2x molar excess of peptide, with expected masses indicated with dashed vertical lines. **K,L.** nsEM 2D class averages (K.) and 3D reconstruction (l.) for D5_05 in the absence (left) and presence (right) of saturating amounts of effector protein (red).

We expressed and purified 31 designs and found that 14 were soluble and had monodisperse SEC profiles (**EFig 23,24**). For the 9 soluble D3 designs, 3 exhibited a distinctive shift in the SEC elution profile in the presence of 2x molar excess of peptide, with the oligomer peak eluting later, indicating a reduction in size upon binding (**Fig 6c,d, EFig. 23**). Mass photometry analysis of D3_29, revealed a shift in mass distribution in the presence of effector consistent with a transition from a hexamer to a trimer, suggesting that D3_29 undergoes peptide-mediated disassembly into its C3 components (**Fig 6e**). In contrast, the 5 soluble D5 designs did not exhibit any discernible disassembly by SEC when effector peptide was added (**EFig 24**). We hypothesized that the D5 assemblies may have more of a cooperative advantage than the D3 assemblies (since 5 new protein interfaces are formed rather than 3) and that utilizing an effector variant with higher affinity for binding to the hinges could thermodynamically outcompete the D5 interfaces. To test this, we employed a globular protein version of the effector, 3hb21, which binds the hinge 10-fold more strongly^46^. We found that, in contrast to the peptide effector, the stabilized protein version of the effector induced the disassembly of D5_05 (**Fig 6h,i**): MP indicated conversion of decamers into pentamers when the effector protein is added, consistent with the D5 to C5 transition (**Fig 6j**). This ability to design systems that respond differently to different effectors further illustrates the tunability of our allosteric design approach.

We characterized the structures of D3_29, and D5_05 using nsEM in the absence and presence of effector. 2D class-averages show multiple top and side views of D3_29, and a 3D reconstruction is consistent with the formation of the D3 dihedral design model in absence of effector (**Fig 6f,g, EFig. 25**). In contrast, in the presence of the peptide, C3s are the predominant species present, corroborating that the addition of peptide breaks the weak dihedral interfaces, consistent with the design goal (**Fig 6f,g, EFig. 25**). Similarly, D5_05 assembled into a D5 dihedral resembling the design model in the absence of the effector protein, but in the presence of effector, disassembled into free C5 components (**Fig 6k,l, EFig. 26**). Thus, our design approach enables the construction of cage-like architectures that undergo effector-triggerable disassembly.

### Conclusions

Our work demonstrates that a wide array of dynamic and allosterically switchable protein assemblies can be generated through geometrically precise combinations of two-state hinge modules and protein-protein interaction modules. Our design strategy effectively couples the binding interfaces across substantial atomic distances (>10Å) without the explicit design of an allosteric sidechain communication network. Instead, tuning of the relative free energies of the different rigid-body coupled states was critical to success; in particular, the design of imperfection in the ground state – in the form of steric clashes, incomplete closure, or weak interfaces – was key to ensure the ability to switch. Many of the allosterically switchable oligomers reported here have no direct functional analog in nature – in terms of the architectures that they can toggle between, and the effectors they respond to – and expand the allosteric potential of proteins into regions of biophysical space unexplored by natural evolution.

There are several exciting applications and next steps for our switchable designs. First, the cyclic and dihedral nanostructures can serve as building blocks in the construction of higher-order switchable symmetric nanomaterials, including sheets^47^, fibers^48^, crystals^49^ and polyhedral cages^50^, whose assembly or disassembly can be precisely controlled by addition of effector. Allosteric controllable rings and cages are particularly attractive for drug packaging and delivery, as assembly can be triggered following mixing with the cargo to be packaged, or disassembly triggered in a target location or cellular context^39,40^. Second, our designed inducible homodimers can be used to drive association of cell surface receptors upon effector binding, enabling feedback control of signaling in adoptive cell therapies^51^. Finally, our approach provides a route towards the design of protein systems that can convert energy into mechanical work, like molecular walkers^52^ and pumps^19^, which in Nature rely on defined transitions between alternative structural states driven by allosteric binding events.

## Supporting information

PDBs_in_paper

## Acknowledgements and Funding

We thank D. D. Sahtoe, R. D. Kiber, Y. Hsia, N. Bethel and A. Favor for helpful discussions and K. VanWormer and L. Goldschmidt for technical support. We also thank Xinting Li and Mila Lamb for mass spectrometry support. This work was supported by the Washington Research Foundation Postdoctoral Fellowship (GR027504, Ar.P.), a National Science Foundation Graduate Research Fellowship (DGE-2140004, A.I.), a Human Frontier Science Program Long Term Fellowship (LT000880/2019, F.P.), the Audacious Project at the Institute for Protein Design (A.B., Ar.P., An.P., A.I., D.B.), a National Energy Research Scientific Computing Center award (BER-ERCAP0022018), the Howard Hughes Medical Institute (D.B.), the Open Philanthropy Project Improving Protein Design Fund (P..L., C.W.D., D.B.) a gift from Microsoft (D.B.), and a grant from DARPA supporting the Harnessing Enzymatic Activity for Lifesaving Remedies (HEALR) program (HR001120S0052 contract HR0011-21-2-0012, D.B.).

## Author Contributions

Ar.P., A.I and F.P. conceived of the hinge-based switchable oligomer concept Ar.P designed the switchable ring systems, with computational assistance from F.P., P.J.Y.L. and C.D. A.I. designed the switchable dihedral systems with computational assistance with A.B. Ar.P and An.P. screened and experimentally characterized the ring assemblies, with assistance from R.S. A.I. screened and experimentally characterized the dihedral assemblies with assistance from R.S. and A.P. C.W. and R.S. obtained and analyzed cryoEM data for sr312 and sr322, with supervision from A.J.B. Ar.P, F.P. and D.B. wrote the manuscript, with comments and assistance from all authors.

## Data Availability

All data is available either in the main text or as supplementary materials. PDB models and sequences for designs shown in main text figures are available in attached Supplementary Data.

## METHODS

### Generation of scaffold library of cyclic Y-state oligomers

Cyclic oligomers were constructed through the modular fusion of two classes of de-novo proteins: (1) Three hinge proteins (cs074, cs221, js007) that were previously confirmed to switch between two defined conformational states, ‘X’ and ‘Y’, in response to peptide-binding^11^ and (2) Heterodimeric alpha-beta proteins (LHDs) that were designed to reversibly associate and dissociate in dynamic equilibrium^12^. To create helical surfaces that would facilitate modular fusion between LHDs and hinges, and to sample a diversity of possible angular fusions between these components, we used the HFuse^28^ software to computationally generate a library of designed helical repeat (DHR) fusions to each monomer within an LHD (an example is shown in **Efig1b**). The junctions between LHDs and DHRs were then redesigned with Rosetta FastDesign with backbone movement to stably pack the junctions between them. These designs were then predicted with alphafold v2^30^ (AF2), and we filtered for designs that reported a model pLDDT (predicted local distance difference test) score>88 and C-α rmsd (root mean square deviation) <2.5 Å to the original Hfuse backbone. Dimer LHD-DHR fusions with two helical termini available for further fusion, one at the C-terminus, and the other at the N-terminus, were then used as building blocks in the subsequent steps, with each terminus being utilized as a surface for fusion to the N and C termini of a hinge (**Fig 1c**). The WORMS^28^ software was used to generate a library of rigid fusions between the LHDs and Hinges (in their peptide-bound ‘Y’ state) that would result in cyclic closure into a C2, C3, C4 or C5 oligomer in the presence of peptide. Splicing of up to 2 terminal helices on the hinges, and 150 terminal residues on the LHD-DHR monomers was permitted during the WORMS-fusion process to generate robust junctions, while preserving binding function at both LHD and hinge interfaces. ‘Yn’ Oligomers with a monomer length of under 450 amino acids were selected for further design. For each peptide-bound Y-state monomer generated by WORMs, we generated models of the X-state structure that it would adopt in the absence of peptide. We did this by aligning segments of the Y-state chimeric protein, N and C terminal to the hinge region, to the known X-state of the corresponding parent hinge. This yielded a batch of X-state models that could be used for comparisons in AF2-filtering in subsequent steps.

### MPNN Redesign of junctions and AF2 filtering for fusion quality

To ensure the solubility and rigidity of these fusions, we redesigned the residues at the WORMS-generated junctions with ProteinMPNN^29^, while fixing the amino-acid identities of residues (1) key to the conformational switching of the hinge and (2) that mediate assembly into rings at the LHD interfaces. The former category included residues that directly interact with the peptide at the cleft as well as ones that assist in packing the backside of the hinge when the peptide is bound **(Efig 1a**). 4 sequences were generated per design using proteinMPNN. These sequences were then predicted as monomers with AF2 in the absence of the peptide with 3 recycles, yielding putative X-state structures. We filtered for MPNN-designed ring proteins with pLDDT>86, and a C-α rmsd of under 2.75 Å to the X-state design model that was generated as described above.

### Prediction of X-state oligomers

To generate the predicted X-state oligomer (Xm), for each design that passed this initial filter, we iteratively docked 5 AF2-predicted monomers end-to-end along their LHD interfaces to generate a series of oligomers, from dimer to pentamer. We recorded the distance between the N-terminal LHD-domain of the first monomer and the C-terminal LHD-domain of the nth monomer in the chain at each docking step by computing the shortest distance between all possible pairings of backbone carbon atoms across the two domains (**Efig.2a**). Oligomeric states in which this atomic distance is minimal were identified to measure the closest approach of the X-state oligomer to cyclic closure. We filtered for designs where this distance was <24Å to enrich for designs that are likely to close in the X-state rather than extend into filaments (**Efig.2b**). A subset of these filtered designs was then manually selected for experimental characterization to span a range of symmetries, shapes and cognate effectors. To determine the extent to which the monomers must individually bend in order to mediate closure in the X state, we predicted them in their assembled form using alphafold multimer v3. We used rigid docks of the X-state AF2-predicted monomers prepared in the previous step as templates for prediction with 3 recycles. In 23/26 cases, the monomers underwent some degree of flexible deviation away from the X-state to close the ring with optimally or near-optimally satisfied LHD interfaces. To quantify this deviation in the closed cases, we measured the rmsd between the ring-incorporated monomer and the free monomer predictions, as shown in **Efig 2c**. AF2-multimer v3 did not predict C4 oligomers or higher as closing into rings, preferring to model them as dihedrals with unsatisfied LHD interfaces.

### Design of static control rings

To assess the importance of designing junctions to target optimal closure in the Y state, but suboptimal closure in the X state, we implemented a modified design procedure where the unbound X state ring is optimized for closure, rather than the Y-state ring. We used WORMS to screen for LHD-hinge fusions that resulted in the perfect end-to-end closure of an X-state cyclic ring containing between 3 and 5 subunits. ProteinMPNN was used to design the junctions between the components, while preserving the key functional residues in the hinge domain, as described above. AF2 was used to predict the monomer structures and validate the fusion junction designs. Designs with a pLDDT >86 and AF2-predicted C-α rmsd <1.5A to X-state design were selected for further characterization.

### Design of inducible homodimers

The WORMS-based fusion approach described above was used to target a C2-symmetric homodimeric assembly in state Y, through the fusion of LHD-DHRs to hinges. ProteinMPNN was used to redesign the junctions of the monomeric subunits, followed by folding with AF2 in the absence of peptide to generate an X-state monomer. We filtered for designs which were predicted with a C-α rmsd <2.75Å to the expected X-state backbone and pLDDT>86. After this, the X-state predicted monomers were docked along one of their LHD interfaces. We used Rosetta to filter for designs with backbone clashes (Rosetta fa_rep score > 10000) in this docked state, corresponding to a sterically prohibited dimer form. The filtered designs were manually inspected to ensure that the LHD produced substantial backbone clashes in the X state, and a subset of 12 were then experimentally characterized.

### Design of symmetric C3 and C5 subcomponents for dihedral assemblies

The WORMS protocol was used, without symmetry constraints, to generate rigid fusions of hinges to previously validated C3^43^ and C5^44^ oligomers. The C-terminus of hinge cs221 was rigidly fused to these cyclic oligomers in an orientation that ensured the N-terminus of the hinge would be accessible for dihedral docking in state X. Fusion junctions between hinges and cyclic oligomers were then redesigned with ProteinMPNN while preserving the oligomeric interfaces of the parent scaffolds along with key functional residues of the hinge protein. To increase the number of oligomers for downstream docking applications, 8 sequences were generated for each rigidly fused oligomer. Redesigned proteins were then predicted as monomers using AF2 with initial guess^53^ and designs with a pLDDT > 85 and an RMSD < 2.5Å were selected for dihedral docking.

### Docking of subcomponents

RPXDock^45^ was used to sample D3 and D5 assemblies containing two copies of a C3 or C5 hinge-fusion, respectively. Docking was guided towards the first 36 residues of component monomers, such that the N-terminal helices of the opposing cyclic oligomers packed against one another along a dihedral plane of symmetry. Docked designs were then sequence optimized along the dihedral interface using ProteinMPNN with the tied residues feature, such that the interface for each chain pair along the dihedral axis contained identical residues. To evaluate and filter for these newly designed dihedral interfaces we extracted C2-symmetric dimeric subunits and predicted their structure with AF2 with initial guess^53^. Designs passing metrics of pLDDT > 85 and C-α RMSD < 2.5 Å were then chosen for further characterization.

### Recombinant expression and purification

Genes were codon-optimized for expression in Escherichia coli (E Coli). DNA fragments encoding designed proteins were ordered as eblocks from IDT and cloned into custom plasmids bearing a T7-promoter driven expression system with a C-terminal SNAC cleavage site and 6xHis-tag (“Protein-GSHHWGSTHHHHHH”) using Golden Gate Assembly. All proteins were expressed in NEB BL21(DE3) E. coli cells using TBII (MpBio) autoinduction media, which was supplemented with ZYM-5052, trace metal mix, 2mM MgSO4 and 50 mg/ml Kanamycin. 50 mL expression cultures were grown at 37C for 6 hours followed by 20C for 24 h with shaking at 225 rpm throughout.

Cells were then harvested by centrifugation at 5000xg and resuspended in 15 mL of TBS lysis buffer (300 mM NaCl, 40 mM Tris, 40 mM Imidazole). Cells were lysed by sonication in the presence of 1 mM Dnase, 1 Pierce™ Protease Inhibitor Mini Tablets, EDTA-free per 100 mL and 1 mM PMSF added immediately before lysis. Cell debris was pelleted by centrifugation at 20000g for 40 min. The supernatant was then added to ∼1mL Ni-NTA Metal affinity Chromatography resin to separate the protein from impurities in a vacuum manifold. The protein was washed with 10x bead volume of TBS (300 mM NaCl, 40 mM Tris, 40 mM Imidazole pH 8) and protein was eluted in 2 mL of 300 mM NaCl, 40 mM Tris, 500 mM Imidazole. Eluted protein was then further purified with size exclusion chromatography (SEC) using Superdex 200 Increase 10/300 GL columns in TBS (40 mM Tris, 300 mM NaCl, pH 8) with 1 mL fractions. Final concentrations were estimated using UV-280 absorbance with a NanoDrop 2000/2000c, relying on molar extinction coefficients and molecular weights predicted from the sequence. LC-To confirm the protein sequence, we measured the molecular mass of each protein by mass spectrometry; intact mass spectra were obtained via reverse-phase LC/MS on an Agilent G6230B TOF on an AdvanceBio RP-Desalting column, and subsequently deconvoluted by way of Bioconfirm using a total entropy algorithm. Sequences picked for further characterization beyond SEC-binding assays were re-ordered as precloned genes from IDT in pet29B expression vectors. The cognate peptides for these proteins (cs074B, cs221B and js007B) were chemically synthesized as previously described^11^.

In constructs with designed disulfides (sr312_y_staple), the expression and purification were the same as described above, with two modifications: (1) 1 mM TCEP was added to the lysis buffer to prevent premature disulfide formation during purification (2) copper phenanthroline was added to the IMAC elution at a final concentration of 10 mM, and the resulting mixture was incubated overnight to encourage full formation of the disulfides.

### Size-Exclusion chromatography binding experiments

To determine the effects of peptide-binding on oligomerization state, we measured shifts in SEC profile in the presence and absence of ∼2X molar excess of peptide (**Fig 2, Efig5**). Protein assemblies and peptide were diluted into TBS (300 mM NaCl, 40 mM Tris, pH 8) to a final monomer concentration of 5 μM and a final peptide concentration of 10 μM in a final volume of 700 μL. Peptide-free and peptide-bound samples were injected serially using an automated FPLC system (AKTA Pure) with a flow rate of 0.5 mL/min on a Superdex 200 Increase 10/300 GL column. Absorbance signals at UV230 and UV280 were measured to monitor the elution profile of protein across the run. For constructs that included a GFP tag, UV473 absorbance was also measured. For the SEC titration series of sr312 shown in **Efig. 20a**, the unbound fraction was calculated by estimating the area under the X3 peak in each measurement with UNICORN 7.3 and dividing it by the area under the X3 peak in the absence of the effector peptide (where 100% of the protein is unbound). The bound fraction is then calculated by subtracting this value from 1. SEC traces of multiple runs that are shown as overlays were run subsequently on the identical FPLC system and column, on the same day, and using the identical buffer to ensure ideal comparability.

### Fluorescence polarization

All FP binding assays were conducted in 96-well black-bottom microplates (Corning 3686) at room temperature (∼25°C). Fluorescence measurements, including parallel intensity, perpendicular intensity and polarization, were taken in a Synergy NEO2 plate reader, with a 530/590 filter cube. For each design, four replicate titration series were prepared per plate by serial two-fold dilution of a starting stock of 20 μM protein into TBS (300 NaCl, 40 mM Tris, pH 8.0 buffer) across 24 wells. TAMRA-labeled peptide was kept across this series at a constant concentration of 1 nM. The final volume of peptide-protein mixture in each well was kept constant at 80 μL. Measured polarization signal was fitted to the following equation to determine K_D_.

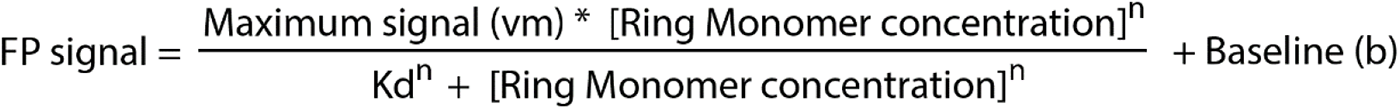

Where, b, Vm, n and K_D_ are fit by nonlinear regression to a set of polarization signal and protein concentration values using the optimize_curvefit function within Scipy. An average of K_D_ from four replicates and their standard error are reported in **Efig.8**.

### TAMRA size exclusion chromatography

TAMRA-labelled peptide at varying concentrations (10,5,2,1,0.5,0.25 μM) was separately mixed with a constant 1 μM concentration of sr312 in a final volume of 200 μL in TBS (40 mM Tris, 300 mM, pH 8.0). Mixtures were serially injected with an autosampler on a High-performance liquid chromatography system (Agilent 1260 Infinity II LC). In addition to absorbance at UV280, TAMRA fluorescence at 590 nm was recorded with an excitation wavelength of 570nm to specifically monitor the elution of labelled peptide. SEC traces were collected over a 9 minute interval from a Superdex 200 GL5/150 Increase column at a flow rate of 0.35 ml/min in TBS (300 NaCl, 40 mM Tris, pH 8). 100 μL fractions were collected across the elution run time.

### Characterization of complexes by Mass Photometry

All mass photometry measurements were carried out in a TwoMP (Refeyn) Mass photometer. For initial characterization of rings in **Fig 2**, protein and peptide were incubated at 1 μM protein concentration, with or without 10 μM peptide, for 20-25 hours at room temperature to allow the system to reach equilibrium. For MP data shown in **Fig 5**., Dihedral samples were incubated at 5 μM with 2X molar excess of either cs221B or effector protein 3hb21 overnight. Samples were then diluted to 200nM monomer concentration immediately prior to measurement to limit overcrowding of the field of view. A 12-well gasket was placed on each slide. 10 uL of buffer was added to one well of this gasket and the camera was brought into focus after orienting the laser to the center of the sample well. 10 μL of sample was added to this droplet and 1-minute videos were collected with either a large field of view (for ring and dihedral complexes) or a small field of view (for inducible homodimers) in AcquireMP. Ratiometric contrast values for individual particles were measured and processed into mass distributions with DiscoverMP. For each design, a sample of 20 nM Beta-amylase – consisting of monomers (56Kda), dimers(112Kda) and tetramers(224Kda) in equilibrium – was used to arrive at a mass calibration; thereby allowing contrast values to be converted into mass values across tested designs. Expected masses for Xm and Yn species were calculated by multiplying LC-MS estimated monomer masses by the number of subunits in different oligomeric configurations. Distributions were exported from DiscoverMP and plotted with a custom script in python. Gaussian distributions were fit to this peak to estimate observed oligomer masses and mass error using normfit in the Scipy package (**Efig.7**).

For sr322 cooperativity measurements in **Fig 4**. And **Efig 20**., a titration series of GFP-tagged peptide spanning 10 μM to 0.125 μM was mixed with a constant concentration of 3 μM sr312. Solutions were incubated and measured as described. Discover MP was used to fit gaussian distributions to bound and unbound species in the mass distributions, with three technical replicates for each mixture. Proportions of bound and unbound hinge could then be estimated with the following equation:

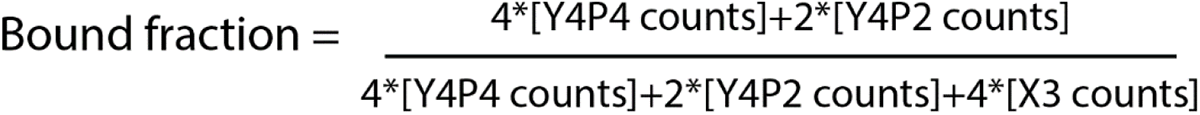

The SciPy optimize_curvefit function was used to fit an occupancy curve to the data collected, allowing for estimation of apparent Kd and hill coefficient (n).

### nsEM on switchable rings and dihedral complexes

Carbon-coated 400 mesh copper grids (01844-F, TedPella,Inc) were first glow-discharged using a PELCO easiGlow cleaning System. SEC-purified proteins were diluted to 0.01 mg/ml with SEC buffer (300 mM NaCl, 40 mM Tris, pH 8.0), and then immediately pipetted onto the glow-discharged grids. The protein solution was allowed to sit on the grid for 1 minute, before being blotted away with Whatman filter paper. 3 uL of 2% uranyl formate stain was added to the grid and then blotted away after 1 minute. A second and third wash of UF stain was added to the grid, allowed to sit for 30 seconds each, before being blotted away. The grid was allowed to air-dry for 5 minutes. Dried grids were then imaged using a FEI Talos L120C TEM (FEI Thermo Scientific, Hillsboro, OR) equipped with a 4K × 4K Gatan OneView camera, at a magnification of 57,000x and pixel size of 2.49 Å. Once a grid-square with satisfactory stain thickness and contrast was identified, EPU software was used to automatically collect 200-400 micrographs across the square. Micrographs were imported into and analyzed using CryoSPARC v4.0.3. 50-100. Patch CTF was used to estimate defocus variation for micrographs. Given the radius of particles, CryoSPARC automatically picked and extracted particles from the CTF corrected micrographs. Particles then were subjected to 2D classification to find 2D averages that could be used as templates for more precise particle picking across the CTF corrected micrographs. After particle picking and extraction from micrographs, a further round of 2D classification was done to find higher resolution averages of the oligomers in various states and orientations.

### Fluorescence Resonance Energy Transfer

FRET constructs for IHA10 were designed by picking two sites on opposing sides of the dimer interface and separately mutating each to cysteine in two point-mutant constructs, IHA10_8C and IHA10_323C. FRET labelling sites were chosen to be close enough to maximize signal in the dimer form (**Efig 22g**), while being far enough from the interface to not sterically block assembly when conjugated with the fluorescent dye. DNA constructs encoding these two point mutants were ordered from IDT as pre-cloned genes in the same expression vector described above. As donor and acceptor dyes, we used AlexaFluor 555 C2 maleimide (donor) and AlexaFluor647 C2 maleimide (acceptor), which were purchased from ThermoFisherScientific. 1mg of each dye was dissolved in 200 uL DMSO to yield a stock solution at 5mM. Cysteine mutants were expressed and purified according to the previously described procedure, with a modification that 0.5 mM TCEP was added to the buffer during lysis, IMAC, and SEC. Additionally, 20 mM sodium phosphate (pH 7.0) instead of Tris-HCl was used as a buffer during SEC. Following the SEC step, a 500 μl solution each of IHA10_8C and IHA10_323C at a concentration of 50 μM was subjected to a 2-hour incubation at room temperature with 500 μM of a single dye (AlexaFluor 555 and AlexaFluor 647 respectively). The labelled samples were then purified by SEC to eliminate excess dye, in a buffer of 20 mM Tris-HCl and 300 mM NaCl at a pH of 8. The FRET titration was conducted at 25°C in 96-well plates (Corning 3686) using a Synergy Neo2 plate reader. The excitation wavelength was 520 nm and emission wavelength was 665 nm (**Fig 5f**). A response curve was fitted to the data by nonlinear regression with a custom python script.

### Luciferase assay for inducible homodimers

Gene fragments encoding inducible dimer designs were ordered as eblocks from IDT and cloned into custom plasmids which included either a C-terminal fusion to the lgBit subunit of NanoLuc or an N-terminal fusion to the smBit of NanoLuc. Ordered sequences also included a C-terminal Histidine tag. As a control for the js007-based hinge designs, we also tagged the corresponding parent hinge, js007A, with each of these subunits to ensure that the luciferase signal is driven primarily by assembly. Proteins were expressed and purified as described above. Luciferase Assays were performed in 40 mM Tris-HCl, 300 mM NaCl, pH 8, 0.05% v/v Tween 20. Reactions were assembled in 96 well plates (Corning 3686) and luminescence was measured in a Synergy Neo2 plate reader (BioTek). LgBit and smBit fused constructs were mixed at equimolar concentrations that ranged from 5-20 nM (reported in figure legend) in a final volume of 80 μL. Mixtures were incubated at room temperature overnight to ensure that the system reaches equilibrium. For single-peptide-concentration comparisons shown in Fig 4, effector peptide was added at a concentration of 20 μM, and 12 replicates were collected for this sample as well as all controls. For titrations shown in Fig S22, a two-fold dilution series was prepared from a starting concentration of 10 μM, and spanned 24 concentrations. The average of four technical replicates across one plate is shown. In all cases, 10 μL of Nano-Glo substrate at 10X dilution was added immediately prior to measurement, with an approximate dead-time of 10-30s.

### CryoEM sample preparation

To prepare the samples, 2 μLs of sr322 with js007 effector peptide (sr322_ js007B), sr312 with cs221B effector peptide (sr312_cs221B), sr322 at 0.971 mg/mL in 150 mM NaCl, 40 mM Tris, pH 8, was applied to glow-discharged C-flat holey carbon grids. Vitrification was performed using a Mark IV Vitrobot at 4°C for sr322_ js007B and sr312_cs221B, 22°C for sr322 with 100% humidity for all. Samples were frozen on glow-discharged 2.0/2.0-T C-flat holey carbon grids for sr322_ js007B and sr312_cs221B, 1.2/1.3-T C-flat holey carbon grids for sr322. Blotting was done using a 5.5-second blot time, a blot force of 0, and a 5-second wait time for sr312_cs221B and sr322_ js007B; a 6.5-second blot time, a blot force of 0, and a 7.5-second wait time for sr322 was used before being immediately plunge frozen into liquid ethane.

### CryoEM data collection

sr322_ js007B, sr322 and sr312_cs221B were collected automatically using SerialEM^54^ and used to control a ThermoFisher Titan Krios 300 kV TEM for sr322 and sr312_cs221B .ThermoFisher Glasios 200 kV TEM both microscopes equipped with a standalone K3 Summit direct electron detector^55^ and operating in super-resolution mode for sr312_cs221B and counting mode for sr322_ js007B and sr322. Random defocus ranges spanned between −0.8 and −1.8 μm using image shift, with one-shot per hole and nine holes per stage move. Altogether, 1398, 3795, 4213 movies with a pixel size of 0.885, 0.4215, 0.843 and a dose of 50, 43, 52 e^−^/Å^2^ were recorded respectively for sr322_ js007B, sr312_cs221B, sr322.

### CryoEM data processing

All data processing was carried out in CryoSPARC^56^. The video frames were aligned using Patch Motion with an estimated B factor of 500 Å^2^. The maximum alignment resolution was set to 3. Outputs were binned to a final pixel size of 1.0288 Å per pixel by setting the output F-crop factor to ½. Defocus and astigmatism values were estimated using the Patch contrast transfer function with the default parameters. In total, 1,614,340 particles were picked in a reference-free manner using Blob Picker and extracted with a box size of 340 for sr322; for sr312_cs221B, and sr322_ js007B a manual picker was first used to pick 590 particles, and 2,804 particles with box sizes of 400, and 256 respectively. An initial round of reference-free two-dimensional (2D) classification was performed in CryoSPARC using 150, 50, 50, classes and a maximum alignment resolution of 6 Å for sr322, for sr312_cs221B, and sr322_ js007B respectively. The best classes were next low-pass filtered to 20 Å and used as templates for a second round of particle picking using Template Picker, resulting in a new set of: 996,592; 971,294; 524,968; particle picks which were extracted with a box sizes of 340, 600, 300 pixels for sr322, sr312_cs221B, and sr322_ js007B respectively. For sr322_b11 only top views along C5 symmetric access were seen, The best 2D class averages were displayed with 66,770 particles. For sr322 and sr312_cs221B: 996,592 and 971,294 particles—were then used for 3D ab initio determination using the C1 symmetry operator, initial ab initio showed density for a clustered species with sr322; and for sr312_221B a preferred orientation failed to produce a good map. For sr322 clustered species 61,497 particles were further processed using non uniform refinement in C1 with a final estimated global resolution 5.91Å. Another round of template picking was used to pick out monomeric sr322 and to pick out more side views for sr312_cs221B with 924,961 and 1,485,952 particles with box sizes of 340 and 600 respectively were next funneled into another round of reference-free 2D classification for sr322, with the best 157,386 particles submitted for Homogenous Refinement in the presence of C1 symmetry. The estimated global resolution of this map was determined to be 6.54 Å. Once symmetry was confirmed in C1, these maps were refined further using Homogenous Refinement in C4 symmetry to an estimated global resolution of 4.44 Å. For sr312_221B, several rounds of two classification were performed to remove excess top views and better classify side views with a final total particle count of 58,251. At this point, an ab initio was generated in C1 in high agreement with the design model and revealing excellent orientational sampling of the input particles. This map was refined with non-uniform refinement and achieved a final estimated global resolution of 4.40 Å. These maps were refined with DeepEMhancer^57^ for sr322, the sr322 clustered species, and sr312_cs221B. Local resolution estimates were determined in CryoSPARC using a FSC threshold of 0.143. 3D maps for the two half-maps, the final unsharpened maps, and the final sharpened maps were deposited in the Electron Microscopy Data Bank under accession numbers EMD-42442, EMD-42491, and EMD-XXX.

### CryoEM model building and validation

The design model of sr322 and sr312_cs221B was used as an initial reference for building the final cryoEM structures. Pymol^58^ and UCSF Chimera^59^ was initially used to break apart the monomeric components and fit them in density. We then further refined the structure using molecular dynamics flexible fitting (MDFF) simulation: Namdinator^60^. This process was repeated iteratively until convergence and high agreement with the map was achieved. Multiple rounds of relaxation and minimization were performed on the complete structures, which was manually inspected for errors each time using Isolde^61^, and Coot^62,63^. Phenix^64^ real-space refinement was subsequently performed as a final step before the final model quality was analyzed using MolProbity^65^. Figures were generated using UCSF ChimeraX^59^. The final structures were deposited in the PDB: 8UP1, 8URE, ####

## SUPPLEMENTARY FIGURES

**Extended Data Figure 1.**
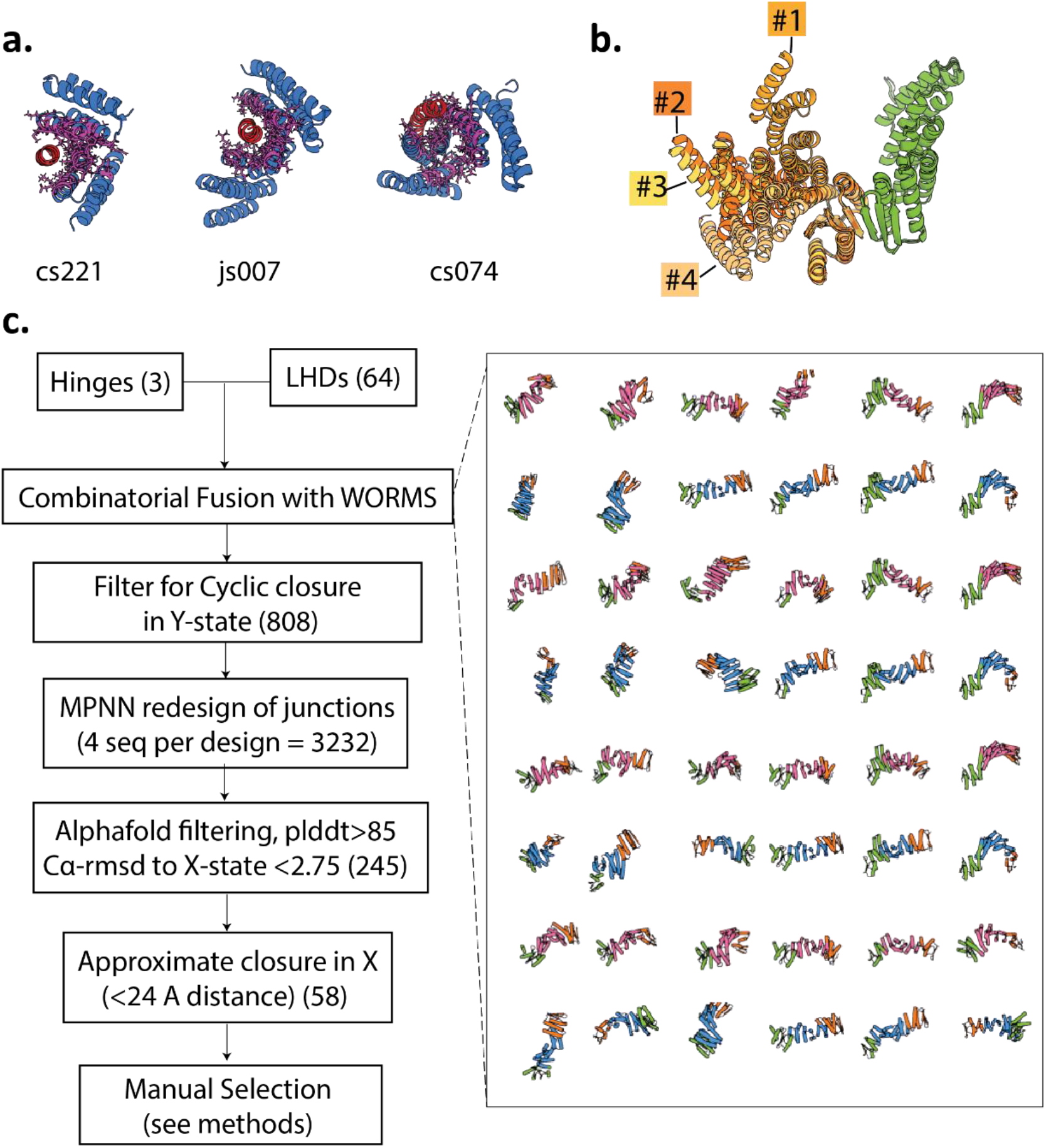
Design procedure for construction of switchable rings, **A.** Three base hinges (blue) used as the “allosteric core” of the switchable ring components, shown in their “Y” state bound to peptide (red). Residues held constant during the design process (in order to preserve the switching properties of the hinges) are shown with red sticks. **B.** Example LHD interface with four alternative DHR fusions showing a range of angles subtended at the ring interfaces. **C.** Left, simplified outline for the finalized design procedure, with numbers of designs produced by or passing each filter at each step. Right, examples of different LHD (orange or green) fusions to hinges (blue X state or pink Y state), to showcase the range of shapes and angles generated during the WORMS sampling process.

**Extended Data Figure 2.**
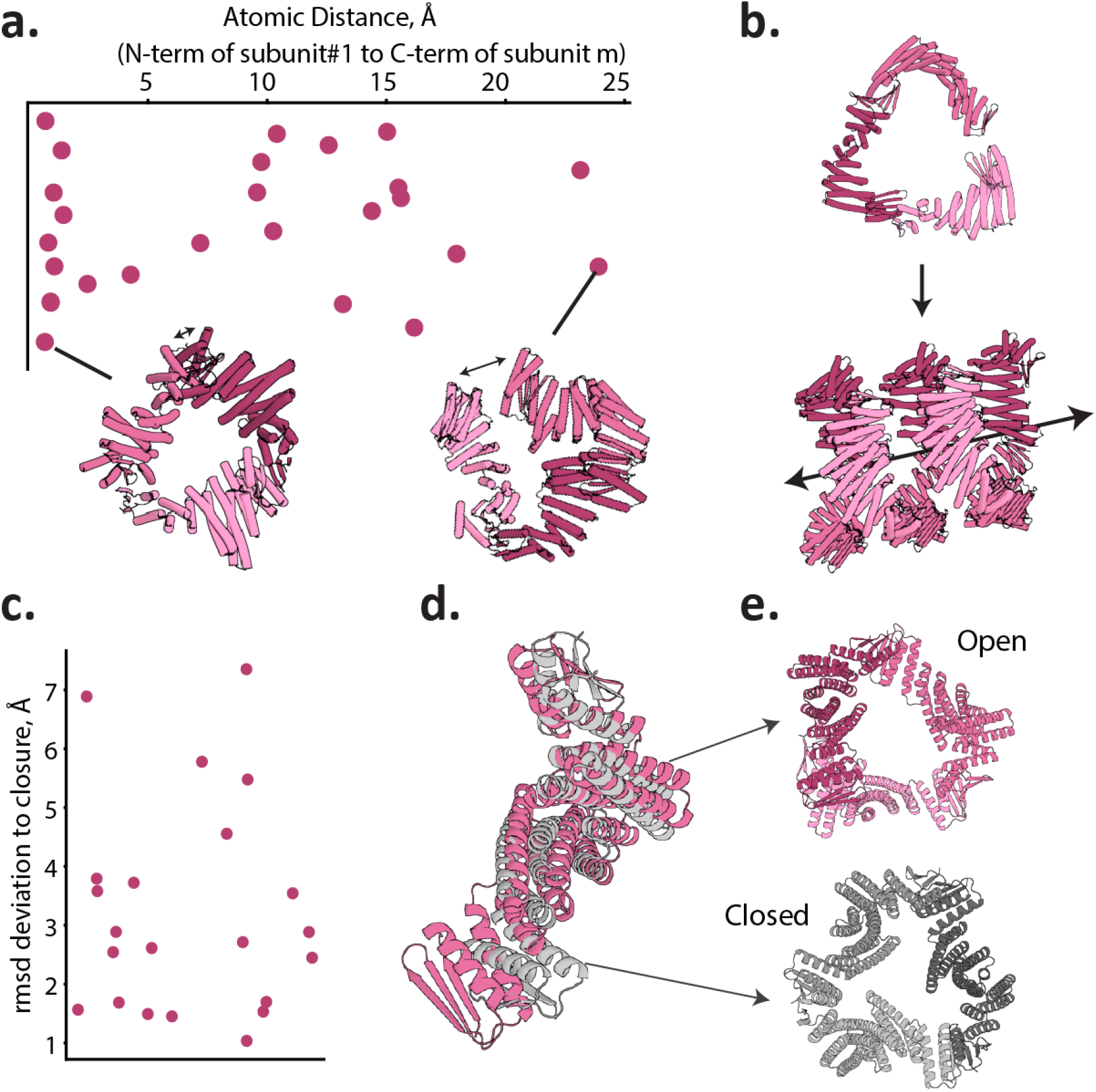
Measurement of imperfect closure geometry in the state “Xm” rings. **A.** Plot of atomic distances between N terminus of the first and C-terminus of the mth docked subunit in the Xm state across tested designs, where the individual subunits are AF2 predictions of the monomer state. The two tested Xm designs with the closest (left) and most distant (right) approach of their ends are shown, where distances are marked with a double headed arrow. Greater distance implies more open rings. **B.** Example of a rejected design whose ring-ends are far enough apart that they associate into open fibers. **C.** Plot of RMSD distance between alphafold multimer predictions of monomers in the free state and in their Xm assembly state. **D.** Example design showing backbone deviation predicted between the free (pink) and assembled (gray) monomer states, modelled using AF2 on a monomer sequence and AF2-multimer-v3 with a template of docked monomers (seen in top of E) respectively. **E**. strained (top) and well-formed (bottom) rings formed by docking the components shown in D as pink and gray.

**Extended Data Figure 3.**
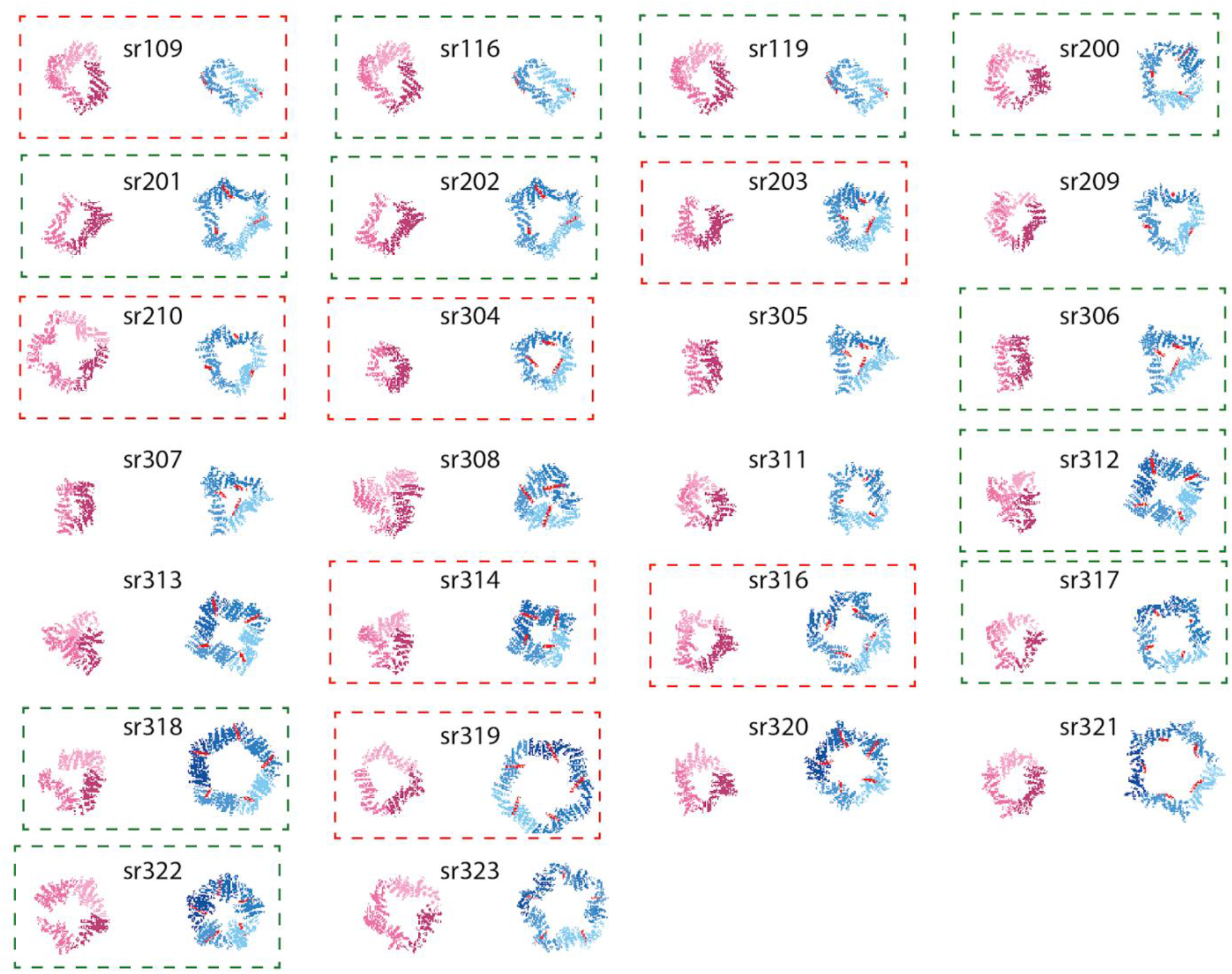
Overview of rings tested for state switching. Xm state of rings is shown in pink, while the Yn state is shown in blue, with peptide colored red. Soluble and SEC-monodisperse designs are indicated with a dashed box (red box: static designs, green box: switchable designs).

**Extended Data Figure 4.**
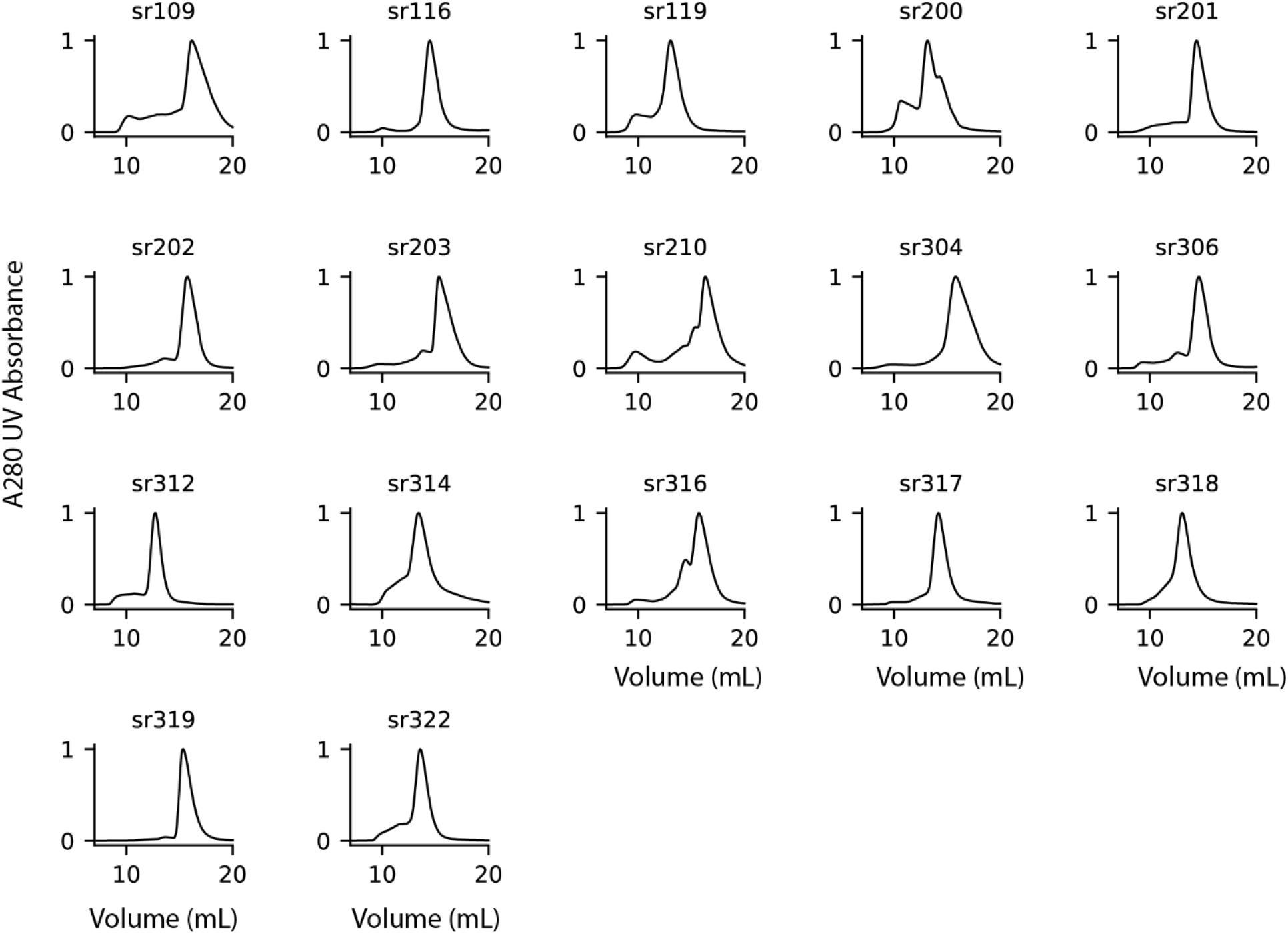
Size exclusion Chromatography (SEC) Purification traces for soluble ring designs.

**Extended Data Figure 5.**
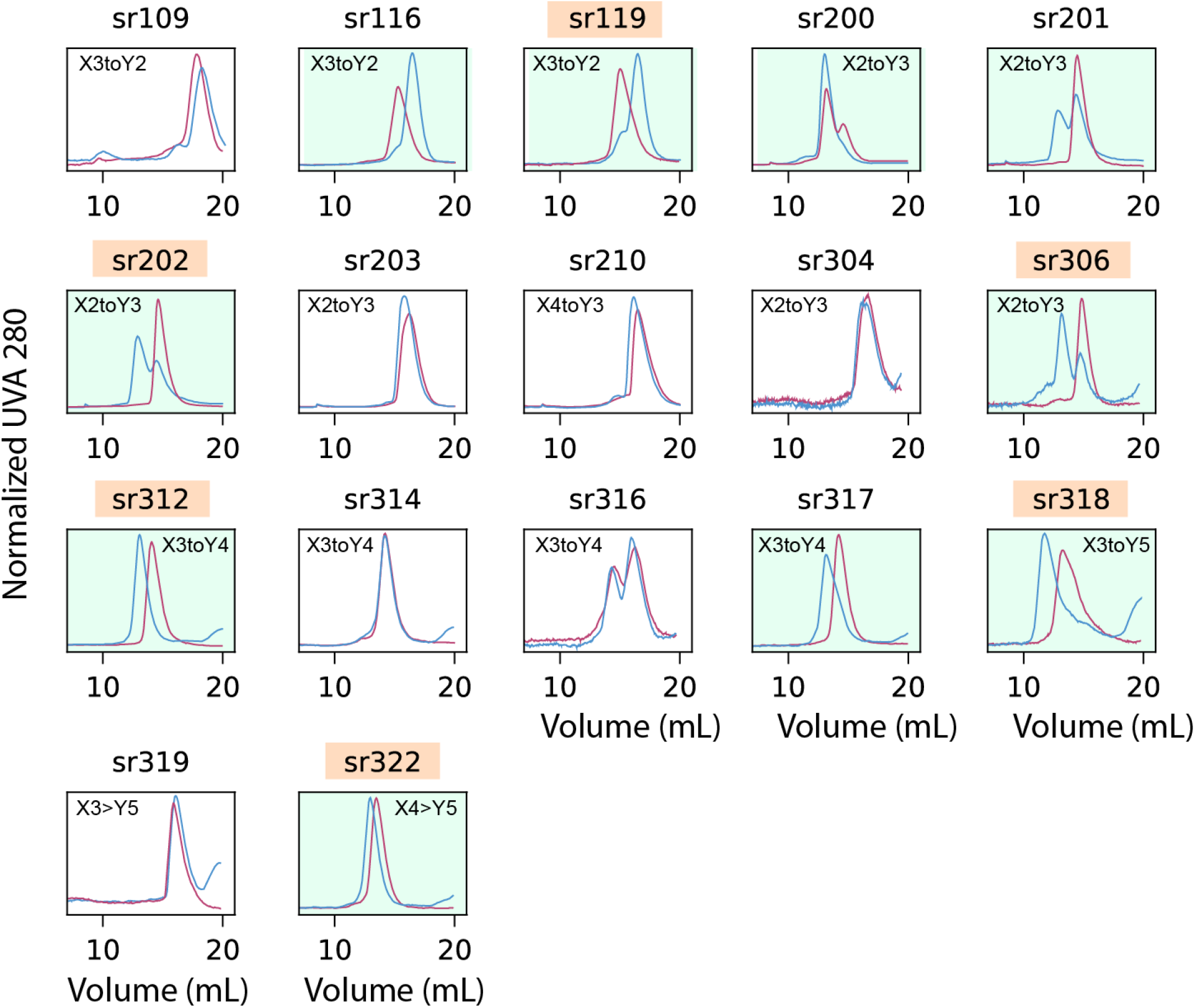
Size-exclusion chromatography analysis of peptide-induced switching. For all designs 5 μM protein was run without peptide (pink) and with 10 μM cognate peptide (blue). Traces represent normalized UV absorbance at 280 nm. Peak near ∼20 corresponds to unbound free peptide. Green colored boxes indicate designs with appreciable change in elution profile when peptide is added, in the expected direction. Names of designs isolated for further testing are highlighted in wheat.

**Extended Data Figure 6.**
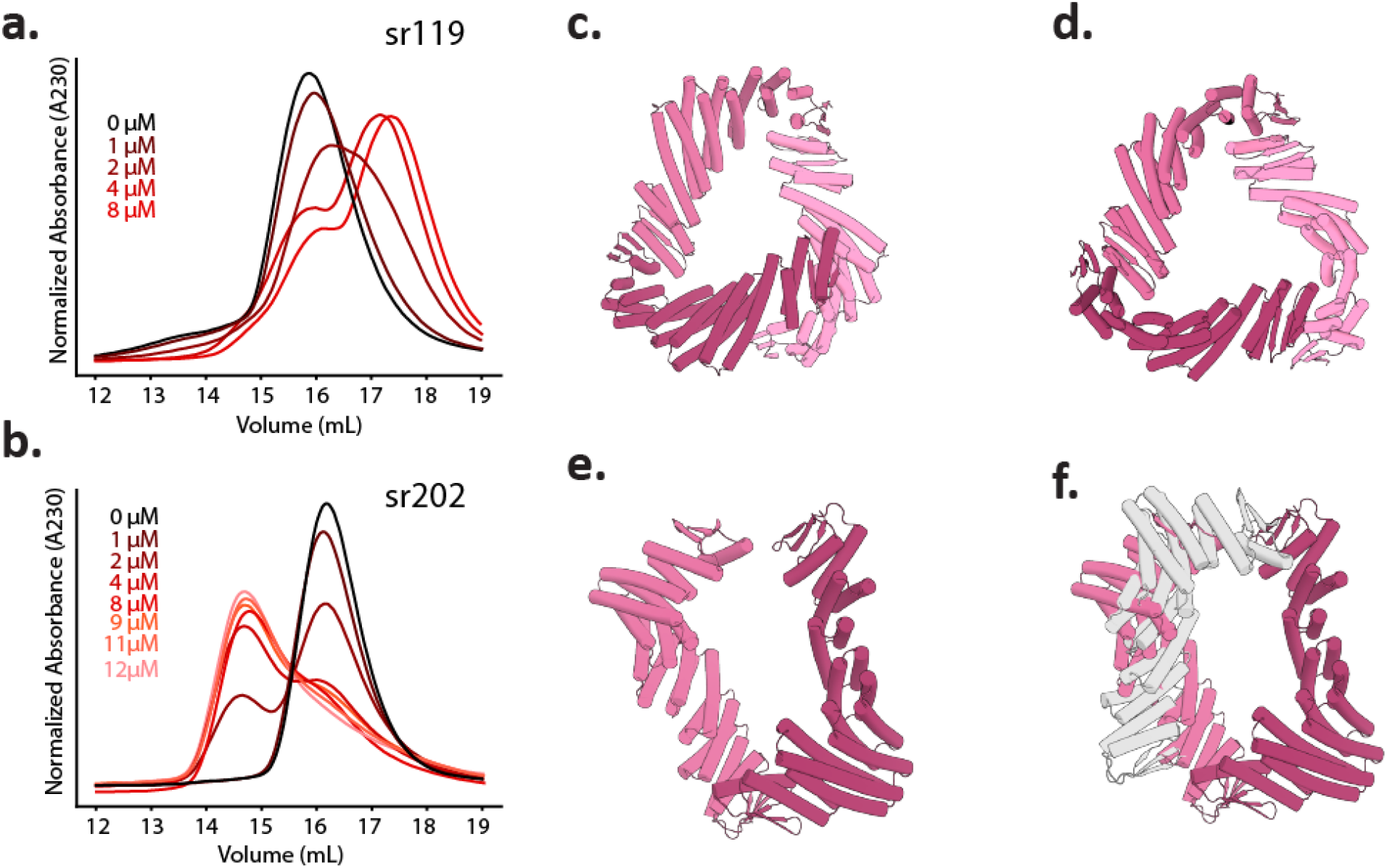
Characterization and behavior sr119 and sr202. **A., B**, SEC titration of peptide, with peptide concentration listed on the left, against a constant monomer concentration of sr119 and sr202 at 5 μM. **C.** sr119 in the strained and sterically clashed state predicted by simple docking of alphafold-predicted free monomers. **D.** Unstrained ring predicted when the three subunits are co-folded with alphafold multimer v3. **E.** sr202 with gap shown prominently in the Xm state. **F.** Docking of a third subunit in the X2 state is sterically occluded by the close approach of the ends of the partially open ring.

**Extended Data Figure 7.**
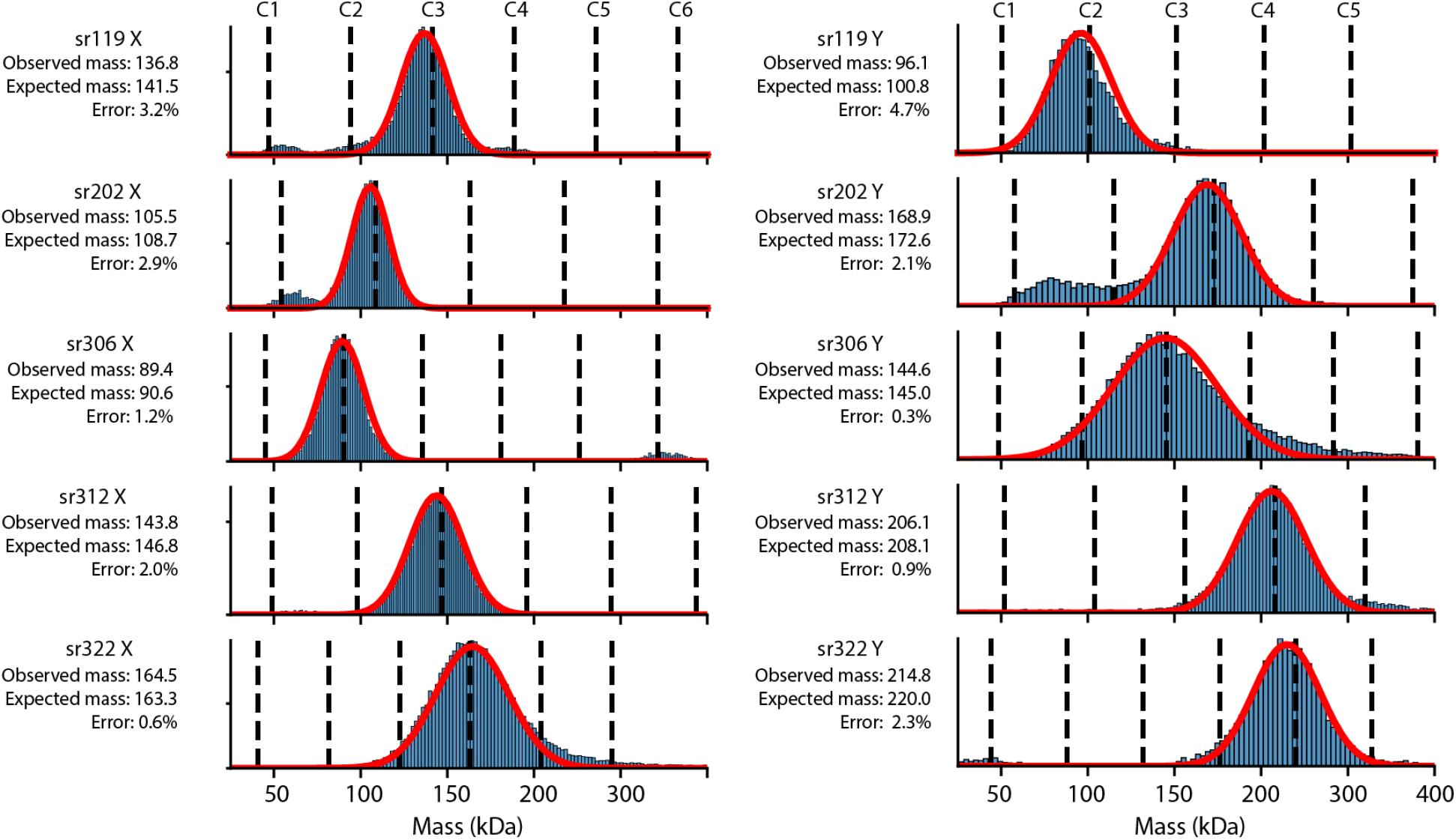
Mass photometry analysis on switchable rings. MP on all designs shown in Fig 2e with mass error, gaussian fit (red) and oligomer masses, both expected and observed, are shown to the left of each panel. Dashed lines indicate expected masses of C1-C6 symmetric homo-oligomers.

**Extended Data Figure 8.**
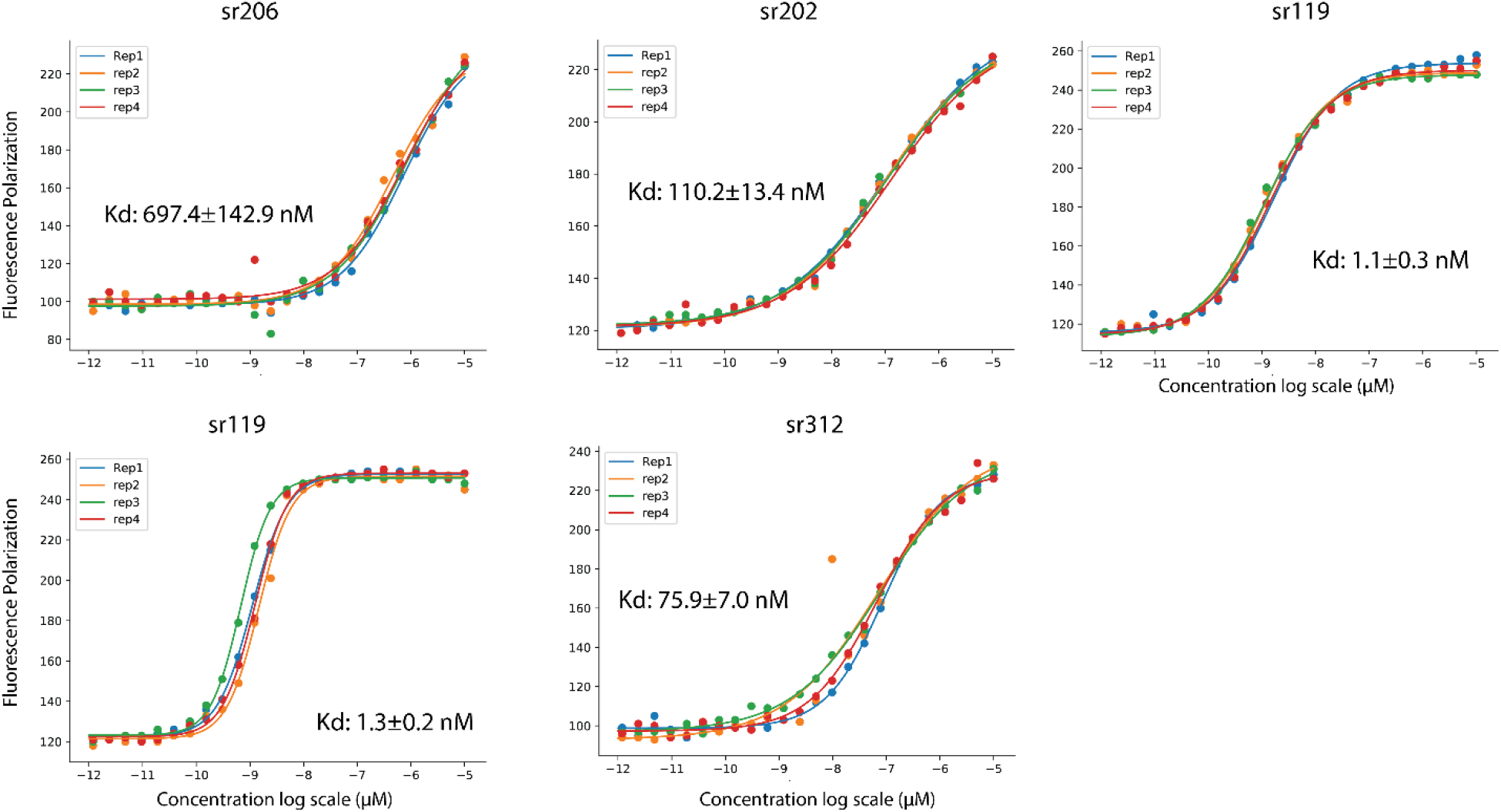
Fluorescence Polarization quantification of affinity of rings. Protein is titrated down in 2-fold dilutions from 10 μM across 24 wells at a constant concentration of 1 nM TAMRA-labelled peptide. Estimates of dissociation constant (Kd) and standard error from four replicate curves are printed in the bottom right.

**Extended Data Figure 9.**
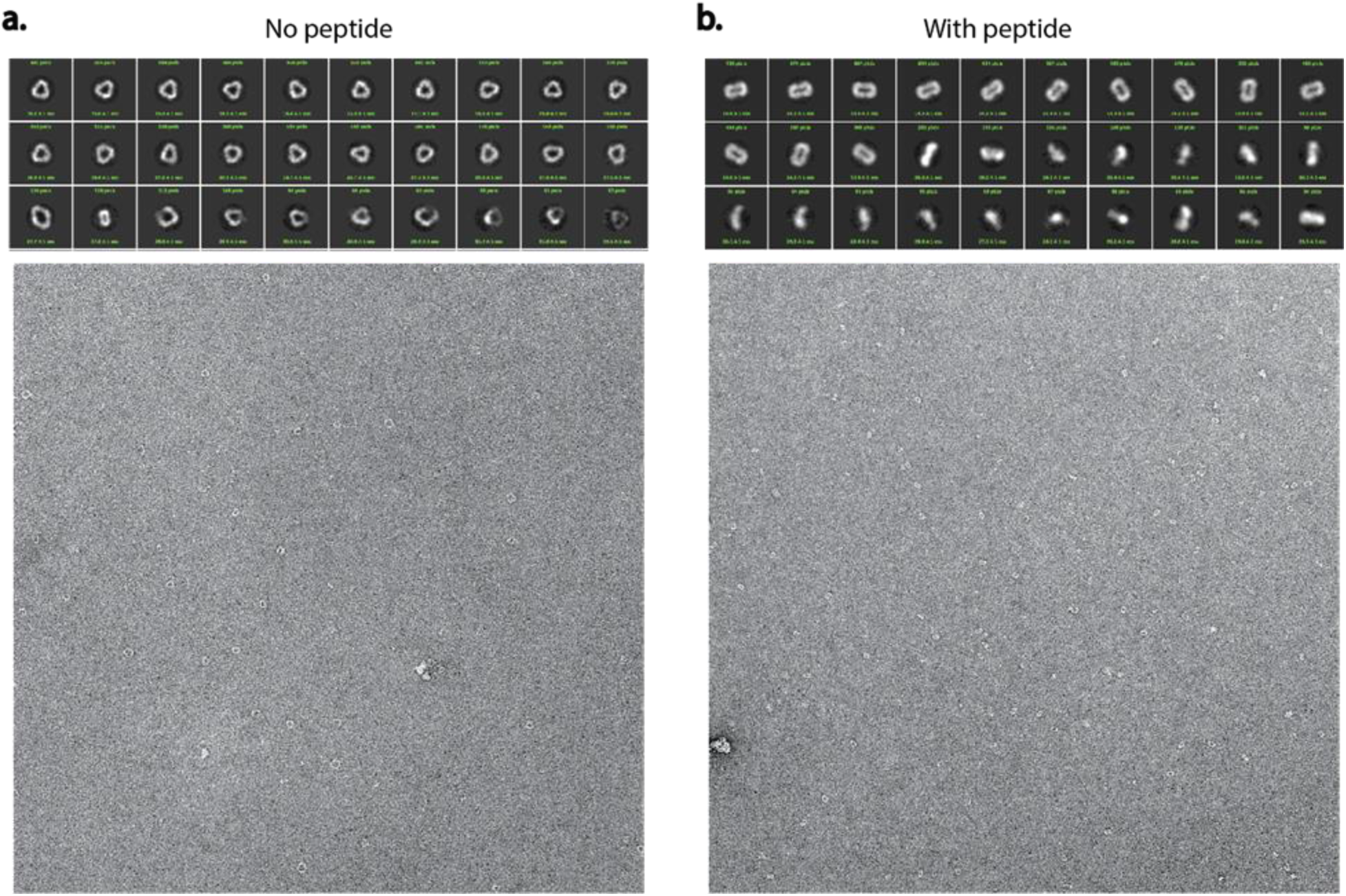
Negative stain EM for sr119. 30 most populated 2D classes (top) and representative micrographs (bottom) in the absence (**A**) and the presence (**B**) of peptide.

**Extended Data Figure 10.**
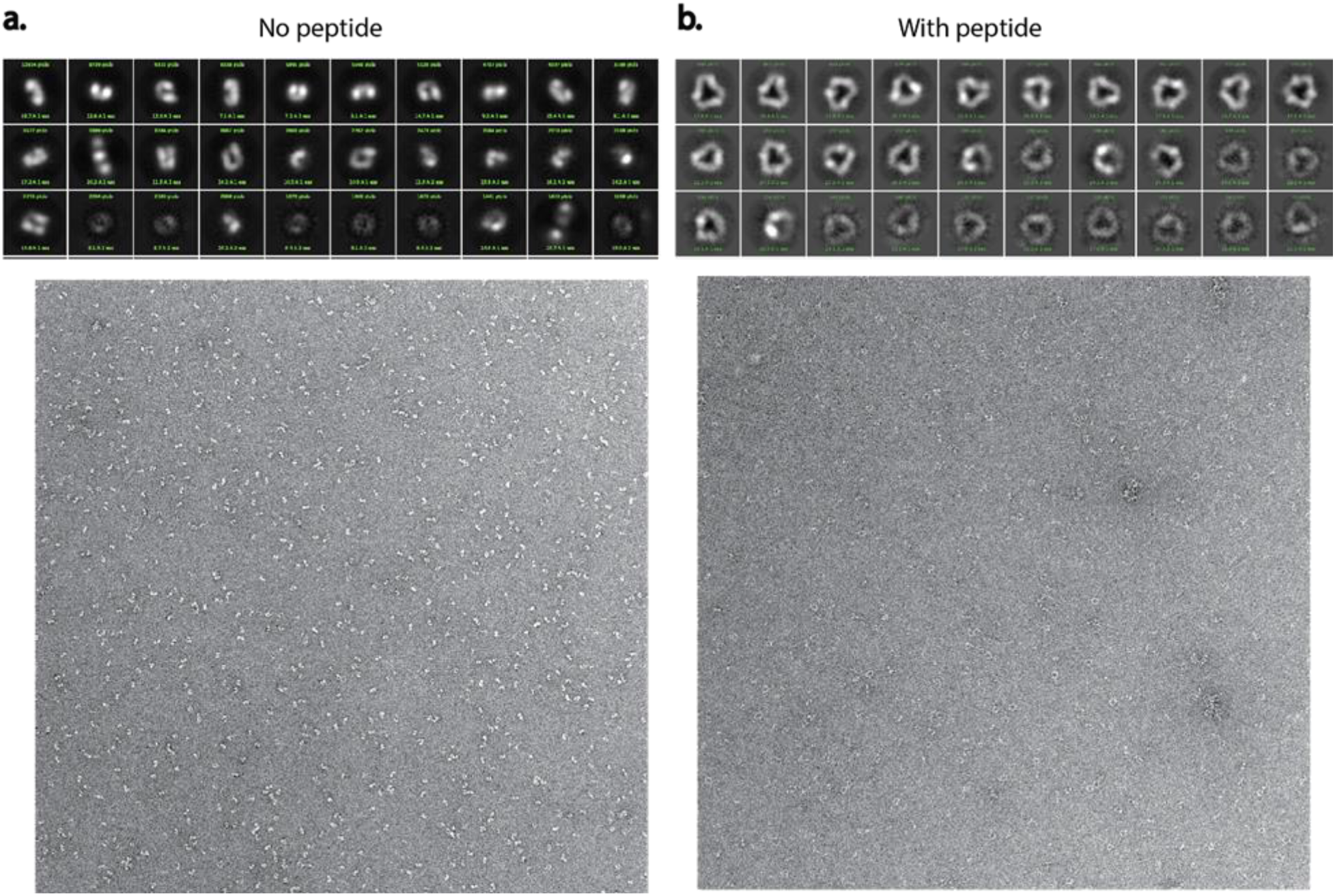
Negative stain EM for sr202. 30 most populated 2D classes (top) and representative micrographs (bottom) in the absence (**A**) and the presence (**B**) of peptide.

**Extended Data Figure 11.**
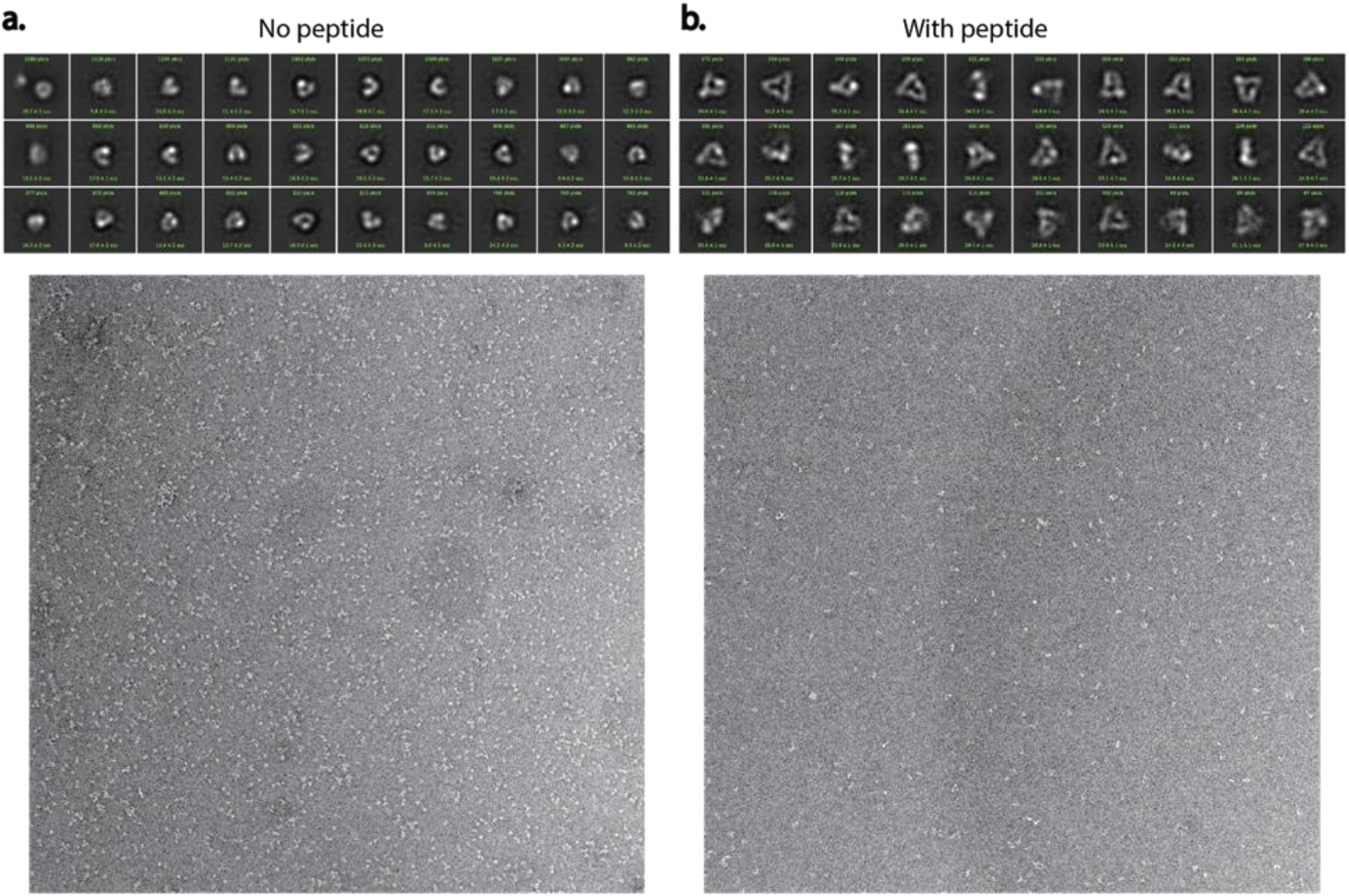
Negative stain EM for sr306. 30 most populated 2D classes (top) and representative micrographs (bottom) in the absence (**A**) and the presence (**B**) of peptide.

**Extended Data Figure 12.**
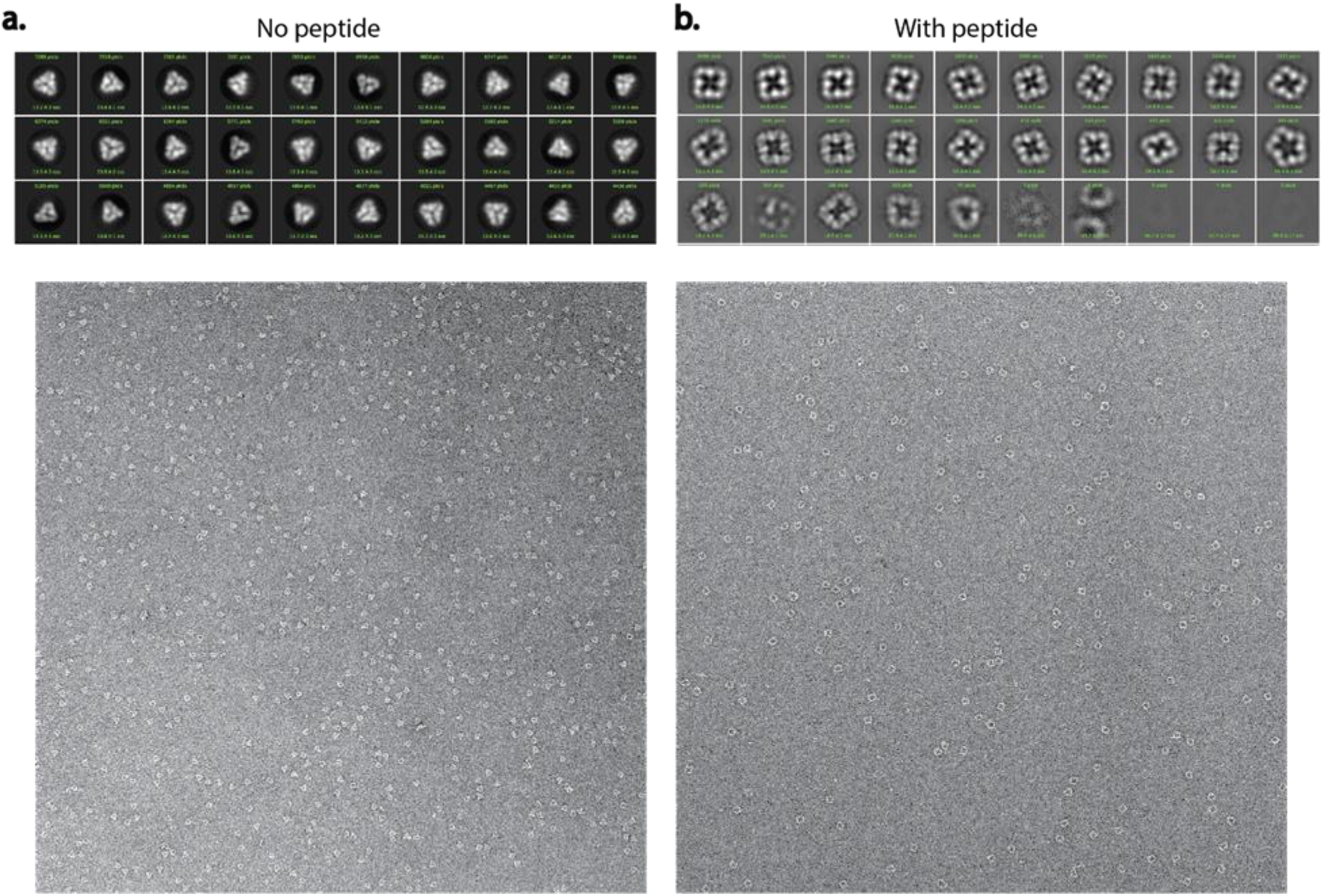
Negative stain EM for sr312. 30 most populated 2D classes (top) and representative micrographs (bottom) in the absence (**A**) and the presence (**B**) of peptide.

**Extended Data Figure 13.**
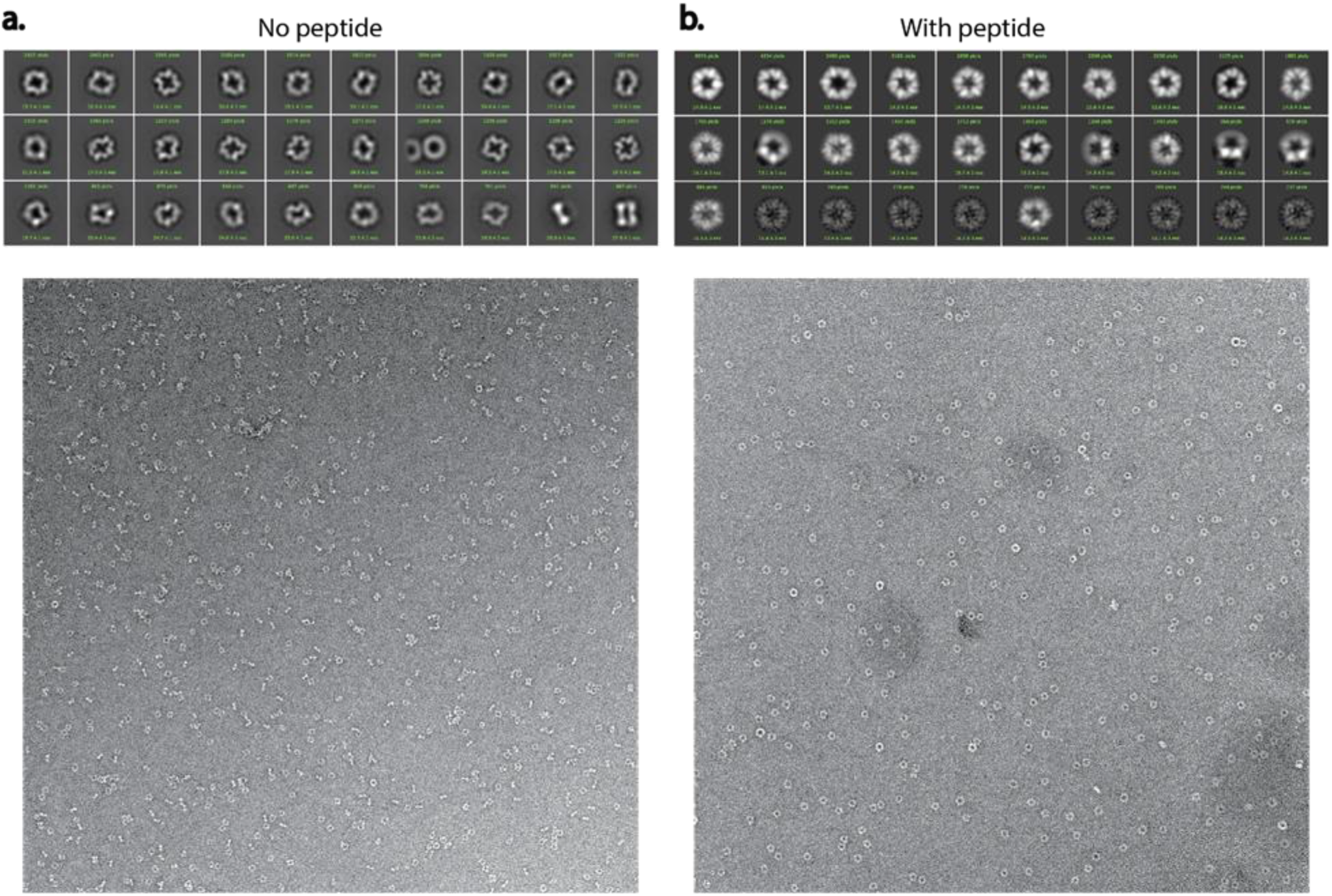
Negative stain EM for sr322. 30 most populated 2D classes (top) and representative micrographs (bottom) in the absence (**A**) and the presence (**B**) of peptide.

**Extended Data Figure 14:**
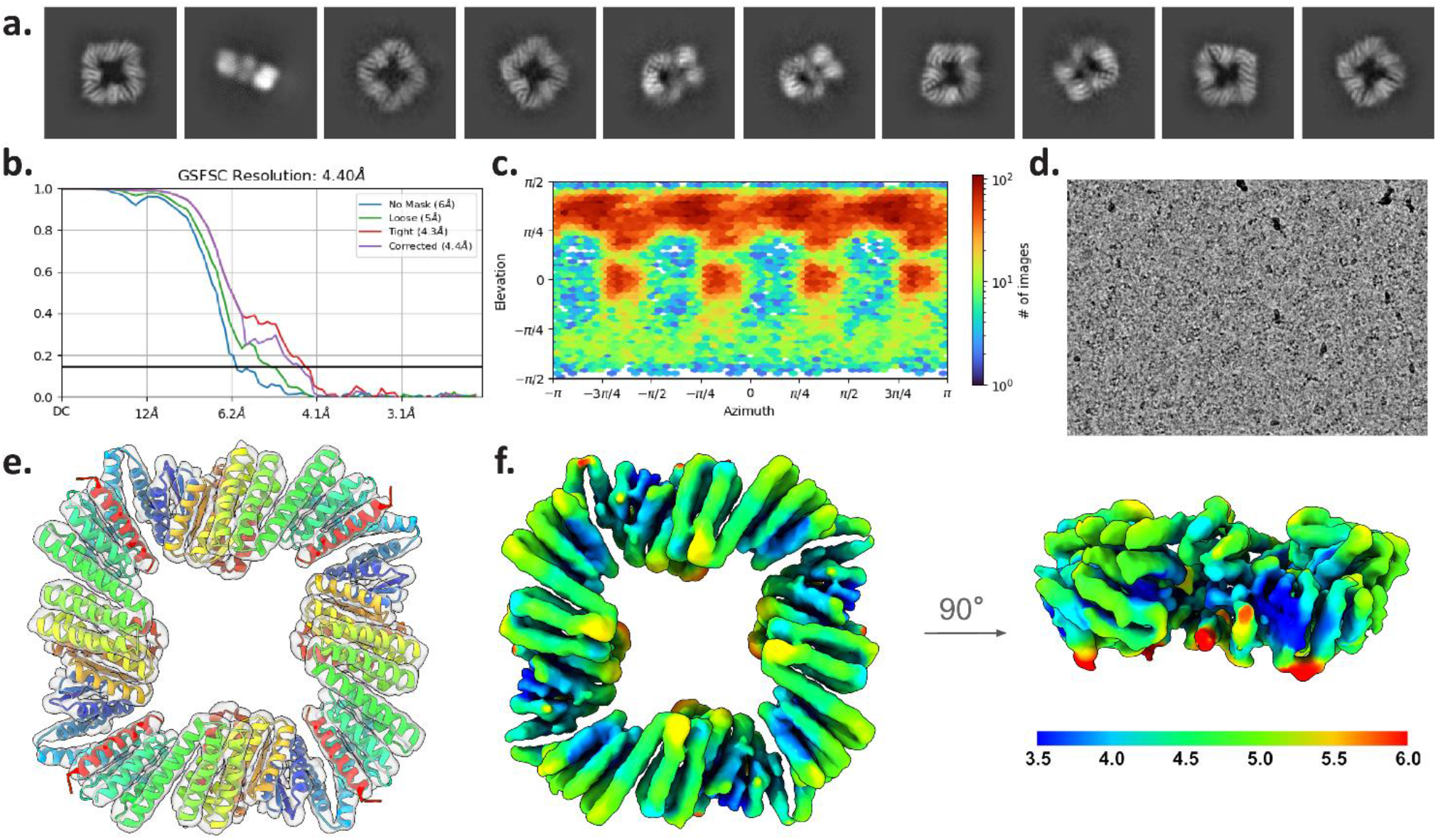
CryoEM data for sr312 in the peptide-bound state. **A**, Representative 2D class averages **B**, Global FSC. **C**, Orientational distribution plot demonstrating full angular sampling **D**, Representative micrograph **E,** Refined 4.40Å structure fit into density in rainbow, effector peptide in red **F,** CryoEM local resolution map in Å

**Extended Data Figure 15:**
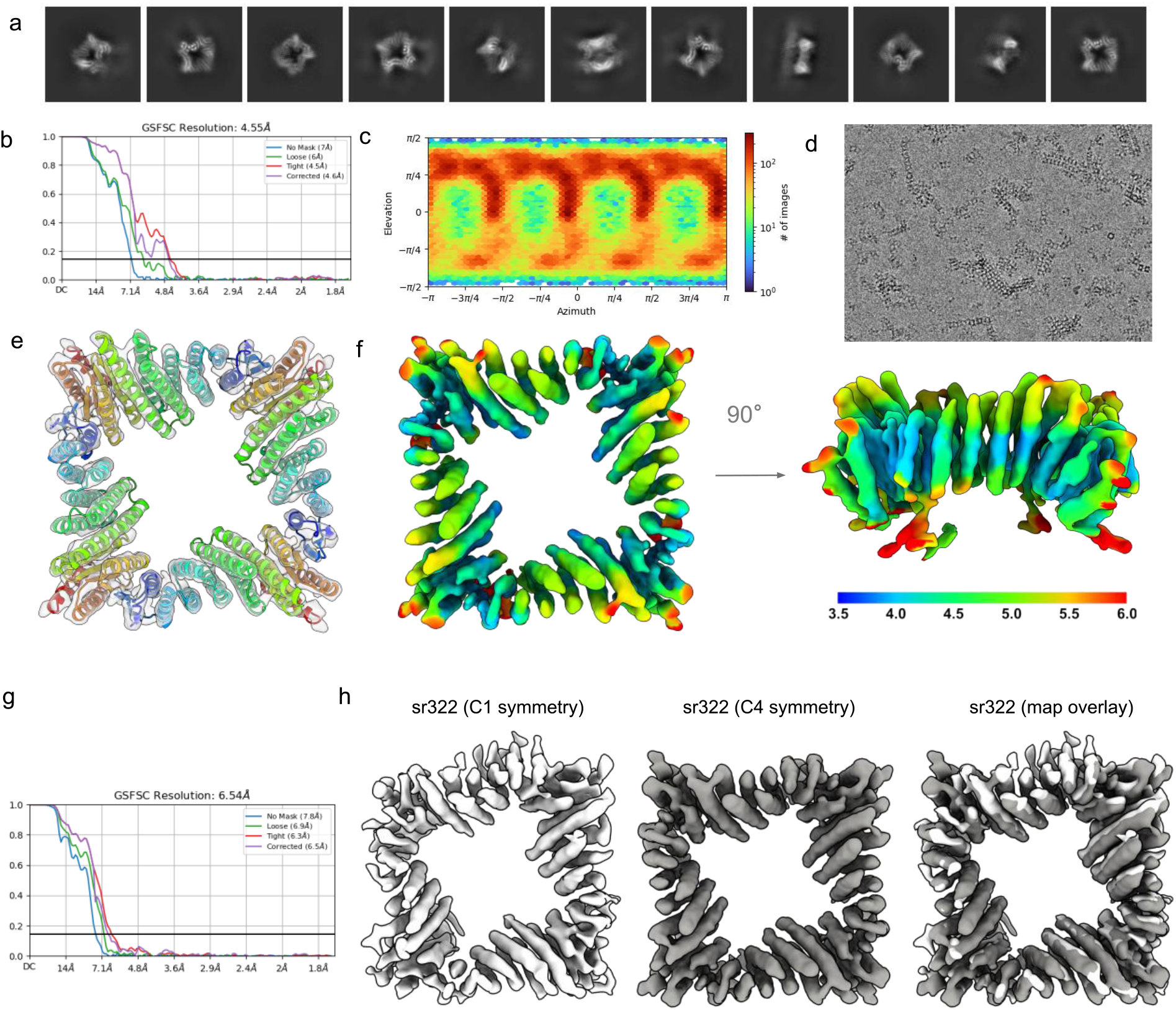
CryoEM data for sr322 in the apo ‘X’ state. **A**, Representative 2D class averages **B**, Global FSC. **C**, Orientational distribution plot demonstrating full angular sampling **D**, Representative micrograph **E,** Refined 4.62Å structure fit into density in rainbow **F,** CryoEM local resolution map in Å. **G**, Global FSC of C1 refinement **H**, Comparison of C1 and C4 refined maps of sr322.

**Extended Data Figure 16.**
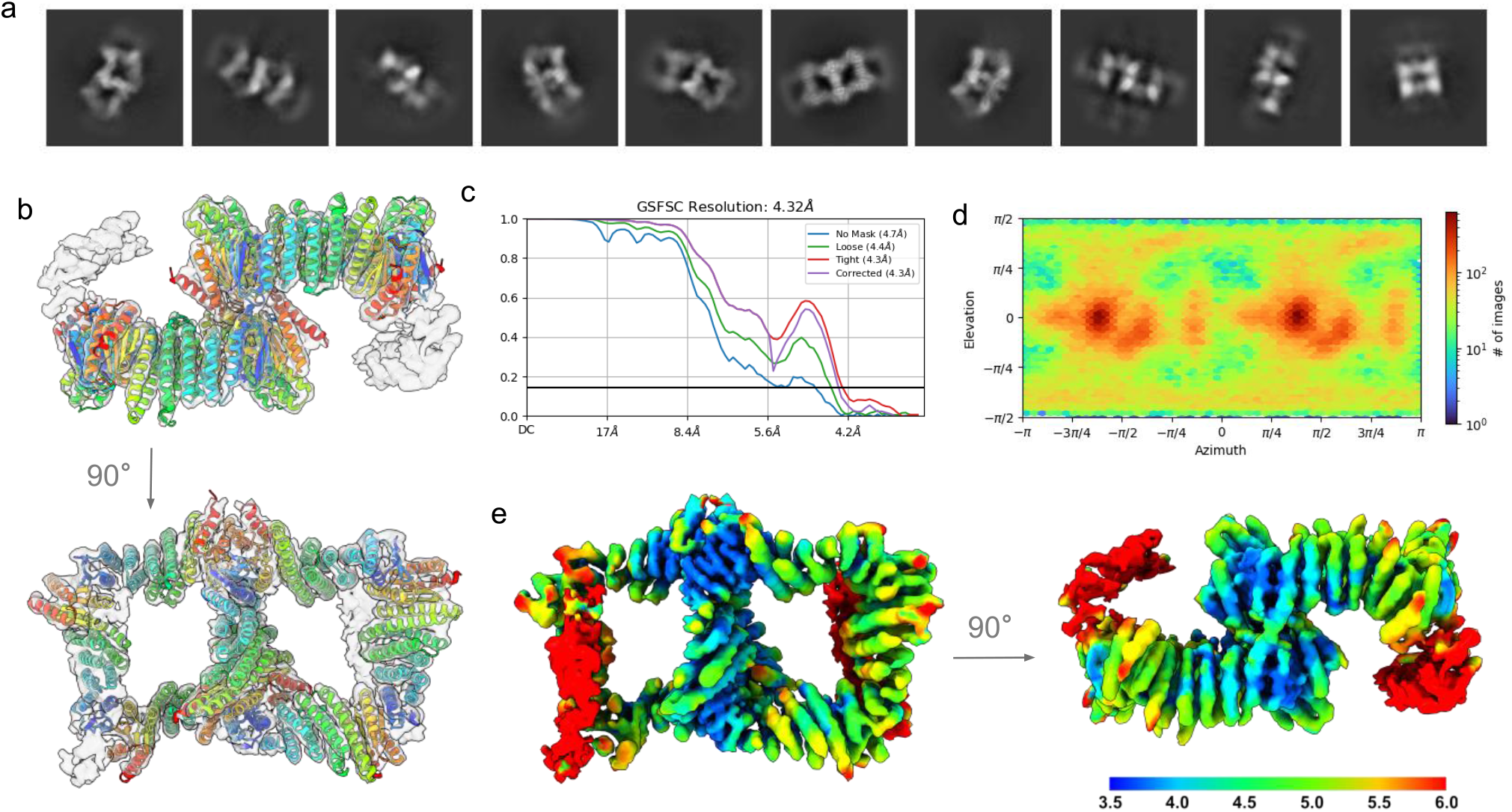
sr322 CryoEM structure data for clustered species. **A**, Representative 2D class averages **B**, Refined 4.32Å structure fit into density in rainbow **C**, Global FSC. **D**, Orientational distribution plot demonstrating full angular sampling **E**, CryoEM local resolution map in Å.

**Extended Data Figure 17.**
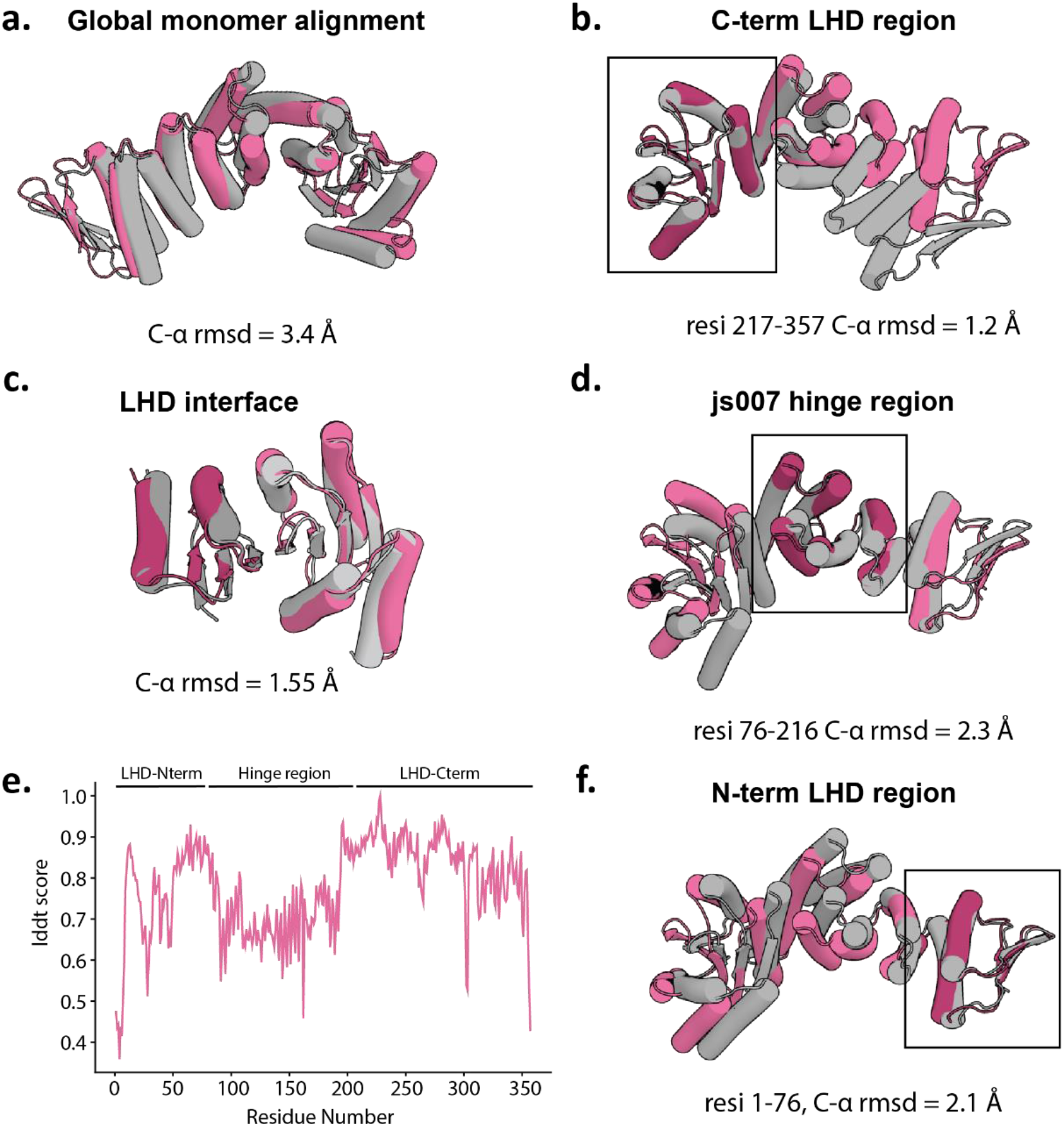
Comparison of sr322 CryoEM structure to AF2 X-state prediction. Alignments shown for Cryo-EM structure (pink) to AF2 model (gray) for **A.** the entire monomer. **B.** the C-terminal LHD domain **C.** the LHD278 interface **D.** the hinge module **F.** the N-terminal LHD domain, with reported RMSD obtained in pyrosetta. **E.** C-α lDDt score for all residues in a comparison between the empirical Cryo-EM monomer and the modelled structure of sr322 in AF2.

**Extended Data Figure 18.**
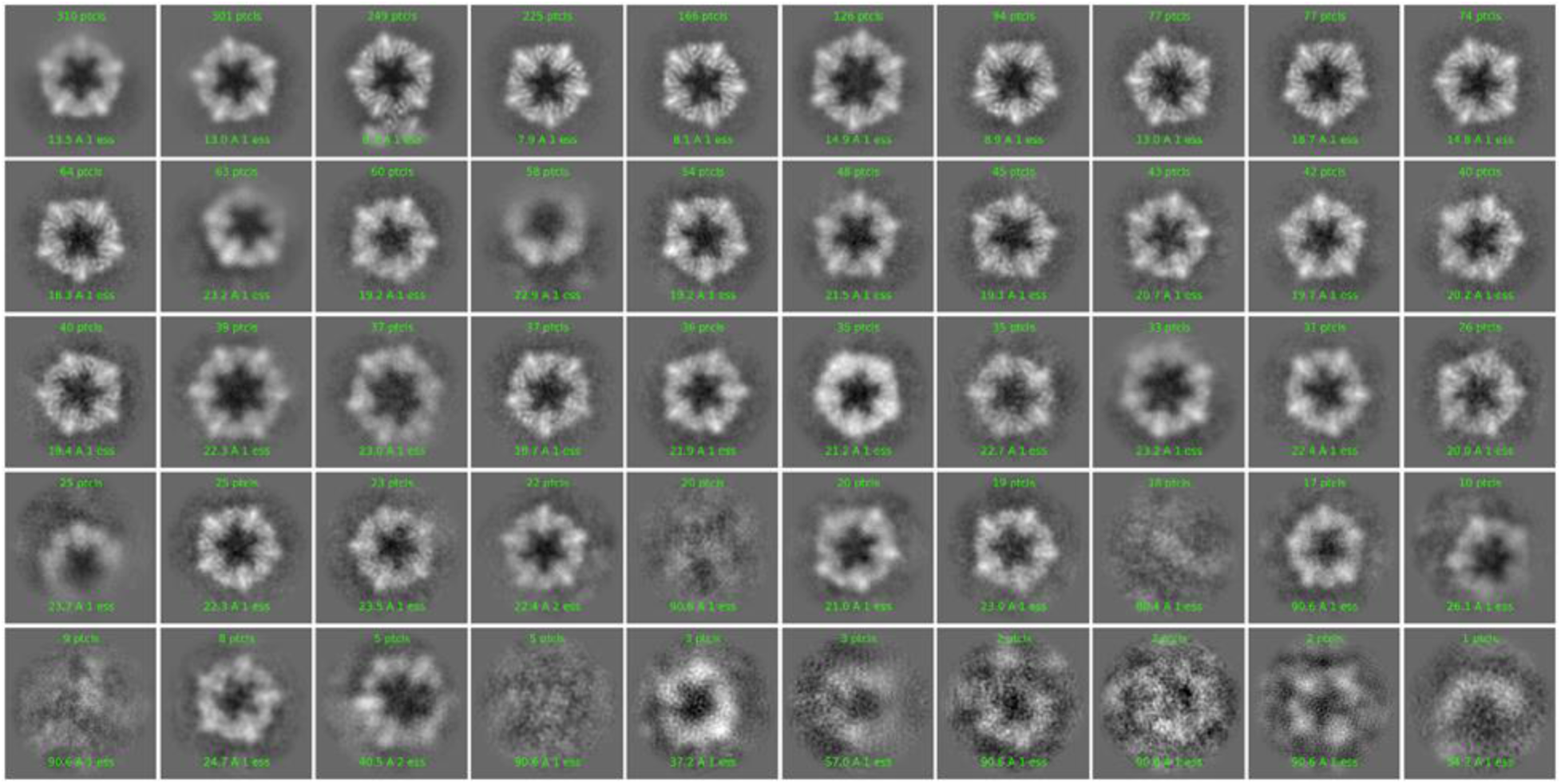
2D-class averages obtained from Cryo-EM on Y-state sr322 with 20-fold excess of peptide.

**Extended Data Figure 19.**
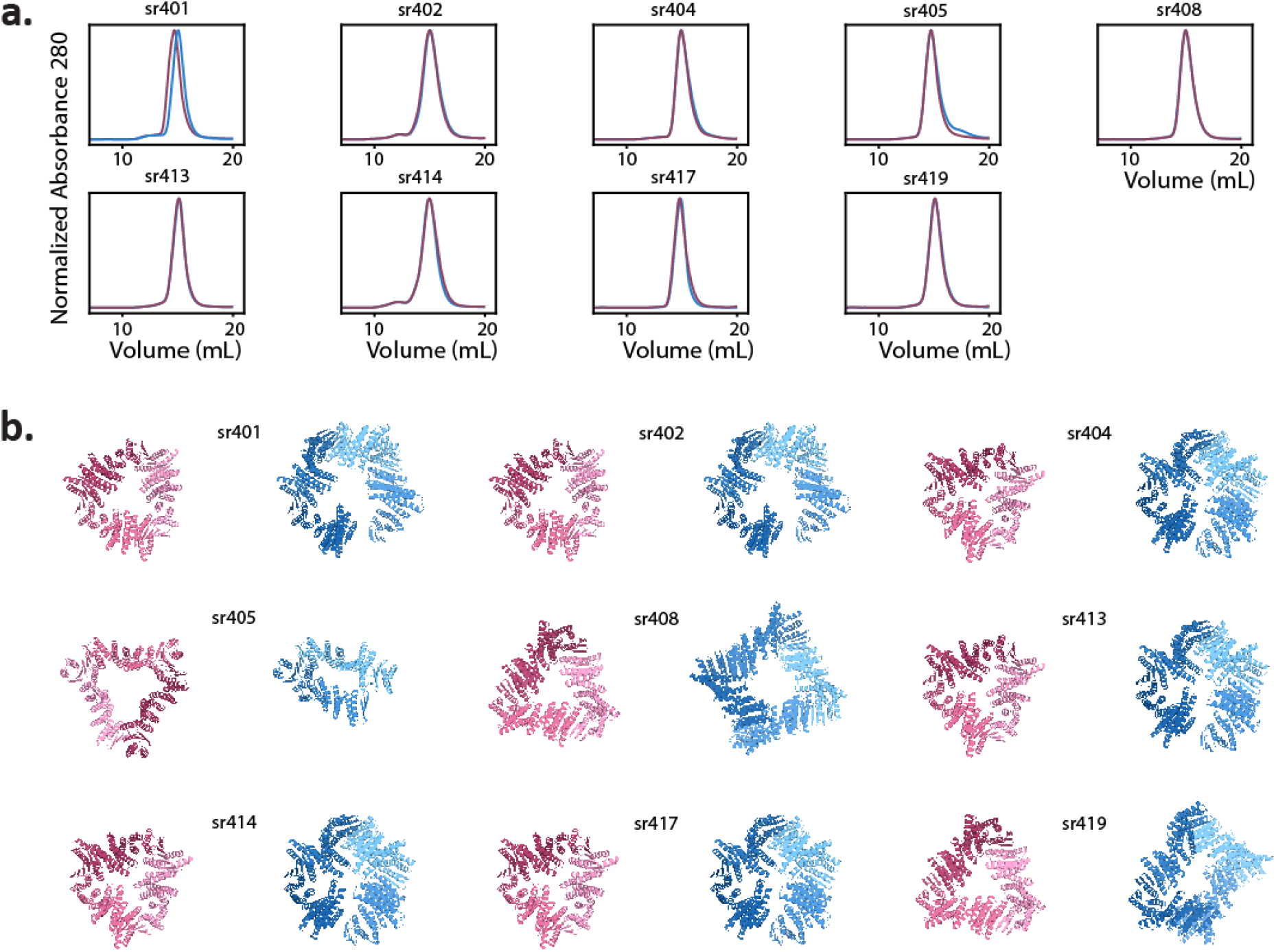
Characterization of static rings. **A.** UVA280 SEC traces for each soluble design, showing comparison between 5 μM protein in the presence (blue) and absence (pink) of 10 μM effector peptide. **B.** Structures of soluble static designs in the Xm (pink) and Yn (blue) states.

**Extended Data Figure 20.**
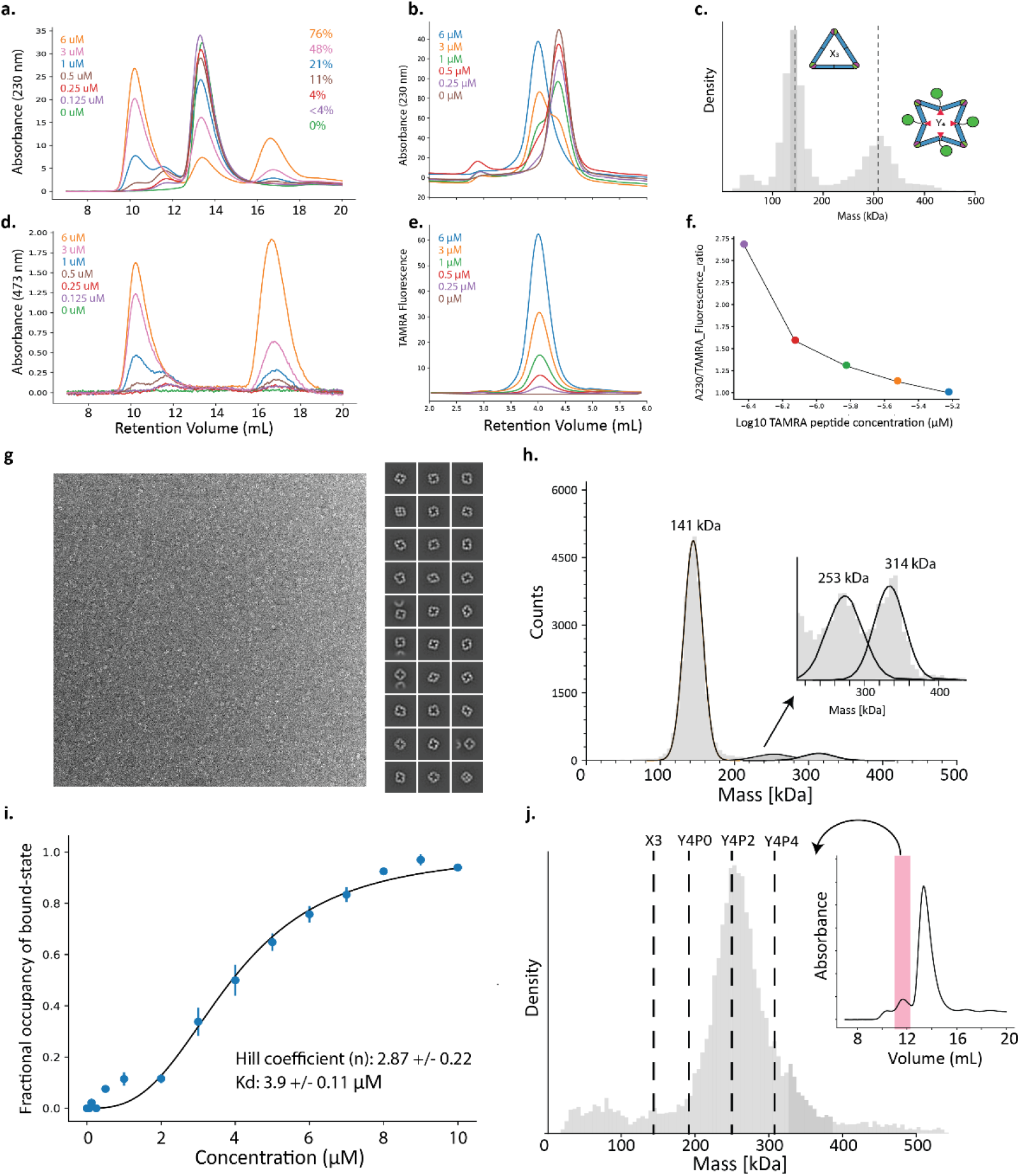
Characterization of cooperativity. **A,D.** SEC titration series of 3 μM protein against variable concentrations GFP-tagged peptide, with A230 and A473 absorbance shown in top and bottom respectively. The concentration of added peptide, and the molar fraction of bound hinges is indicated on the right. **B,E.** HPLC SEC titration series of 3 μM protein against variable TAMRA-labeled peptide concentrations, with A230 and TAMRA fluorescence (Excitation: 530 Emission: 590) plotted in top and bottom respectively. **C.** Mass photometry data showing clear separation between the X3 and Y4P4 states at 3 μM protein and 1 μM peptide. **F.** Ratio of TAMRA signal to A230 signal across titration series shown in **B. G.** nsEM micrograph (left) and 2D-class averages (right) for sr312_Y_staple. **H.** Mass distribution for mixture of 3 μM sr312 and 0.5 μM GFP_tagged 221, with 253 kDa species intermediate in mass between Y4P4 and X3 shown in inset. **I.** Fractional saturation curve of sr312 across a titration series of GFP-tagged 221B peptide, with bound fractions estimated from mass photometry and estimated hill coefficient (n) and Kd listed. **J.** Mass distribution of intermediate Y2P2 species isolated from SEC at 11% saturation (inset, with fraction used highlighted in red). Vertical dashed lines denote expected masses for labeled species.

**Extended Data Figure 21.**
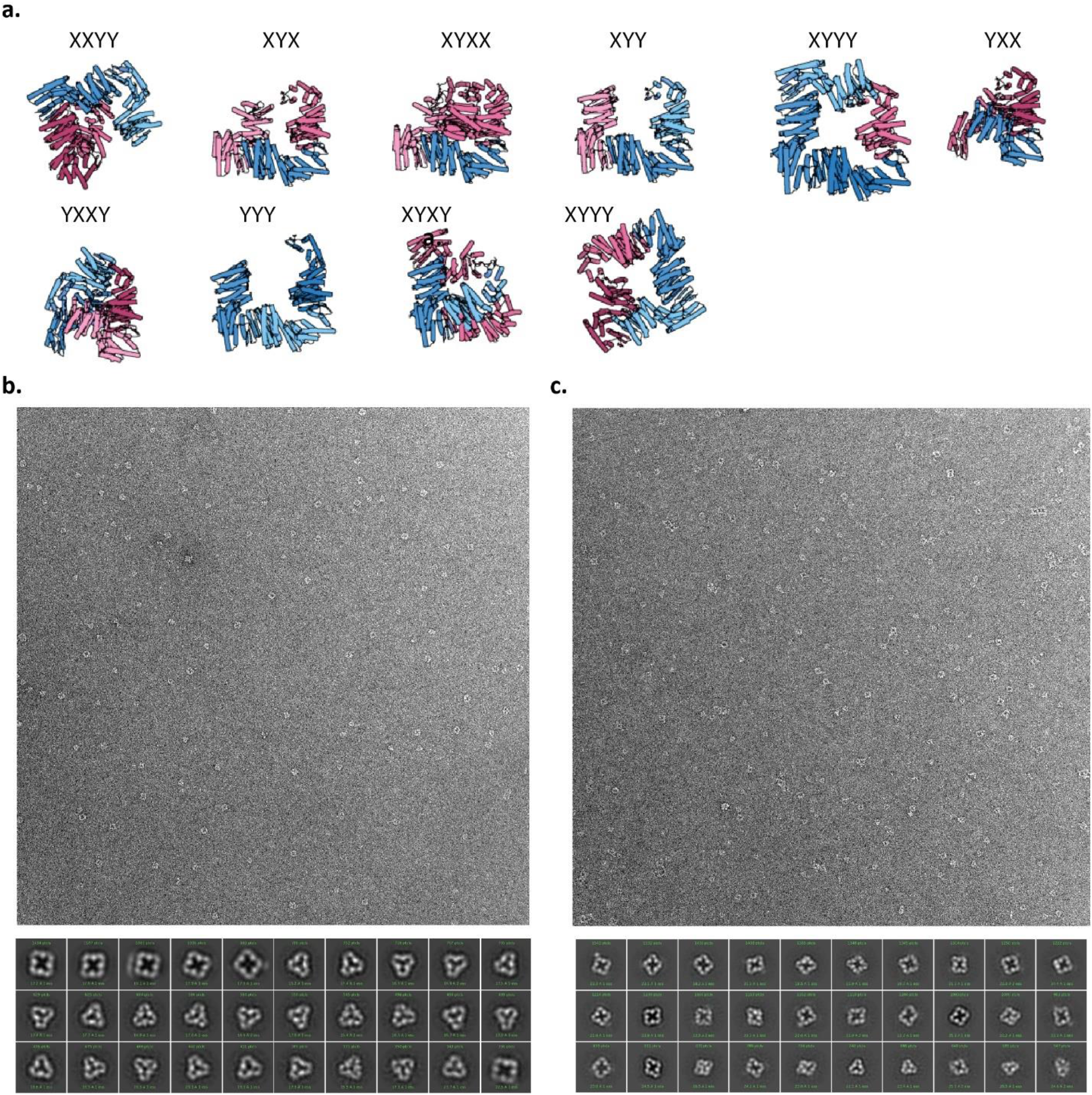
a. Range of partially bound XY oligomeric states of sr312 that would be expected to be populated in a non-cooperative scheme of peptide-binding, or that are transiently populated in the KNF model. X-state chains (pink) and Y-state chains (blue) are indicated for each modeled oligomer, with the number of chains and their states labeled. **B.** nsEM micrograph (top) and most populated 2D class averages (bottom) for the 50% ligated sample of sr312 shown in Fig 4e**. C.** nsEM micrograph (top) and most populated 2D class averages (bottom) for a partially ligated species isolated via SEC from the ∼10% ligated sample of sr312 shown in **Efig 20j**.

**Extended Data Figure 22.**
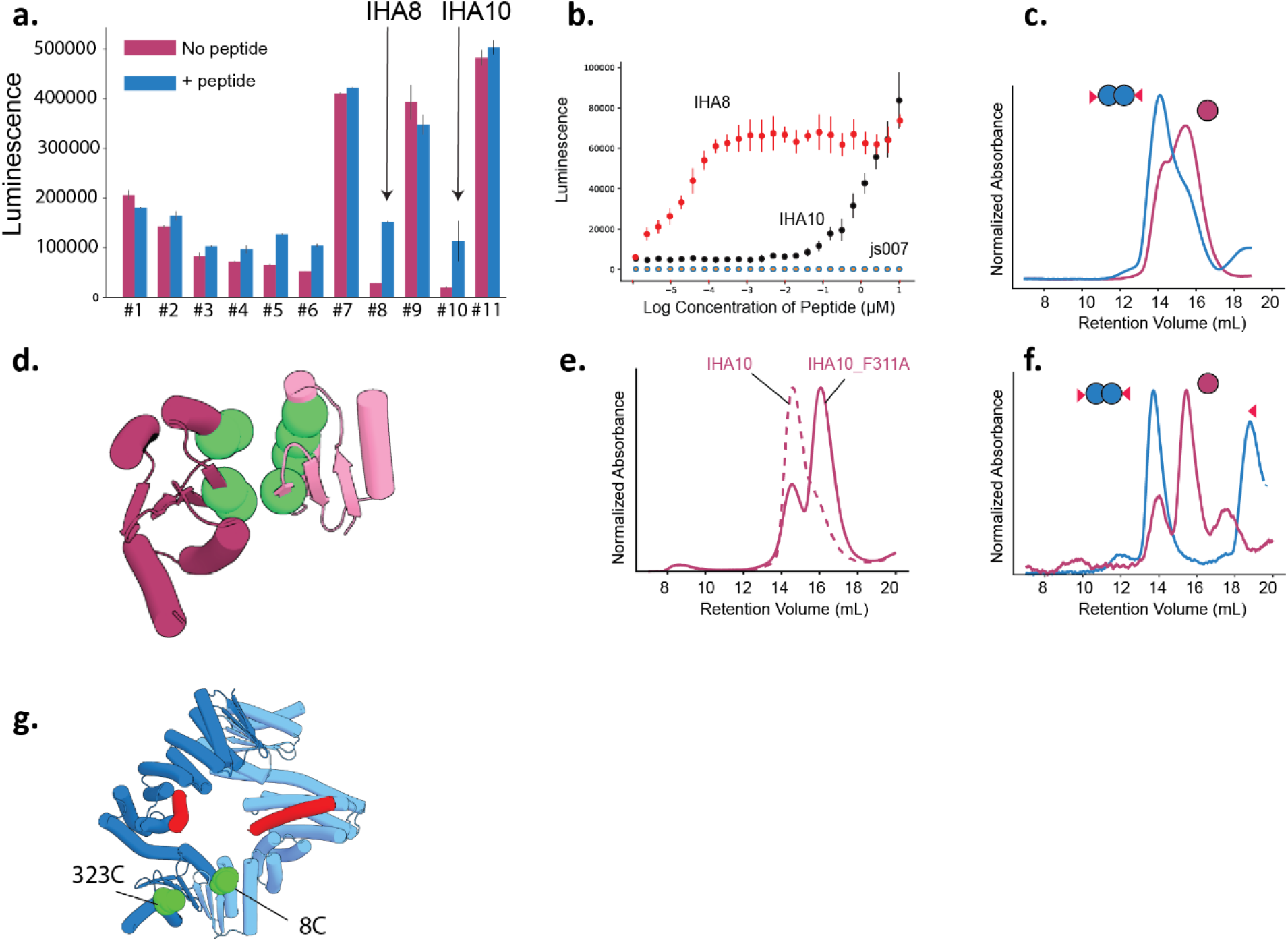
Characterization of inducible homodimer systems. **A.** NanoBit luciferase measurements from 11 soluble designs at 5 nm of each tagged-luciferase construct shown in the absence (pink) and presence (blue) of 5 μM cognate peptide. **B.** Luminescence readings for each inducible dimer design derived from an equimolar mixture of NanoBit luciferase parts tagged to the design, where each component is held constant at 10nM, while the peptide is titrated in two-fold steps down from a maximum concentration of 10μM. Tested designs include IHA10 (black), IHA8 (red) and a lg and smBit tagged hinge construct, js007, (gray) that does not assemble into oligomers as a control for off-target luciferase activity. **C.** SEC on IHA10 at 1 μM in the presence and absence of 10 μM peptide **D.** Interfacial sites (green spheres) that were mutated to alanine in IHA10. The LHD regions of the two opposing chains are shown in different pink shades. **E.** SEC comparison of IHA10 (dashed pink) and the alanine mutant IHA10_F311A (solid pink) at 2 μM**. F.** SEC on 1μM IHA10_F311A in the presence (blue) and absence (pink) of 20 μM peptide. The additional peak in the blue trace corresponds to free, unbound peptide. **G.** Two interface-adjacent sites in IHA10 that were chosen for FRET labelling are shown as green spheres, with Y-state chains (blue) and peptide (red) displayed. 8C and 323C were labeled with AlexaFluor 555 and AlexaFluor 647 respectively.

**Extended Data Figure 23.**
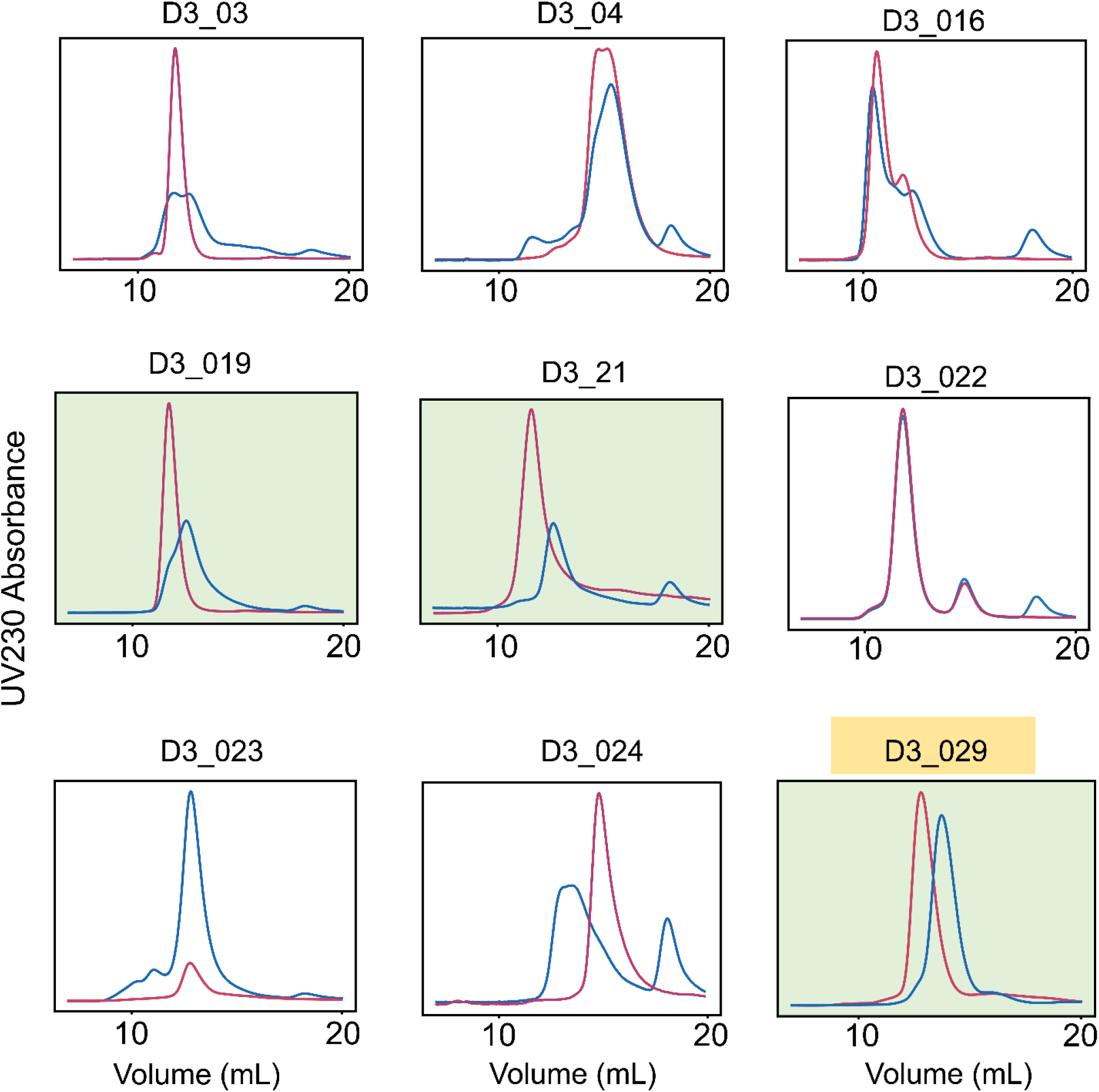
SEC characterization of switchable D3 designs. D3 Designs were tested for peptide-responsiveness at a monomer concentration of 5 μM. UVA230 SEC elution traces in the absence of effector (pink) and in the presence of 10 μM effector peptide cs221B (blue) are shown. Designs with a green background showed a rightward peak shift in the presence of the effector peptide. The additional peak in the blue traces at ∼18 ml corresponds to excess effector peptide.

**Extended Data Figure 24.**
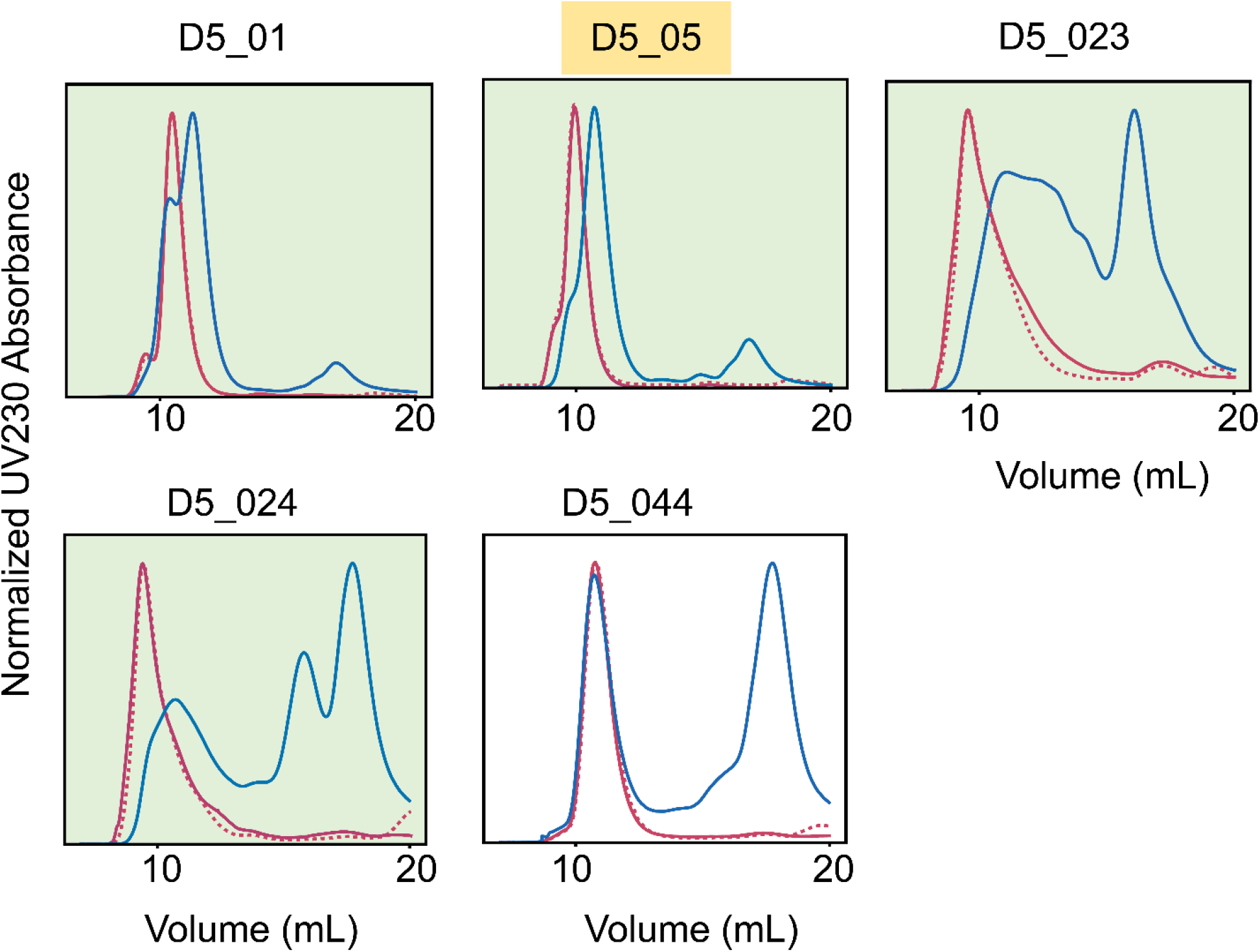
SEC characterization of switchable D5 designs. D5 Designs were tested for peptide-responsiveness at a monomer concentration of 5 μM. UVA230 SEC elution traces in the absence of effector (solid pink) and in presence of 10 μM effector peptide cs221B (dashed pink) or 10 μM effector protein 3hb21 (blue) are displayed. Designs with a green background showed a rightward peak shift in the presence of the effector protein. The additional peak in the blue traces at ∼18 ml corresponds to excess effector protein.

**Extended Data Figure 25.**
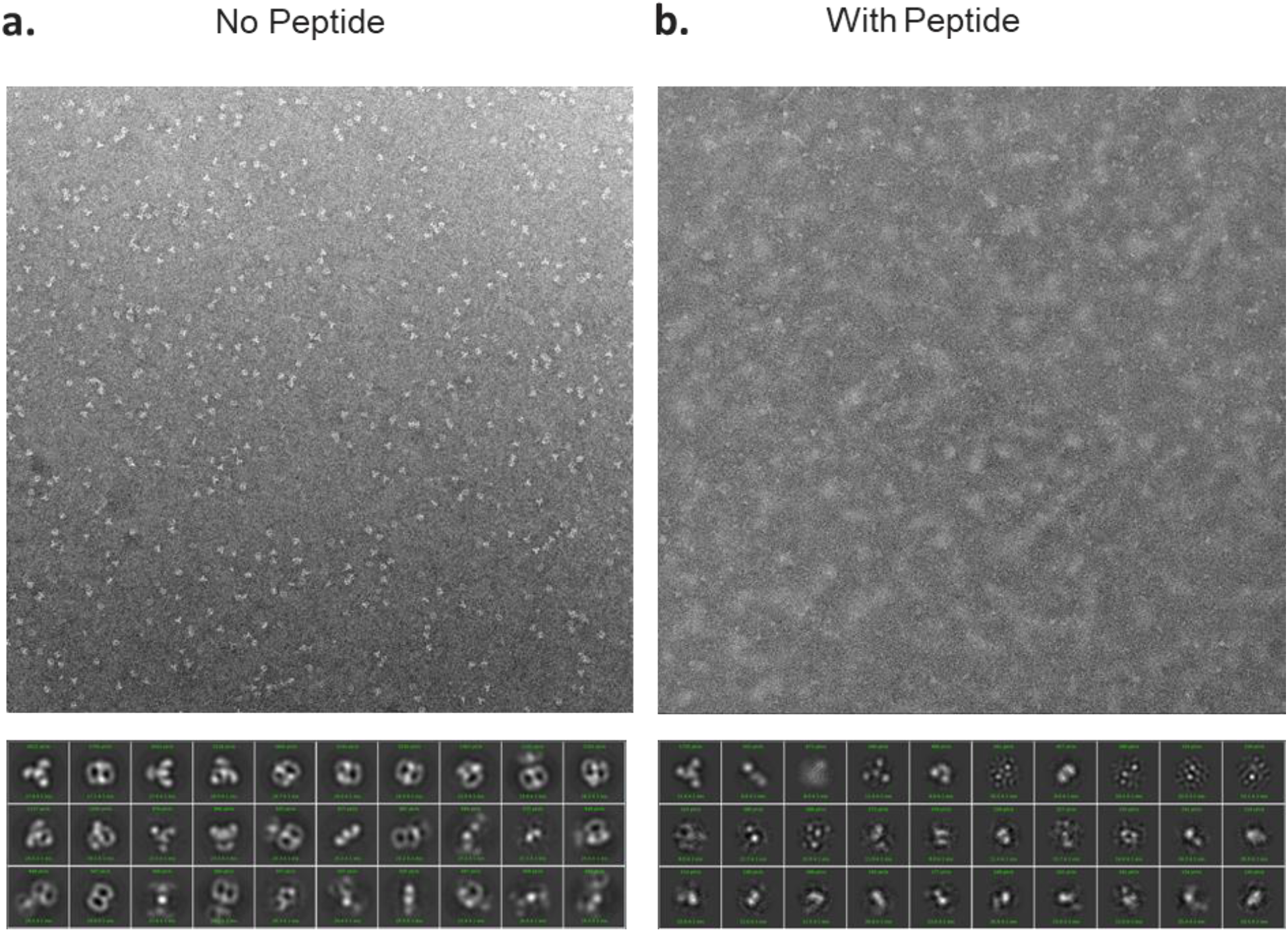
Negative stain EM for D3_29. Representative micrographs (top) and 30 most populated 2D class averages (bottom) in the absence (**A**) and the presence (**B**) of peptide.

**Extended Data Figure 26.**
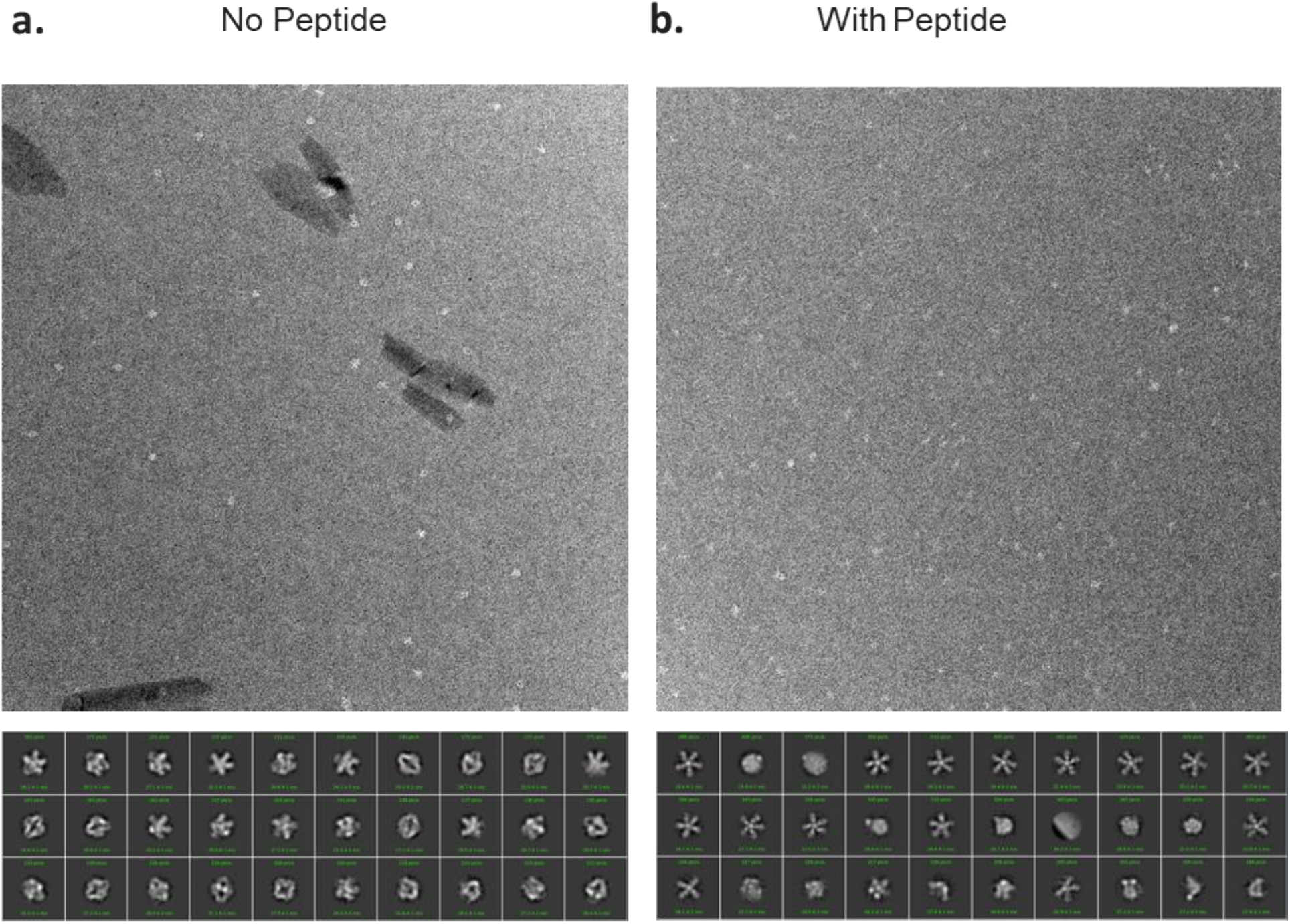
Negative stain EM for D5_05. Representative micrographs (top) and 30 most populated 2D class averages (bottom) in the absence (**A**) and the presence (**B**) of effector protein.

**SUPPLEMENTARY TABLE I.**
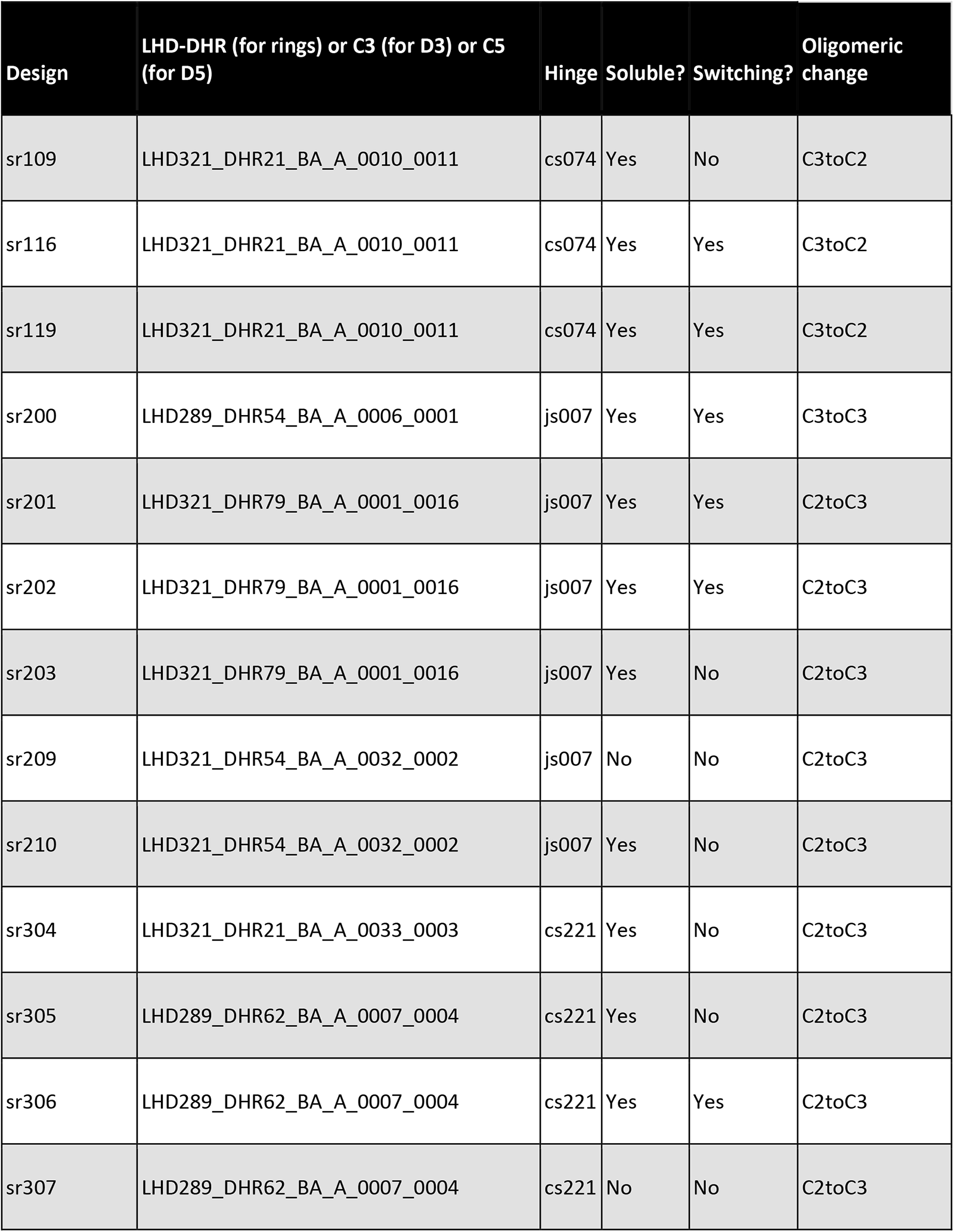
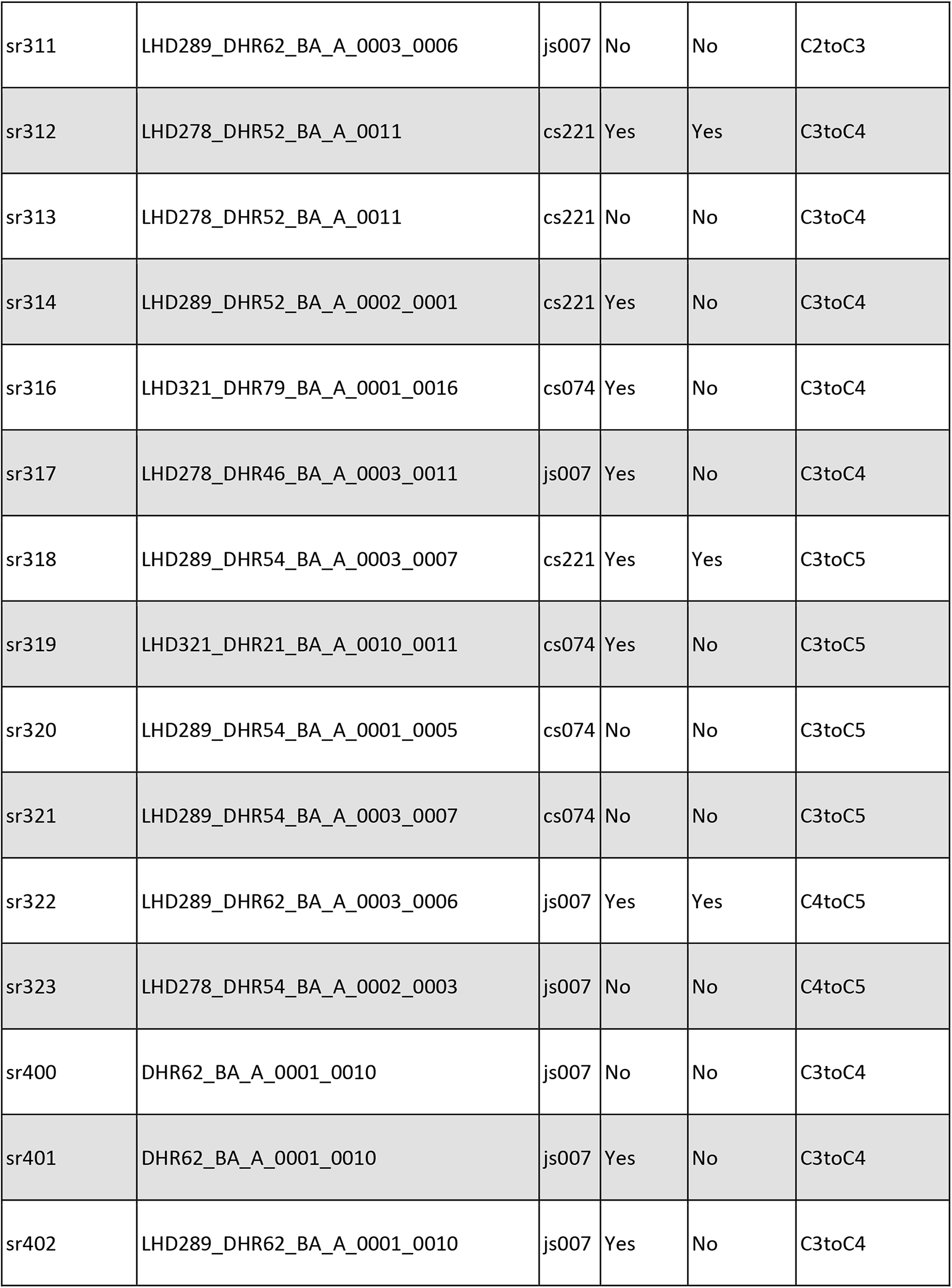
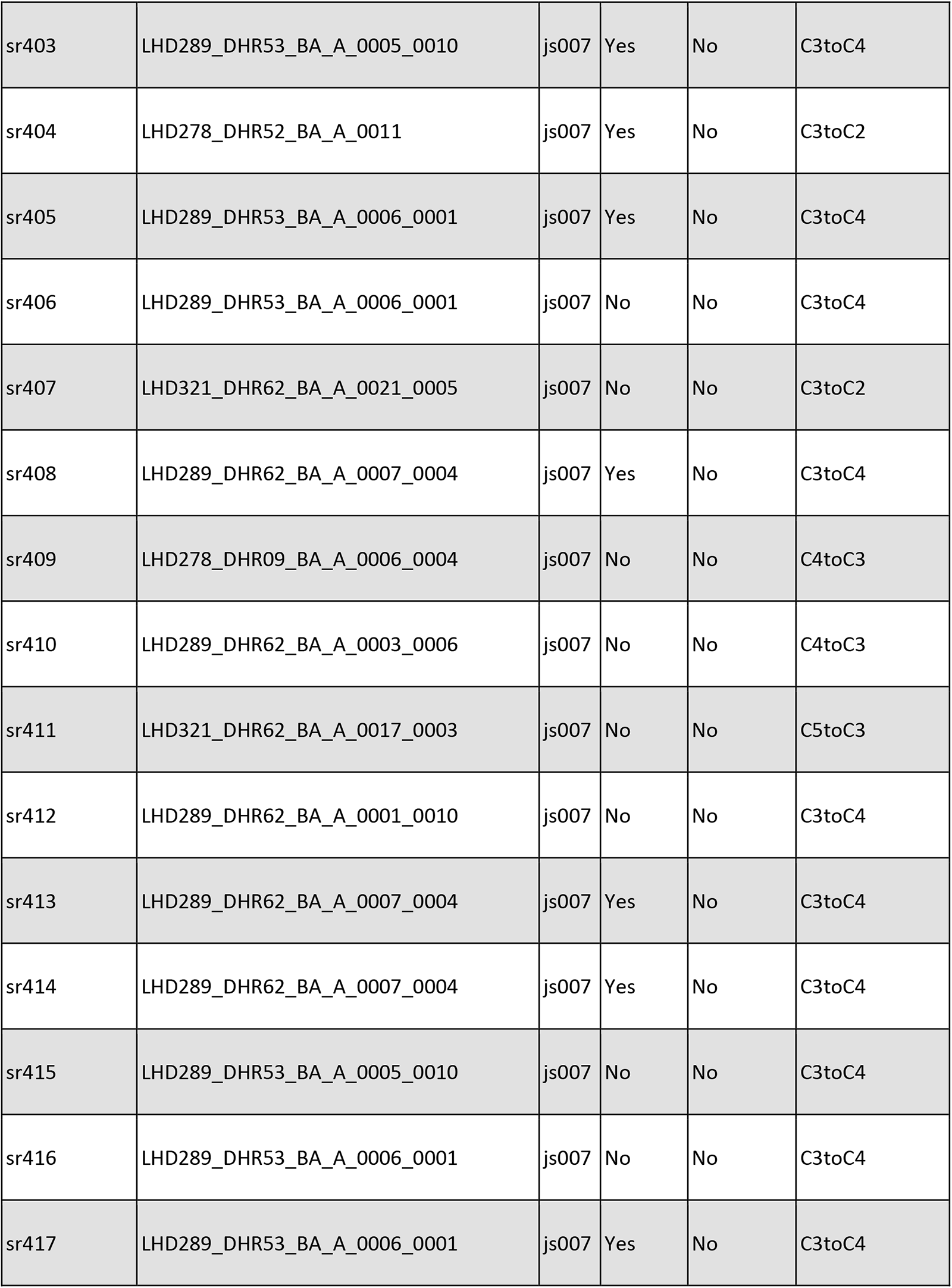
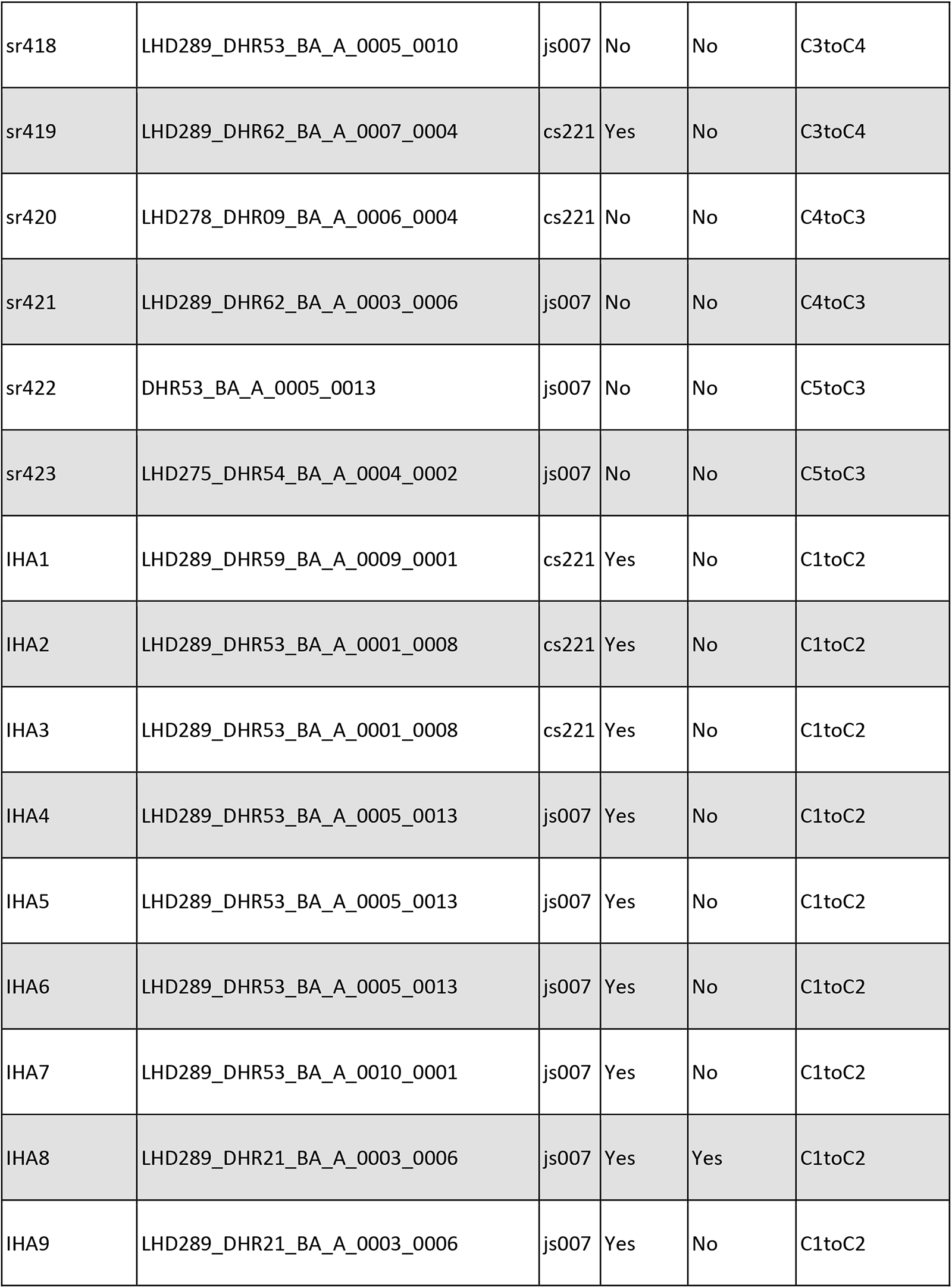
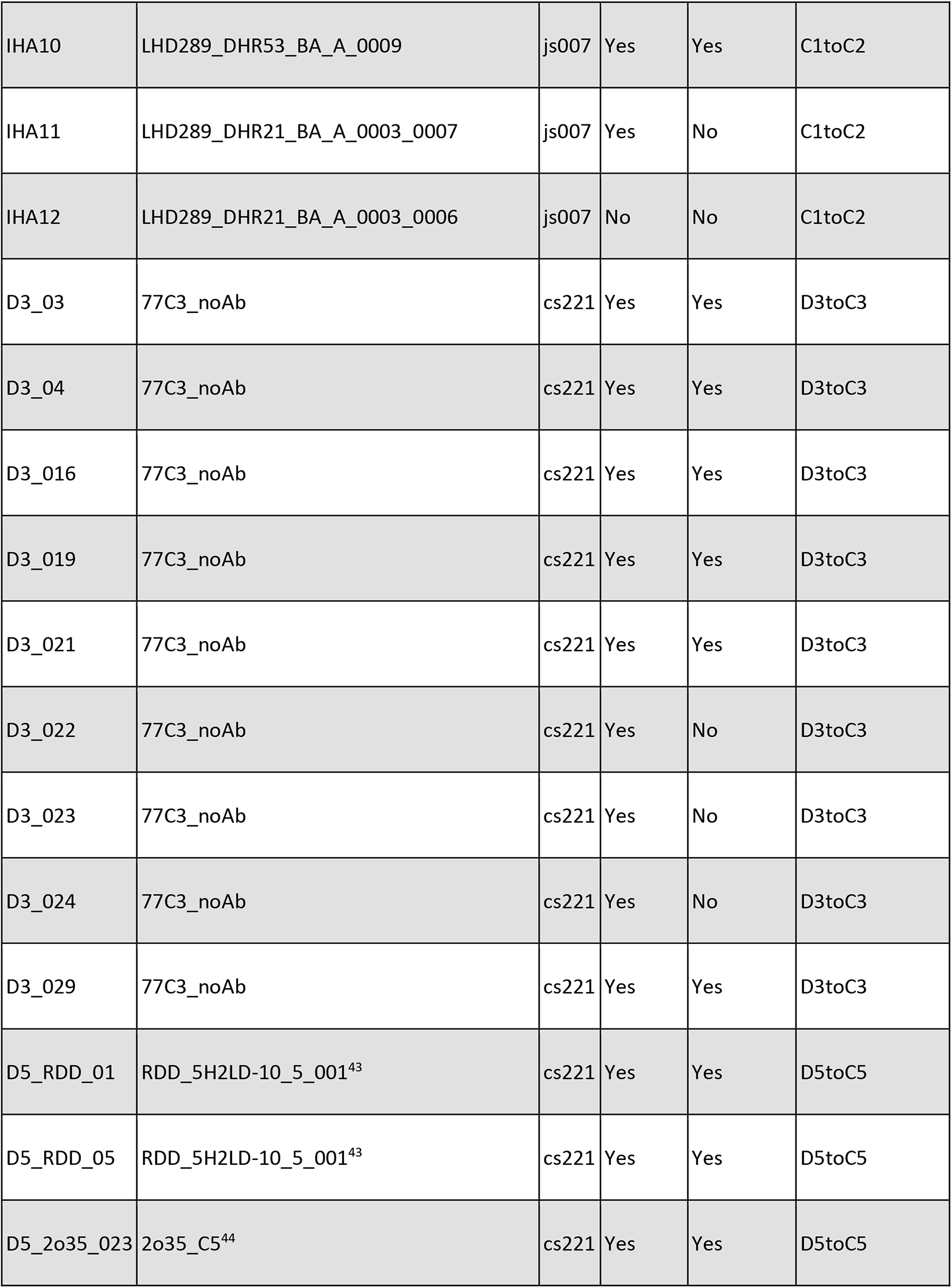
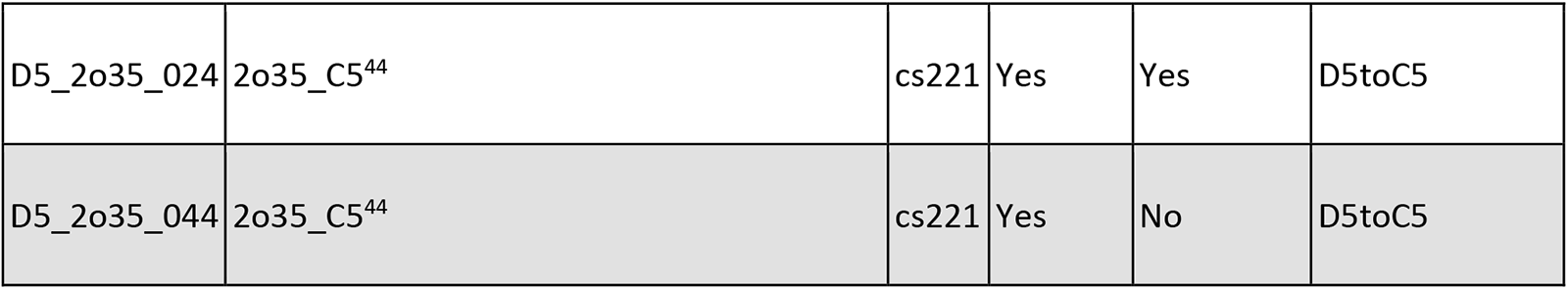

## SUPPLEMENTARY NOTE 1

Under the concentration conditions of cryo-EM (>40 µM), we also observed an additional higher-order assembly state that is not observed under the nanomolar assay conditions of negative stain EM or mass photometry. In this state, closed C4 rings of sr322 intricately stack into fibers. A 4.32 Å cryo-EM structure of this clustered species is shown in **Efig 16**. Additionally, we also visualized the holo C5 sr322 assembly in the bound Y-state upon addition of the effector peptide. Unfortunately, the determination of the 3D structure for this state faced challenges due to a notable orientational bias. Nevertheless, the high-resolution 2D class averages of the Y-state highlight a C5 ring that matches the Y-state computational design models in both diameter and shape (**Fig 3e**, **Efig. 18**). We also observed a minority population of C6 rings in this context (**Fig 3e**, **Efig. 18**).

## References

1. Link H, Kochanowski K, Sauer U. Systematic identification of allosteric protein-metabolite interactions that control enzyme activity in vivo. Nat Biotechnol. 2013;31(4). doi:10.1038/nbt.2489

2. Nussinov R, Tsai CJ, Liu J. Principles of allosteric interactions in cell signaling. J Am Chem Soc. 2014;136(51):17692–17701. doi:10.1021/ja510028c

3. Wodak SJ, Paci E, Dokholyan N V., et al. Allostery in Its Many Disguises: From Theory to Applications. Structure. 2019;27(4):566–578. doi:10.1016/j.str.2019.01.003

4. Fastrez J. Engineering allosteric regulation into biological catalysts. ChemBioChem. 2009;10(18):2824–2835. doi:10.1002/cbic.200900590

5. Pirro F, Schmidt N, Lincoff J, et al. Allosteric cooperation in a de novo-designed two-domain protein. Proc Natl Acad Sci U S A. 2020;117(52):33246–33253. doi:10.1073/pnas.2017062117

6. Reynolds KA, McLaughlin RN, Ranganathan R. Hot spots for allosteric regulation on protein surfaces. Cell. 2011;147(7):1564–1575. doi:10.1016/j.cell.2011.10.049

7. Pincus D, Pandey JP, Feder ZA, Creixell P, Resnekov O, Reynolds KA. Engineering allosteric regulation in protein kinases. Sci Signal. 2018;11(555):1–12. doi:10.1126/scisignal.aar3250

8. Süel GM, Lockless SW, Wall MA, Ranganathan R. Evolutionarily conserved networks of residues mediate allosteric communication in proteins. Nat Struct Biol. 2003;10(1):59–69. doi:10.1038/nsb881

9. Rivalta I, Sultan MM, Lee NS, Manley GA, Loria JP, Batista VS. Allosteric pathways in imidazole glycerol phosphate synthase. Proc Natl Acad Sci U S A. 2012;109(22). doi:10.1073/pnas.1120536109

10. Monod, J., Wyman, J., & Changeux JP. On the nature of allosteric transitions: a plausible model. Jounal Mol Biol. 1965;12(1):88–118.

11. Praetorius F, Leung PJY, Tessmer MH, et al. Design of stimulus-responsive two-state hinge proteins. Science. 2023;381(6659):754–760. doi:10.1126/science.adg7731

12. Sahtoe DD, Praetorius F, Courbet A, et al. Reconfigurable asymmetric protein assemblies through implicit negative design. Science *(80-)*. 2022;375(6578). doi:10.1126/science.abj7662

13. Foley EDB, Kushwah MS, Young G, Kukura P. Mass photometry enables label-free tracking and mass measurement of single proteins on lipid bilayers. Nat Methods. 2021;18(10):1247–1252. doi:10.1038/s41592-021-01261-w

14. Perutz MF. Stereochemistry of cooperative effects in haemoglobin: haem–haem interaction and the problem of allostery. Nature. 1970;228(5273):726-734.

15. Bowman GR, Bolin ER, Hart KM, Maguire BC, Marqusee S. Discovery of multiple hidden allosteric sites by combining Markov state models and experiments. Proc Natl Acad Sci U S A. 2015;112(9):2734–2739. doi:10.1073/pnas.1417811112

16. Kern D, Zuiderweg ERP. The role of dynamics in allosteric regulation. Curr Opin Struct Biol. 2003;13(6):748–757. doi:10.1016/j.sbi.2003.10.008

17. Goodsell DS, Olson AJ. Structural symmetry and protein function. Annu Rev Biophys Biomol Struct. 2000;29(1):105–153.

18. Wang J, Stieglitz KA, Cardia JP, Kantrowitz ER. Structural basis for ordered substrate binding and cooperativity in aspartate transcarbamoylase. Proc Natl Acad Sci U S A. 2005;102(25):8881–8886. doi:10.1073/pnas.0503742102

19. Robert K. Nakamoto, Joanne A. Baylis Scanlon Mka-S. The Rotary Mechanism of the ATP Synthase. Arch Biochem Biophys. 2008;476(1):43–50. doi:10.1016/j.abb.2008.05.004.The

20. Lin Z, Rye H. GroEL-mediated protein folding: Making the impossible, possible. Crit Rev Biochem Mol Biol. 2006;41(4):211–239. doi:10.1080/10409230600760382

21. Jaffe, Eileen K. LHS. The morpheein model of allostery: Evaluating proteins as potential morpheeins. Methods Mol Biol. 2012;796:217–231. doi:10.1007/978-1-61779-334-9

22. Churchfield LA, Medina-Morales A, Brodin JD, Perez A, Tezcan FA. De Novo Design of an Allosteric Metalloprotein Assembly with Strained Disulfide Bonds. J Am Chem Soc. 2016;138(40):13163–13166. doi:10.1021/jacs.6b08458

23. Rivalta I, Sultan MM, Lee N-S, Manley GA, Loria JP, Batista VS. Allosteric pathways in imidazole glycerol phosphate synthase. Proc Natl Acad Sci. 2012;109(22):E1428–E1436. doi:10.1073/pnas.1120536109

24. Motlagh N. H, Wrabi JO, Li J, Hilser VJ. The ensemble nature of allostery. Nature. 2014;508(7496):331–339. doi:10.1038/nature13001.The

25. Ahmed, Mostafa H., Ghatge, Mohini S., Safo MK. Hemoglobin: Structure, Function and Allostery. Subcell Biochem. 2020;94:345–382. doi:10.1007/978-3-030-41769-7

26. Bai F, Branch RW, Nicolau D V., et al. Conformational spread as a mechanism for cooperativity in the bacterial flagellar switch. Science *(80-)*. 2010;327(5966):685–689. doi:10.1126/science.1182105

27. Siddiq MA, Hochberg GK, Thornton JW. Evolution of protein specificity: insights from ancestral protein reconstruction. Curr Opin Struct Biol. 2017;47:113–122. doi:10.1016/j.sbi.2017.07.003

28. Hsia Y, Mout R, Sheffler W, et al. Design of multi-scale protein complexes by hierarchical building block fusion. Nat Commun. 2021;12(1):1–10. doi:10.1038/s41467-021-22276-z

29. Dauparas J, Anishchenko I, Bennett N, et al. Robust deep learning–based protein sequence design using ProteinMPNN. Science *(80-)*. 2022;378(6615):49–56. doi:10.1126/science.add2187

30. Jumper J, Evans R, Pritzel A, et al. Highly accurate protein structure prediction with AlphaFold. Nature. 2021;596(7873):583-589. doi:10.1038/s41586-021-03819-2

31. Zhu W, Shenoy A, Kundrotas P, Elofsson A. Evaluation of AlphaFold-Multimer prediction on multi-chain protein complexes. Bioinformatics. 2023;39(7). doi:10.1093/bioinformatics/btad424

32. Khmelinskaia A, Bethel NP, Fatehi F, Antanasijevic A, Borst AJ. Local structural flexibility drives oligomorphism in computationally designed protein assemblies. Published online 2023:1–32.

33. Li W, Norris AS, Lichtenthal K, et al. Thermodynamic coupling between neighboring binding sites in homo-oligomeric ligand sensing proteins from mass resolved ligand-dependent population distributions. Protein Sci. 2022;31(10):1–17. doi:10.1002/pro.4424

34. Koshland DE, Némethy G, Filmer D. Comparison of Experimental Binding Data. Theor Model proteins Contain subunits. 1966;5(1):365–385.

35. Milligan G, Smith NJ. Allosteric modulation of heterodimeric G-protein-coupled receptors. Trends Pharmacol Sci. 2007;28(12):615–620. doi:10.1016/j.tips.2007.11.001

36. Chaudhry C, Plested AJR, Schuck P, Mayer ML. Energetics of glutamate receptor ligand binding domain dimer assembly are modulated by allosteric ions. Proc Natl Acad Sci U S A. 2009;106(30):12329–12334. doi:10.1073/pnas.0904175106

37. Kang S, Davidsen K, Gomez-Castillo L, et al. COMBINES-CID: An Efficient Method for de Novo Engineering of Highly Specific Chemically Induced Protein Dimerization Systems. J Am Chem Soc. 2019;141(28). doi:10.1021/jacs.9b03522

38. Dixon AS, Schwinn MK, Hall MP, et al. NanoLuc Complementation Reporter Optimized for Accurate Measurement of Protein Interactions in Cells. ACS Chem Biol. 2016;11(2). doi:10.1021/acschembio.5b00753

39. Sahandi Zangabad P, Karimi M, Mehdizadeh F, et al. Nanocaged platforms: Modification, drug delivery and nanotoxicity. Opening synthetic cages to release the tiger. Nanoscale. 2017;9(4). doi:10.1039/c6nr07315h

40. Yu X, Weng Z, Zhao Z, Xu J, Qi Z, Liu J. Assembly of Protein Cages for Drug Delivery. Pharmaceutics. 2022;14(12). doi:10.3390/pharmaceutics14122609

41. Yang EC, Divine R, Miranda MC, Borst AJ, Sheffler W. Computational design of non-porous, pH-responsive antibody nanoparticles. bioRxiv. 2023;9:1–26. 10.1101/2023.04.17.537263

42. Boyken SE, Benhaim MA, Busch F, et al. De novo design of tunable, pH-driven conformational changes. Science *(80-)*. 2019;364(6442):658–664. doi:10.1126/science.aav7897

43. Divine R, Dang H V, Ueda G, et al. Designed proteins assemble antibodies into modular nanocages. Science *(80-)*. 2021;372(6537):1–22.

44. Bethel NP, Borst AJ, Parmeggiani F, et al. Precisely patterned nanofibres made from extendable protein multiplexes. Nat Chem. Published online 2023. doi:10.1038/s41557-023-01314-x

45. Sheffler W, Yang EC, Dowling Q, et al. Fast and versatile sequence-independent protein docking for nanomaterials design using RPXDock. PLoS Comput Biol. 2023;19(5):1–22. doi:10.1371/journal.pcbi.1010680

46. Praetorius F, Leung PJY, Tessmer MH, et al. Design of stimulus-responsive two-state hinge proteins. bioRxiv. Published online 2023:2023.01.27.525968. https://www.biorxiv.org/content/10.1101/2023.01.27.525968v1%0Ahttps://www.biorxiv.org/content/10.1101/2023.01.27.525968v1.abstract

47. Ben-Sasson AJ, Watson JL, Sheffler W, et al. Design of biologically active binary protein 2D materials. Nature. 2021;589(7842). doi:10.1038/s41586-020-03120-8

48. Shen H, Fallas JA, Lynch E, et al. De novo design of self-assembling helical protein filaments. Science *(80-)*. 2018;362(6415). doi:10.1126/science.aau3775

49. Li, Zhe, Shunzhi Wang, Una Nattermann, Asim K. Bera, Andrew J. Borst, Muammer Y., Yaman MJB, et al. “. Accurate computational design of three-dimensional protein crystals. Nat Mater. 2023;1(8).

50. Hsia Y, Bale JB, Gonen S, et al. Design of a hyperstable 60-subunit protein icosahedron. Nature. 2016;535(7610). doi:10.1038/nature18010

51. Caliendo F, Dukhinova M, Siciliano V. Engineered cell-based therapeutics: Synthetic biology meets immunology. Front Bioeng Biotechnol. 2019;7(MAR). doi:10.3389/fbioe.2019.00043

52. Hancock WO, Howard J. Kinesin’s processivity results from mechanical and chemical coordination between the ATP hydrolysis cycles of the two motor domains. Proc Natl Acad Sci U S A. 1999;96(23). doi:10.1073/pnas.96.23.13147

53. Bennett NR, Coventry B, Goreshnik I, et al. Improving de novo protein binder design with deep learning. Nat Commun. 2023;14(1):1–9. doi:10.1038/s41467-023-38328-5

54. Mastronarde DN. SerialEM: A program for automated tilt series acquisition on Tecnai microscopes using prediction of specimen position. In: Microscopy and Microanalysis. Vol 9. ; 2003. doi:10.1017/s1431927603445911

55. Sun M, Azumaya CM, Tse E, et al. Practical considerations for using K3 cameras in CDS mode for high-resolution and high-throughput single particle cryo-EM. J Struct Biol. 2021;213(3). doi:10.1016/j.jsb.2021.107745

56. Punjani A, Rubinstein JL, Fleet DJ, Brubaker MA. CryoSPARC: Algorithms for rapid unsupervised cryo-EM structure determination. Nat Methods. 2017;14(3). doi:10.1038/nmeth.4169

57. Sanchez-Garcia R, Gomez-Blanco J, Cuervo A, Carazo JM, Sorzano COS, Vargas J. DeepEMhancer: a deep learning solution for cryo-EM volume post-processing. Commun Biol. 2021;4(1). doi:10.1038/s42003-021-02399-1

58. The PyMOL Molecular Graphics System, Version 1.2r3pre, Schrödinger, LLC. Title.

59. Pettersen EF, Goddard TD, Huang CC, et al. UCSF ChimeraX: Structure visualization for researchers, educators, and developers. Protein Sci. 2021;30(1). doi:10.1002/pro.3943

60. Kidmose RT, Juhl J, Nissen P, Boesen T, Karlsen JL, Pedersen BP. Namdinator – Automatic molecular dynamics flexible fitting of structural models into cryo-EM and crystallography experimental maps. IUCrJ. 2019;6:526–531. doi:10.1107/S2052252519007619

61. Croll TI. ISOLDE: A physically realistic environment for model building into low-resolution electron-density maps. Acta Crystallogr Sect D Struct Biol. 2018;74(6). doi:10.1107/S2059798318002425

62. Emsley P, Cowtan K. Coot: Model-building tools for molecular graphics. Acta Crystallogr Sect D Biol Crystallogr. 2004;60(12 I). doi:10.1107/S0907444904019158

63. Emsley P, Lohkamp B, Scott WG, Cowtan K. Features and development of Coot. Acta Crystallogr Sect D Biol Crystallogr. 2010;66(4). doi:10.1107/S0907444910007493

64. Liebschner D, Afonine P V., Baker ML, et al. Macromolecular structure determination using X-rays, neutrons and electrons: Recent developments in Phenix. Acta Crystallogr Sect D Struct Biol. 2019;75. doi:10.1107/S2059798319011471

65. Williams CJ, Headd JJ, Moriarty NW, et al. MolProbity: More and better reference data for improved all-atom structure validation. Protein Sci. 2018;27(1). doi:10.1002/pro.3330

